# Holobiont Evolution: Population Theory for the Hologenome

**DOI:** 10.1101/2020.04.10.036350

**Authors:** Joan Roughgarden

## Abstract

This article develops mathematical theory for the population dynamics of microbiomes with their hosts and for holobiont evolution caused by holobiont selection. The objective is to account for the formation of microbiome-host integration.

Microbial population-dynamic parameters must mesh with the host’s for coexistence.

A horizontally transmitted microbiome is a genetic system with “collective inheritance”. The microbial source pool in the environment corresponds to the gamete pool for nuclear genes. Poisson sampling of the microbial source pool corresponds to binomial sampling of the gamete pool. However, holobiont selection on the microbiome does not lead to a counterpart of the Hardy-Weinberg Law nor to directional selection that always fixes microbial genes conferring the highest holobiont fitness.

A microbe might strike an optimal fitness balance between lowering its within-host fitness while increasing holobiont fitness. Such microbes are replaced by otherwise identical microbes that contribute nothing to holobiont fitness. This replacement can be reversed by hosts that initiate immune responses to non-helpful microbes. This discrimination leads to microbial species sorting. Host-orchestrated species sorting (HOSS) followed by microbial competition, rather than co-evolution or multi-level selection, is predicted to be the cause of microbiome-host integration.

## 1 Introduction

The combination of a host with its microbiome has been termed a “holobiont” (Margulis 1991). A “microbiome” is an “ecological community of commensal, symbiotic, and pathogenic microorganisms” that shares the “body space” of a host (Lederberg and McCray 2001). The union of the host nuclear genes with the genes from all the microbes in its microbiome is a “hologenome” (Zilber-Rosenberg and Rosenberg 2008). The configuration of the hologenome in a particular holobiont is a “hologenotype” (Roughgarden 2020). “Holobiont selection” is natural selection on holobionts wherein a holobiont’s survival and/or fecundity depends on the phenotype brought about by its hologenotype.

Studies reveal integration between microbiomes and their hosts extending to development, metabolism, physiology and behavior (Gilbert *et al*. 2012, McFall-Ngai *et al*. 2013, Bordenstein and Theis 2015, Theis *et al*. 2016). New peer-reviewed papers documenting microbiome-host integration appear every day for groups ranging from sponges, corals, various other marine invertebrates, as well as terrestrial invertebrates and vertebrates (Google watch set for “holobiont”). Moreover, experimental studies of joint microbiome-host dynamics and evolution are increasingly appearing (*eg*. Burns *et al*. 2017, Wang *et al*. 2020, 2021).

The purpose of this paper is two-fold. The first is to understand the dynamics of coupled microbe and host populations. What conditions permit the coexistence of microbes within their hosts? The second is to determine whether a microbiome can form an extended genetic system that contributes to holobiont evolution. And in particular, a major task is to account for the formation of microbiome-host integration.

## 2 Preliminary Concepts

A holobiont contains two genetic systems that together comprise the hologenome—the microbial genes and the nuclear genes. Most of this paper focuses on the microbial genes because this component of holobiont inheritance is conceptually understudied relative to nuclear inheritance. The nuclear component is also considered toward the end of the paper in the section on microbiome-host integration.

### 2.1 Mode of Transmission

The inheritance of the microbiome is tied to the mode of microbial transmission—horizontal or vertical. Vertical transmission of microbes invites analogy with the transmission of nuclear genes. It is facilitated by mechanisms such as the acquisition of a maternal microbiome by embryos as they pass through the birth canal and the acquisition of a parental microbiome by chicks incubating in a nest. A many such mechanisms have been described during the last two decades (Rosenberg and Zilber-Rosenberg 2018.)

In many groups the phylogeny of the hosts parallels the composition of their microbiomes, a phenomenon called “phylosymbiosis”—this is taken as evidence of host specificity (*e*.*g*. Moeller *et al*. 2016, Mallott and Amato 2021). However, as Lim and Bordenstein (2020) note, “Phylosymbiosis does not necessarily imply vertical transmission.”

Especially in marine environments, horizontal transmission appears to be the generic mode of microbe transmission, with vertical transmission being a special case. For example, in a few species of corals the zooxanthellae are transmitted vertically in the coral’s gametes (Hirose and Hidaka 2006) and in a few brooding species, planulae larvae are directly released from the coral polyps, possibly also allowing for vertical transmission of zooxanthellae and other microbes acquired during the brooding (Atoda 1947, Harrison and Wallace 1990, Prasetia *et al*. 2017). But the vast majority of zooxanthellae are acquired from the open sea water surrounding the coral (Babcock *et al*. 1986, Trench 1993).

Even terrestrial groups may offer less vertical transmission than might be expected. In a live-bearing cockroach, researchers found only one component of the microbiome to be transmitted vertically, the rest horizontally (Jennings 2019). Another study with cockroaches found that the microbiome in early development was acquired vertically from the parent on egg casings but the microbiome thereafter acquired its microbes horizontally (Renelies-Hamilton *et al*. 2021). Experiments with new-born, microbiota-free honeybees showed that horizontal transmission from hive materials and social contact with nest-mates established a typical gut composition (Powell *et al*. 2014) whereas vertical transmission from queen to workers takes place in bumblebees (Su *et al*. 2021). Thus in terrestrial environments, although vertical transmission occurs to some extent in some groups, horizontal transmission is widespread too.

At this time, many models have been developed using an assumption of vertical transmission or combined vertical/horizontal transmission (Lipsitch *et al*. 1996, Fitzpatrick 2014, Shapiro and Turner 2014, Foster *et al*. 2017, Hurst 2017, Roughgarden 2017, Zeng *et al*. 2017, Osmanovic *et al*. 2018, Roughgarden *et al*. 2018, van Vilet and Doebeli 2019, Roughgarden 2020, Bruijning *et al*. 2021, Obeng *et al*. 2021, Xiong *et. al*. 2022.) Generally speaking, the models tend to argue that vertical transmission promotes, and horizontal transmission degrades, the effectiveness of holobiont selection. Still, Roughgarden (2020) showed that holobiont selection can be effective even with solely horizontal transmission. So, this paper differs from the preceding models in emphasizing horizontal transmission. Vertical transmission can be added in future research as situations require.

### 2.2 Collective Inheritance

Horizontal transmission brings about a kind of inheritance of its own, “collective inheritance,” as distinct from the “lineal inheritance” of a vertically transmitted microbiome, as detailed in the left panel of Figure 1. Collective inheritance is perfectly Darwinian because it supports evolutionary descent through modification, just as lineal inheritance does. However, collective inheritance is not consistent with Neo-Darwinism which relies on Mendelian inheritance (Wright 1931).

**Figure 1:**
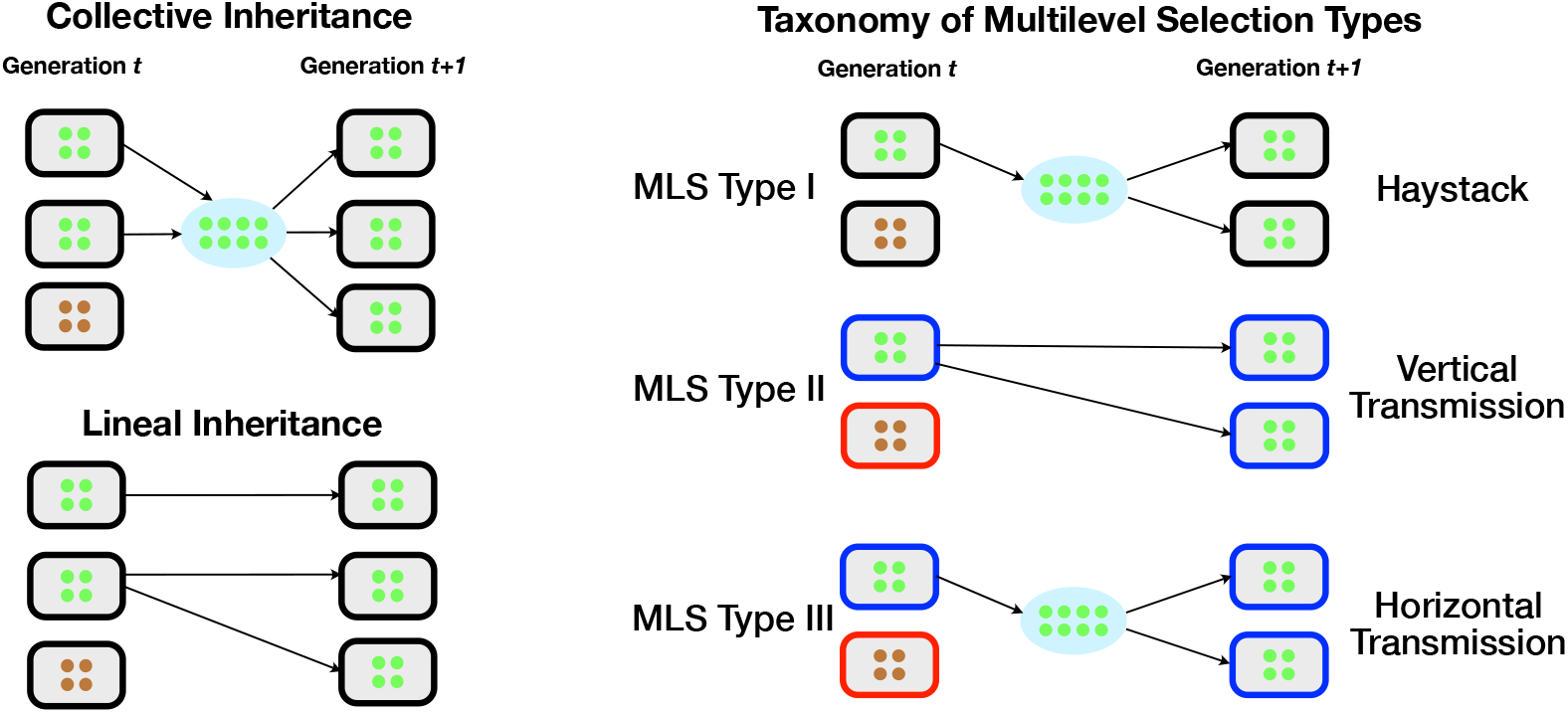
The left panel, top, illustrates collective inheritance as occurs with horizontal microbe transmission and left panel, bottom, illustrates lineal inheritance as occurs with vertical microbe transmission. In both cases the green microbes are selected and the brown ones are not. With collective inheritance the selected parents contribute their microbes to a collective from which the microbes are distributed into the larval hosts. With lineal inheritance the selected parents contribute microbes directly to their own offspring larvae. Either way, the next generation consists of larval hosts whose microbiomes reflect the selection process from the preceding generation. The right panel presents a taxonomy of multilevel selection types. In multilevel selection 1 (MLS1), the environment provides locations indicated as black rectangles with rounded corners. In these locations the organisms undergo natural selection favoring selfish genes, in brown. After selection, offspring are released into a common pool that are redistributed to the locations for the next generation. Locations with many altruistic genes, in green, deliver more offspring into the common pool than locations dominated by selfish genes. If the between-location productivity advantage of altruistic genes exceeds their within-location disadvantage to selfish genes, then altruistic genes evolve overall. Thus, the organisms evolve but the locations themselves do not because they are physical features of the environment. (Maynard Smith in 1964 referred to such locations as haystacks.) In multilevel selection 2 (MLS2), small organisms (microbes) are contained in larger organisms (hosts). While in the host, the microbes undergo natural selection favoring brown selfish microbes. However, hosts containing green altruistic microbes, shown as blue rectangles with rounded corners, survive and reproduce with their microbes better than hosts with brown selfish genes, shown as orange rectangles with rounded corners. The pair is inherited as a unit from one generation to the next. Multilevel selection 3 (MLS3) is defined here as a combination of MLS1 and MLS2. MLS3 shares with MLS1 that microbes are released into a common pool from which they are redistributed into hosts for the next generation. MLS3 shares with MLS2 that hosts themselves evolve along with the microbes. Holobiont selection with vertical transmission is somewhat similar to MLS2 and with horizontal transmission to MLS3. However, holobiont selection differs from MLS by including density dependence within the hosts.

Some skepticism concerning the possibility of evolution *via* holobiont selection with horizontally transmitted microbiomes flows from a mistaken assumption that the mode of inheritance must be lineal for evolution by natural selection to proceed. Lewontin (1970) famously wrote that evolution by natural selection requires three conditions: (1) phenotypic variation, (2) differential fitness, and (3) heritable fitness. The third requirement is stated too narrowly. The third requirement should state that the offspring fitness must correlate with the fitness of the *selected* parents. That is, the offspring must resemble the selected parents including, but not limited to, their immediate parents. Further criticism of the Lewontin criteria appears in Papale (2020).

Other skepticism about holobiont selection pertains to whether a microbiome is intact— instead, it may change during the host’s life time through diet and contact with the environment (*c*.*f*. Knowlton and Rohwer 2003, Moran and Sloan 2015). This paper refers to “core” components of the microbiome that are relatively permanent (*eg*. Jorge *et al*. 2020, Unzueta-Martínez, *et al*. 2022). The transient component of the microbiome might be addressed in future research with a modeling approach drawn from island biogeography (MacArthur and Wilson 1963).

### 2.3 Multilevel Selection

Holobiont selection bears some similarity to concepts of multilevel selection. However, it does not accord with a well-known distinction between multilevel selection 1 (MLS1) and multilevel selection 2 (MLS2) as detailed in the right panel of Figure 1. (Mayo and Gilinsky 1987, Damuth and Heisler 1988, Okasha 2006).

Coevolution, as with flowers and their pollinators, differs from holobiont selection. In co-evolution the fitnesses in species depend on the gene frequencies and population sizes of other species at the same organizational level. (Roughgarden 1983, Dieckmann and Law 1996; Carmona *et. al*. 2015). The coupled equations for coevolution differ from coupled equations for a host with its microbes because host and microbes occupy different organizational levels.

This research contribution is organized as a single narrative together with mathematical supplementary material. The supplementary material is self-contained and can be read by itself, except for reference to some figures found only in the narrative. The narrative—which is this article, offers a verbal account of what is shown in the supplementary material.

## 3 Host with One Microbial Strain

This section investigates the population dynamics of a host with a single-species microbiome. It discusses conditions for the coexistence of the host with its microbes. This population-dynamic analysis is first step prior to analyzing the evolution that results when the microbiome consists of two (or more) microbial strains.

### 3.1 Model

The model assumes that larval hosts are colonized at random by microbes according to a Poisson distribution (Ellis and M. Delbrück 1939). The colonization takes place from environmental source pools (Yamashita *et al*. 2014, Nitschke 2015, Amend *et al*. 2022). The assumption of Poisson colonization implies that some larval hosts remain uncolonized by any microbes while others are colonized by one or more microbes. This assumption was previously used in Roughgarden (2020). Any microbes left over in the microbial pool die. (Nitschke 2015 reports that unincorporated *Symbiodinium* live up to seven days.) Also, colonization takes place when hosts are born not during their later life, consistent with the model’s focus on core rather than transient components of the microbiome.

The generation time for the microbes is assumed to be short relative to the generation time of the host. As a result, the microbes come to population-dynamic equilibrium within the host during each host generation. This assumption in turn implies that the initial condition for larval hosts colonized by one or more microbes is erased. That is, the number of microbes in a mature host is either 0, for larval hosts not colonized by any microbes, or *k* for all larval hosts colonized by one or more microbes because the microbes in all the colonized hosts come to the same within-host equilibrium abundance, *k*. The assumption of short microbial to host generation times allows results to be derived mathematically, unlike Roughgarden (2020) that lacked this assumption and relied on computer iteration instead.

The Poisson distribution has one parameter, sometimes called the “Poisson density”, often denoted as *µ*. If *µ* is low, then many larval hosts are left uncolonized whereas if *µ* is large almost all larvae are colonized. The *µ* depends on the ratio of microbes in the microbial source pool to the number of larval hosts in the host source pool. If there are few microbes per larval host, then *µ* is low and few hosts are colonized, and conversely if there are many microbes per host, then many hosts are colonized. The *µ* changes through time as the ratio of microbes to larval hosts change. According to the Poisson distribution, the probability that a host is colonized by one or more hosts is *P*(*t*) = 1 − *e*^−*µ*(*t*)^ and not colonized by any microbes is 1 − *P*(*t*) = *e*^−*µ*(*t*)^.

Let *H*(*t*) be the number of larvae in the host source pool and *G*(*t*) the number of microbes in the microbial source pool at time *t*. (The notation is *H* for “host” and *G* for “guest”.) The Poisson density parameter, *µ*(*t*), is then assumed to equal *d × g*(*t*), where *g*(*t*) is the ratio of microbes to hosts in their source pools, *G*(*t*)/*H*(*t*), and *d* is an important parameter, usually small, between 0 and 1. The earlier model did not use the parameter *d*, which in effect was assumed to equal 1. Here, the *d* assumes considerable importance.

A low *d* indicates a dilute microbial source pool relative to the host source pool, and a relatively high *d* indicates a dense microbial source pool relative to the host source pool. The *d* might arise from physical mixing processes in the water column that contains the microbial and host source pools. For example, a low *d* might arise if the host source pool is concentrated near one spot on the benthos while the microbial source pool is broadly distributed in the water column. The *d* might also be interpreted as a coefficient of transmission such as found in models of disease dynamics in epidemiology. For example, a low *d* might describe a microbe transmitted only through physical contact and a high *d* might describe a microbe transmitted through water droplets in the air. Furthermore, the *d* might be interpreted as a recruitment coefficient such as found in marine-biology models whereby a high *d* indicates a high recruitment rate by microbes to empty hosts. The *d* may also be a host trait indicating the degree to which the host accepts or rejects the microbes, perhaps mediated through the host’s immune system. Accordingly, a low *d* indicates a microbe strain rejected by the host and a high *d* a microbial strain whose colonization is allowed or even promoted by the host. Depending on context, *d* may be called the dilution factor, the transmission parameter, or the colonization parameter.

The life cycle of a holobiont with a microbiome consisting of one horizontally transmitted microbial strain is diagrammed in the left panel of Figure 2. The life cycle begins with *H*(*t*) and *G*(*t*) for the host and microbial source pools. Then the larval hosts are colonized by microbes according to a Poisson distribution that establishes the hologenotypes and their frequencies. After colonization, microbes proliferate within the juvenile hosts and come to an equilibrium abundance. At this point the hologenotypes are {0} and {*k*} indicating holobionts containing 0 and *k* microbes in them respectively. The frequencies of these hologenotypes are *h*_0_(*t*) and *h*_1_(*t*). Next, the holobiont fitnesses depends on the hologenotype. Let *W*_0_ be the holobiont fitness with an empty microbiome and *W*_1_ be the holobiont fitness with a strain-1 microbial abundance of *k*. The *W* is interchangeably referred to as the holobiont fitness or as the host fitness depending on context. Finally, each holobiont contributes larval hosts to the host source pool based on *W*_0_ or *W*_1_ depending on whether the holobiont has microbes or not. That is, each holobiont without any microbes contributes *W*_0_ larvae to the new host source pool and each holobiont with *k* microbes contributes *W*_1_ larvae to the new host source pool. Turning to the microbes, in population genetics an organism’s overall fitness is typically assembled from its components, say for two loci or two alleles at one locus, based on either an additive or multiplicative formula (*eg*. Ridley 2004). Here, the microbe’s multilevel fitness is assumed to be multiplicative based on contributions from both the within-host fitness and between-host fitness according to *w* = *k × W*_1_. (Upper case pertains to hosts and lower case to microbes.) Hence, the holobionts with microbes in them contribute *k × W*_1_ microbes to the microbial source pool. The next go-around of the life cycle begins with the repopulated larval host and microbial source pools.

**Figure 2:**
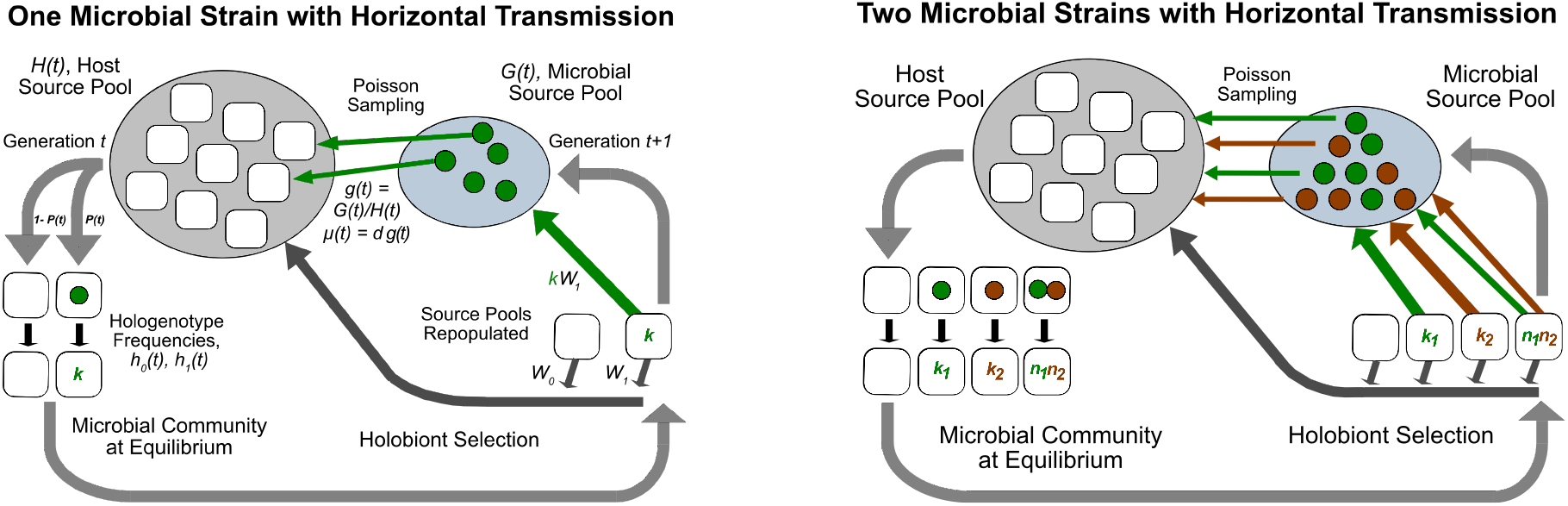
Left: Holobiont life cycle for one microbial strain. Large gray arrows surrounding the diagram indicate progress through a generation starting at t and leading to t + 1. The rectangles with rounded corners in black indicate hosts and the green circles indicate microbes. Right: Holobiont life cycle where microbiome consists of one or two microbial strains (green or brown circle).

The hosts contribute *W*_0_ or *W*_1_ larval hosts to the host source pool depending on their hologenotype. These fitnesses are “single-level” because they depend only on microbe *number* and not microbe *fitness*—the hosts are assumed to be indifferent to the welfare of the microbes in them and sense only their number. Hence, microbe abundance is a trait from the standpoint of the host. But microbe abundance is not a host trait in the usual sense because the host does not contain a gene for the microbe abundance, *per se*, as it does for other traits such as size, color etc. Instead, the microbe abundance is controlled by the *microbe’s* own population dynamics. Still, selection on the host indirectly controls the microbial abundance because of “population-dynamic feedback.” If the microbe is beneficial, then the selection favoring hosts whose microbe strains have a large *k* increases the number of high-*k* microbes in the microbial pool that in turn leads to a higher number of microbes colonizing the larval hosts in the next generation. Conversely, if the microbe is deleterious, then the selection favoring hosts whose microbe strains have a low *k* increases the number of low-*k* microbes in the microbial pool that in turn leads to a lower number of microbes colonizing the larval hosts in the next generation. Because of population-dynamic feedback, microbe abundance effectively becomes a host trait.

In contrast, the microbe fitness *is* “multilevel” because a microbe’s overall success at supplying the microbial pool for the next generation depends somehow on *both* its own fitness and the fitness of the host it resides in. In ecological terminology, the microbes undergo “*K*-selection” within their hosts and “*r*-selection” between hosts, *cf*. MacArthur 1962, Roughgarden 1971).

A physiological interpretation for the product, *k × W*, supposes that a component of *W* refers to host survival. Consider a solitary coral. Compare one polyp whose zooxanthellae strain has a *k*_1_ = 50 and another polyp whose strain has a *k*_2_ = 100. Now, the 100 algal cells of strain-2 supply their coral host with twice as much photosynthate as the 50 cells of strain-1. With the extra photosynthate, twice as many hosts with strain-2 survive desiccation from global warming (*W*_2_ = 2) than hosts with strain-1 (*W*_1_ = 1). Hence, surviving strain-2 hosts release 100 *×* 2 microbes to the pool. But surviving strain-1 hosts release only 50 *×* 1 microbes because strain-1 has a lower *k* and because the death of many strain-1 hosts denies their microbes the chance to reproduce.

In future work the multilevel microbe fitness might be taken as a general function, *w* = *f* (*k, W*_1_), where *f* is an increasing function of both its arguments. In particular, a reviewer has suggested that the multilevel microbe fitness might be taken as *w* = *b × k × W*_1_ where *b* is the number of daughter microbes contributed by each microbe to the microbe source pool. Here, *b* is taken as 1. The reviewer has suggested that the *b* divides out in subsequent analysis, leaving the analysis unaffected. This possibility merits further investigation.

Overall, the model contains only four parameters: *k*, which is the equilibrium number of microbes in the microbe-containing hosts; *d*, which is the colonization parameter; *W*_0_, which is the fitness of holobionts without microbes; and *W*_1_, which is the fitness of holobionts with *k* microbes in each. The model is defined mathematically in Eqs. 4–5 in the Supplementary Material.

### 3.2 Analysis

#### Host with Beneficial Microbes

Figure S2 details conditions for three scenarios for the population dynamics of a beneficial microbe with a host. Scenario-1 is where the microbe’s *k* is above a certain threshold shown in the figure. In this scenario the microbes coexist with the host and both microbes and host populations increase geometrically according to Figure S1. In scenario-1, the microbe can increase when rare and also can coexist with the host.

Scenario-2 is where the microbe’s *k* is between the threshold for scenario-1 and a certain lower threshold, also shown in Figure S2. In scenario-2 the microbe can increase when rare but cannot coexist with the host. In this puzzling situation, the microbes increase when introduced to the host population but do not increase fast enough to keep up with the host’s increase. Hence the *frequency* of hosts containing microbes declines to zero even though both microbe and host populations are increasing in absolute terms. Scenario-2 is illustrated in Figure S3. This scenario is termed “microbial shedding” because the host population effectively sheds its microbes without altogether eliminating them.

In scenario-3, the microbe’s *k* is below the threshold for scenario-2 in Figure S2. In scenario-3 the microbes can neither increase when rare nor coexist with the host and so are completely eliminated, as shown in Figure S4.

The prospect of host productivity exceeding microbe productivity leading to microbe shedding may seem empirically unlikely. Nonetheless, this mathematical possibility resurfaces in the quantitative analysis of further cases. The qualitative insight behind scenario-2 is brought out with an analogy. Image two airplanes, one of which is fueling the other behind it. The airplanes have to match speeds to support a hose connecting them to deliver the fuel. If one plane cannot slowdown enough or the other cannot speed up enough to match speeds, no connection can occur. Indeed, if both planes can fly forward, but the front plane flies faster than the rear plane, then the distance between them gradually increases. Similarly, if one species is to live within another, as microbes do with their hosts, then their speeds of population growth must match.

Microbial shedding occurs here with geometrically growing populations. One might conjecture that microbial shedding relies on the absence of density dependence in the host. However, if host population were density regulated, then some counterpart of the conditions for synchronization of microbe-host population-dynamics would still emerge. If the environmental carrying capacity for the host were *K*, and for the within-host microbes were *k*, some condition involving both *K* and *k* would still be needed for coexistence. In the limit that *K* is large, the situation might converge to that in this paper where *K* → ∞. Furthermore, density dependence cannot be taken as a default assumption as decades of challenges to density-dependent population regulation have documented (Andrewartha and Birch 1954, Sale1977, Strong 1986, Hanski 1990).

#### Host with Deleterious Microbes

If the microbes are deleterious, then a fourth scenario is possible as detailed in Figure S5. Suppose that deleterious microbes are introduced to a holobiont population. Initially, the holobiont population can grow because most holobionts lack any deleterious microbes. But as the microbe population increases, the average growth rate of the holobiont population declines as more holobionts harbor the deleterious microbes. If the microbes increase to the point where the average holobiont fitness drops below 1, then the holobiont population declines. This scenario is one where the microbe and host coexist in the sense that the condition for a microbe-host equilibrium ratio to exist is satisfied, but the equilibrium deleterious microbe to host ratio is large enough to drive the average holobiont fitness below 1. In this situation, the microbe enters the host population, then both microbe and host coexist in a stable ratio, and finally both decline together to extinction. This scenario is illustrated in Figure S6.

No scenarios have been found that involve microbe-host oscillation.

## 4 Host with Two Microbial Strains

This section extends the model of a host with one microbial strain to include two strains. The presence of two strains provides hologenotypic variation among the holobionts upon which holobiont selection can operate.

### 4.1 Model

To set the stage, recall the setup for one locus with two alleles in population genetics. The alleles are *A*_1_ and *A*_2_ with frequencies in the gamete pool of *p*_1_ and (1 − *p*_1_), respectively. The three zygotic genotypes, *A*_1_ *A*_1_, *A*_1_ *A*_2_ and *A*_2_, *A*_2_, are formed from binomial sampling of the gamete pool and occur in Hardy-Weinberg ratios of 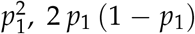 and (1 − *p*_1_)^2^. In keeping with this paper’s terminology, one might say instead that the *A*_1_ and *A*_2_ alleles “colonize” the empty zygotes from the gametic source pool according to binomial sampling. The “copy number” of the *A*_1_ allele is 2 in a *A*_1_ *A*_1_ homozygote and is 1 in a *A*_1_ *A*_2_ heterozygote, and similarly for the copy number of *A*_2_ in its homozygote and the heterozygote.

Here, the microbes of each strain supply genes for the hologenome. The abundance of a microbial strain is the copy number of the gene in that strain. The four hologenotypes are {0,0}, {*k*_1_, 0}, {0, *k*_2_}, and {*n*_1_, *n*_2_} where *k*_1_ and *k*_2_ are the within-host equilibrium abundances of single-strain microbiomes consisting of strain-1 and of strain-2 respectively, and *n*_1_ and *n*_2_ are the within-host equilibrium abundances in a two-strain microbiome obtained from a within-host population-dynamic model.

The ratio of these four hologenotypes after colonization results from independent Poisson sampling of each microbial strain from the microbial source pool. Let the Poisson probability of a host being colonized by one or more microbes of strain-1 as *P*_1_ and for strain-2 as *P*_2_. Accordingly, the probability of empty hosts not being colonized by any microbes of strain-1 is (1 − *P*_1_) and of not being colonized by any microbes of strain-2 is (1 − *P*_2_). Then the Poisson ratios for the four hologenotypes, {0,0}, {*k*_1_, 0}, {0, *k*_2_}, and {*n*_1_, *n*_2_}, are (1 − *P*_1_) (1 − *P*_2_), *P*_1_ (1 − *P*_2_), (1 − *P*_1_) *P*_2_ and *P*_1_ *P*_2_.

The life cycle of a holobiont with a microbiome consisting of two horizontally transmitted microbial strains is diagrammed in the right panel of Figure 2. The life cycle begins with the three state variables for the source pools, *H*(*t*), *G*_1_(*t*) and *G*_2_(*t*) where *H*(*t*) is number of empty hosts in the host pool at time *t*, and *G*_1_(*t*) and *G*_2_(*t*)) are the number of microbes of strain-1 and strain-2 in the microbial pool at time *t*. The total number of microbes in the microbial source pool at time *t, G*(*t*), is *G*_1_(*t*) + *G*_2_(*t*). The ratio of strain-1 microbes to hosts in their source pools, *g*_1_(*t*), is *G*_1_(*t*)/*H*(*t*), and similarly *g*_2_(*t*) is *G*_2_(*t*)/*H*(*t*). The ratio of both microbe strains combined to hosts in their source pools, *g*(*t*), is *G*(*t*)/*H*(*t*). The frequency of strain-1 in the microbial pool at time *t, p*_1_(*t*) is *G*_1_(*t*)/*G*(*t*) and for strain-2, *p*_2_(*t*) is *G*_2_(*t*)/*G*(*t*). The Poisson density parameters for strain-1 at time *t, µ*_1_(*t*), is *d*_1_ *× g*_1_(*t*) and for strain-2, *µ*_2_(*t*) is *d*_2_ *× g*_2_(*t*), where *d*_1_ is the dilution factor for strain-1 and *d*_2_ is the dilution factor for strain-2.

The two strains independently colonize the empty hosts according to a Poisson distribution. Once the juvenile hosts have been initially populated, the microbes proliferate within their hosts, coming to an equilibrium microbiome community. The process of attaining the population-dynamic equilibrium within the hosts erases the initial conditions with which the hosts were colonized. Hence, the equilibrium abundances of single-strain microbiomes consisting of only strain-1 or strain-2 are *k*_1_ and *k*_2_, regardless of the number of colonizing microbes. Similarly, the equilibrium abundances in a dual-strain microbiome consisting of strain-1 and strain-2 are *n*_1_ and *n*_2_ regardless of the number of colonizing microbes. The *n*_1_ and *n*_2_ are the equilibrium within-host microbe population sizes from the Lotka-Volterra (LV) competition equations or other species-interaction model.

Next, the holobiont fitness depend on the hologenotype. Let the holobiont fitness of the hologenotypes, {0, 0}, {*k*_1_, 0}, {0, *k*_2_} and {*n*_1_, *n*_2_} be *W*_0_, *W*_1_, *W*_2_ and *W*_12_, respectively. The holobionts release their empty larval hosts into the host source pool based on these fitnesses.

Turning now to the microbes, the products of within-host microbe population sizes and the holobiont fitnesses are combined into measures of multilevel microbe fitness per holobiont. These measures represent simultaneous success in both within-holobiont *K*-selection and between-holobiont *r*-selection. The multilevel microbe fitnesses per holobiont are: *w*_1,1_ is *k*_1_ *× W*_1_, *w*_1,12_ is *n*_1_ *× W*_12_, *w*_2,12_ is *n*_2_ *× W*_12_ and *w*_2,2_ is *k*_2_ *× W*_2_. These coefficients refer to the multilevel success of a specific microbial strain within a specific microbiome. Thus, *w*_1,1_ is the multilevel fitness per holobiont of a strain-1 microbe in a single-strain microbiome consisting only of strain-1, *w*_1,12_ is the multilevel fitness per holobiont of a strain-1 microbe in a dual-strain microbiome, and so forth. Each single-strain holobiont with *k*_1_ microbes in it contributes *k*_1_ *× W*_1_ microbes of strain-1 to the microbial source pool. Each dual-strain holobiont contributes *n*_1_ *×W*_12_ microbes of strain-1 and *n*_2_ *× W*_12_ microbes of strain-2 to the microbial source pool, and so forth. The next go-around of the life cycle begins with the repopulated larval host and microbial source pools.

Overall, the model now contains ten parameters: *k*_1_ and *k*_2_, which are the equilibrium number of microbes in the single-strain hosts hosts; *n*_1_ and *n*_2_ which are the equilibrium number of microbes of each strain in the two-strain hosts; *d*_1_ and *d*_2_, which are the colonization parameters for each strain; *W*_0_, which is the fitness of holobionts without microbes; *W*_1_, which is the fitness of single-strain holobionts with *k*_1_ strain-1 microbes in each; *W*_2_, which is the fitness of single-strain holobionts with *k*_2_ strain-2 microbes in each; and *W*_12_, which is the fitness of dual-strain holobionts with *n*_1_ strain-1 microbes and *n*_2_ strain-2 microbes in each. If the *n*_1_ and *n*_2_ are obtained from the LV competition equations, the LV reciprocal competition coefficients, denoted as *a*_12_ and *a*_21_, together with *k*_1_ and *k*_2_ are sufficient to determine *n*_1_ and *n*_2_. Mathematically, the model is defined by Eqs. 18, 24, and 25 in the Supplementary Material.

### 4.2 Analysis

The model’s predictions are best viewed in relation to specific situations. It is impractical to give an exhaustive list of all possible outcomes as was done for the host-single strain model of the previous section. The following subsections present some scenarios of holobiont selection on microbial genes that can be contrasted with corresponding scenarios of ordinary natural selection on nuclear genes.

#### No Hardy-Weinberg Analogue

A feature of classical population genetics is the possibility of selective neutrality provided the mating system consists of random union of gametes (or random mating). If the fitnesses for all the genotypes are equal, then any initial allele frequency persists unchanged—the Hardy-Weinberg Law.

The colonization process producing a random assortment of microbe strains in the hosts according to Poisson sampling is analogous to the mating system producing a random assortment of alleles in the nucleus according to binomial sampling. However, the Poisson colonization process is *not* neutral and it produces changes in the frequency of the microbial strains by itself even if the microbial strains have the same fitnesses. There is no analogue of the Hardy-Weinberg Law for the hologenome. Instead, the Poisson colonization process supplies a strong pull to the center. The effect of any selection is combined with this central pull to yield a net result. Figure S10 illustrates what happens to two identical strains that start out with different frequencies. After several generations, both strain frequencies converge to 1/2, as illustrated in Figure S10.

#### Limited Response to Directional Selection

In classical population genetics, directional selection results in fixation of the favored allele and elimination of the alternative allele. Now suppose that the same two alleles from the nucleus are instead found in two strains of the microbiome— microbes are the vehicles delivering genes to the host instead of gametes. If so, is the outcome of directional selection on microbial genes similar to that of directional selection on nuclear genes? Not necessarily. Directional holobiont selection in favor of say, strain-2, does *not* necessarily result in the elimination of strain-1 and fixation of strain-2. Instead, polymorphism may result, as illustrated in Figure S13. The polymorphism reveals the strong pull to the center caused by the colonization dynamics. The holobiont selection in favor of strain-2 does pull the frequency of strain-2 up above the center at 1/2 and pushes the frequency of strain-1 down below the center of 1/2, but nonetheless, strain-1 is not eliminated nor is strain-2 fixed.

The power of the colonization process to override the holobiont selection is controlled by the colonization parameter, *d*. If *d* is low enough, then the directional selection is able to drive the frequency of the inferior strain-1 down to 0 and the frequency of strain-2 up to 1, as illustrated in Figure S14. Furthermore, Figure S15 illustrates the dependence of the equilibrium frequency of strain-1 as a function of *d*, showing that the frequency of strain-1 is zero if *d* is low enough.

The colonization process matters because *d* controls whether the strains are able to express their fitness differences. If *d* is high, then the strains often co-occur in the same host and thus share the same holobiont fitness. Hence, neither can realize an advantage over the other. Conversely, if *d* is low, then the strains often occupy different hosts by themselves and the fitness differences between the strains are expressed so that strain-2 can benefit from its advantage over strain-1. High colonization rates homogenize the hologenotypes across the larval hosts. Conversely, a low colonization rates allow hologenotypic variation to form that is then acted upon by holobiont selection leading to fixation of the selectively favored microbial gene.

This analysis demonstrates that hologenotype frequencies show a limited response to directional holobiont selection, and the extent to which they do respond is qualified by the colonization parameter. This limited response implies that holobiont selection has a limited power to produce holobiont adaptation.

The limited response to holobiont selection has precedence in the classical population genetics of multiple locus genetic systems (Moran 1964, Karlin 1975). There, the evolutionary outcome depends on combining the dynamics of selection with the dynamics of recombination. Here, the evolutionary outcome depends on combining the dynamics of selection with the dynamics of colonization. Evolutionary biology over the years has relied on the canonical one-locus-two-allele setup as a metaphor in which selection completely determines the outcome. It is more realistic, both in classical population genetics and here as well, to regard evolutionary outcomes as resulting from mixing multiple processes, only one of which is natural selection.

The Supplementary Material further explores further situations where the two-strain holobiont is superior or inferior to either of the single-strain holobionts—these correspond to the other standard one-locus-two-allele cases in classical population genetics.

#### Microbial Colonization-Extinction Coexistence

In the previous scenario the microbes did not directly interact with each other. They did affect the host fitness differently, but were otherwise identical just as two alleles in a nucleus might affect host fitness differently but not directly interact with each other. Consider now the converse situation. Suppose the two strains of microbes do interact with each other within the host but each strain has the same effect on the host. From the host’s point of view, the strains are identical, but from the microbes’ point of view, one strain is superior to the other—say, strain-1 out-competes strain-2 whenever the two strains are within the same host. Can strain-2 persist in the holobiont population despite losing in competition to strain-1 whenever both occur together? Yes, the two strains may coexist in a colonization-extinction equilibrium within the holobiont population.

The reason that the strains can coexist despite the competitive asymmetry is that the colonization process provides some empty hosts that end up being colonized only by strain-2, thereby providing a refuge for the competitively inferior strain from the competitively superior strain. However, the production of strain-2 microbes in these refuge hosts must be high enough to compensate for their inability to produce anything in those hosts where strain-1 is also present. Figure S11 illustrates the elimination of strain-2 because its production in hosts where it is by itself does not compensate for the loss of production in hosts where strain-1 is also present, whereas Figure S12 illustrates the coexistence of both strains. This coexistence-scenario is an instance of a meta-population model for patch dynamics featuring a colonization-extinction equilibrium (Levins and Culver 1971, Amarasekare and Possingham 2001).

#### Polymorphism between Altruistic and Selfish Microbes

Can dynamics at the host and microbe levels interact? In the scenario involving directional selection, the fitness differences were solely at the host level. In the scenario involving microbial colonization-extinction coexistence, the fitness differences were solely at the microbe level. Here, the two levels interact.

Suppose again that strain-1 always excludes strain-2 in a host where both are present but now also allow directional holobiont selection in favor of strain-2. That is, strain-2 (the altruistic microbe) sacrifices its competitive ability with respect to strain-1 (the selfish microbe), but receives a higher fitness at the holobiont level. Can holobiont selection favoring strain-2 rescue it from going extinct?

Yes, if the degree of altruism conferred to the host by strain-2 is high enough. Figure S22 illustrates holobiont selection rescuing an altruistic microbe that would otherwise be excluded by the selfish microbe. The Supplementary Material provides the mathematical details.

This section has presented several scenarios for how the microbiome responds to intermicrobial dynamics combined with selection on the host. The model, simple as it is, still involves ten parameters and allows a great many situations to be modeled. Other scenarios can be analyzed with this model without altering the model itself but simply by choosing different parameter values. For example, in all the scenarios presented here the host fitness in the absence of microbes, *W*_0_, has been assumed to equal 1. The possibility that the host requires the microbes to survive might be modeled by assuming *W*_0_ *<* 1. With this assumption, scenarios in which host and microbe coexist depending on sufficient microbe colonization abilitiy can be explored. Undoubtedly empirical situations arise to motivate still other scenarios.

What is remarkable about the hologenome is the difference for the host between the population genetics of its nuclear genes *vs*. its microbial genes. Although nuclear genes can indirectly affect each other by differentially impacting the host phenotype they generally do not directly impact each other even though mechanisms of intragenic conflict such as segregation distortion and meiotic drive do exist (*e*.*g*. Sandler and Golic 1985, Taylor and Ingvarsson 2003, Lindholm *et al*. 2016). In contrast, the host’s microbial genes always directly impact each other and enjoy their own community ecology spanning the host and their source pools in the environment. This, together with the action of holobiont selection, leads overall to remarkably different evolutionary dynamics compared with the classical evolutionary dynamics of one-locus two-allele nuclear genes.

## 5 Microbiome-Host Integration

A common conjecture in discussions of holobiont evolution is that selection on the holobiont as a unit produces a mutualistic interaction between microbiome and host that is adaptive for the holobiont. The remarkable integration between microbiomes and their hosts might be interpreted as a mutualistic coadaptation that results from holobiont selection. This section investigates how microbiome-host integration might form and whether the integration is correctly viewed as a mutualism and whether the integration results from holobiont selection.

### 5.1 Host with Two Nuclear Alleles and Two Microbial Strains

To investigate the formation of holobiont integration one might extend the preceding models to include two vertically transmitted nuclear alleles together with two horizontally transmitted microbial strains. Such a model would describe the coupled population dynamics and genetic changes of both host and microbes jointly.

Figure 3 presents the life cycle for this setup. At the start of each generation the genetically varied larval hosts acquire their microbiomes by independent Poisson sampling of the two strains from the microbial source pool leading to twelve “hologenotypes”. The microbiomes come to a community equilibrium within each host generation that depend on the genetic composition of their hosts. Then holobiont selection occurs. The hosts contribute gametes to a gamete pool. Then larval hosts are assembled with binomial sampling of these gametes (random union of gametes). Meanwhile, the microbiomes contribute microbes to a microbial source pool that are then available for Poisson sampling to begin the next generation.

**Figure 3:**
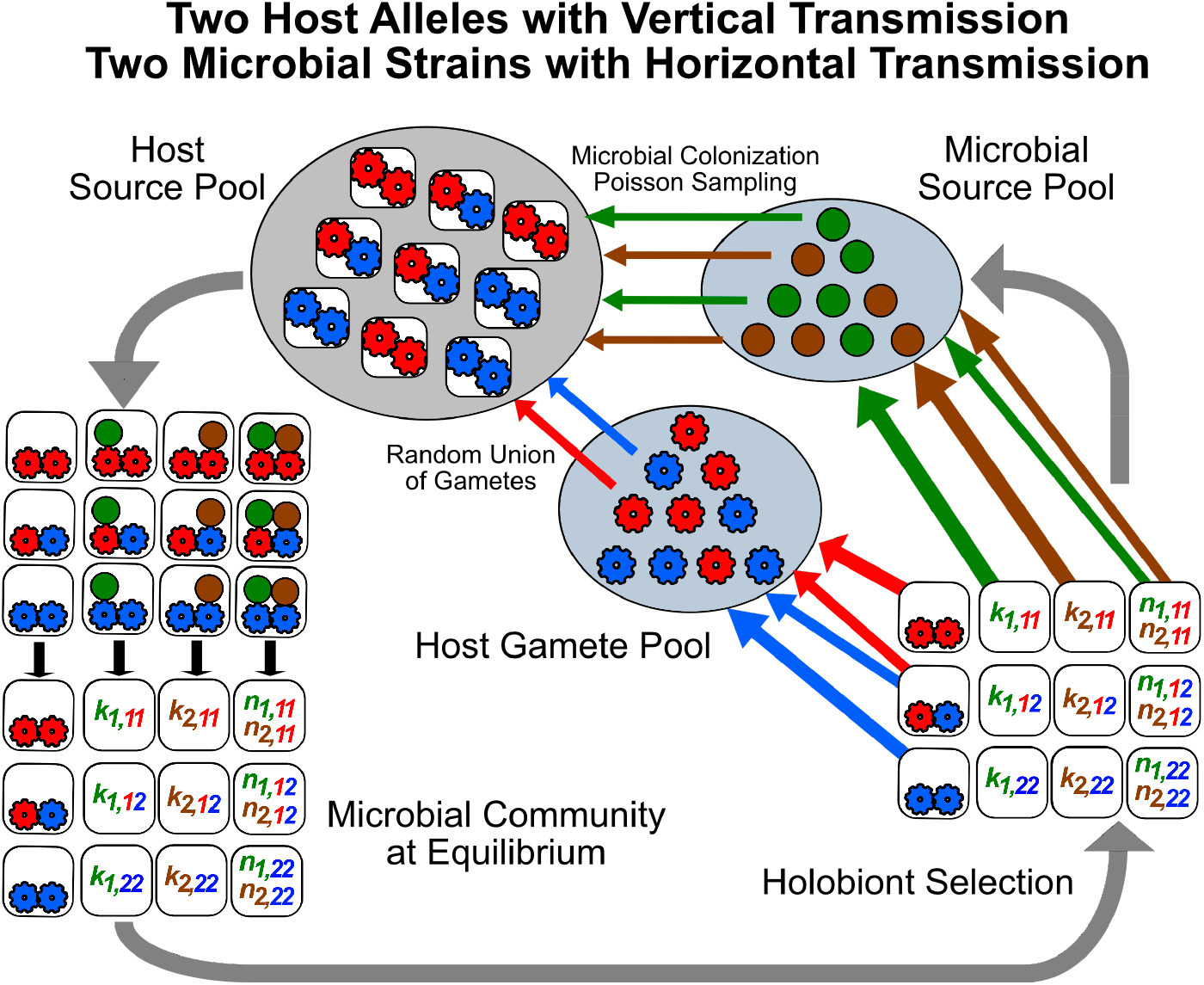
Holobiont life cycle where microbiome consists of one or two microbial strains (green or brown circles) and the host nucleus contains two alleles (red or blue gears).

The equations for this model are a straight-forward extension of the preceding models, but twelve equations are needed for the twelve hologenotypes. The two-microbe model of the preceding section required ten parameters for one genotype of host. With three host genotypes, thirty parameters would be needed for the twelve hologenotypes. Although this setup would potentially allow all conceivable scenarios of microbiome-host interaction to be studied, the model is so large and complicated that the work required for its analysis is not obviously time well spent.

### 5.2 Phenotypic “Effort” Model for Microbiome-Host Integration

Instead, an alternative approach is offered that differs in how the model coefficients are interpreted. In the preceding sections, the microbial and host fitnesses and the colonization coefficients were taken as primitive and simply stipulated for each example. Here, these parameters are interpreted in terms of underlying phenotypic models for the microbes and the host that express what a host and/or a microbe does.

What a host does, or a microbe does, is its “effort”. For example, algae in the microbiome might contribute sugars to the host. The amount of sugar the algae contributes is its “effort”, denoted as *x*, which might be expressed in units of micrograms of sugar contributed per time. This effort lowers the microbe’s within-host fitness, *k*, but increases the host’s fitness, *W*. Similarly, the host might expend effort, *z*, to inhibit the colonization of microbes by manufacturing antibodies, or might promote colonization by producing chemical attractants. The units for this effort might be the amount of ATP needed per time to produce the antibodies or attractants. The net production of this effort considers both the costs of manufacture and the benefits that the antibodies or attractants confer.

Thus, the host fitness is a function of both the microbes’ effort and its own effort, *W*(*x, z*); the microbe’s within-host fitness is a function of its own effort, *k*(*x*); and the microbe’s colonization parameter is a function of the host’s effort, *d*(*z*). The question to be answered is whether holobiont selection adjusts the values of *x* and *z* to promote holobiont adaptation and maximize holobiont fitness.

#### Microbial Altruism *vs*. Selfishness

One might suppose that multilevel selection would lead to an optimal tradeoff by a microbe between its within-host carrying capacity and the between-host fitness of the holobiont. Let the microbe’s effort be called its “altruistic effort”. What is the optimal degree of altruism a microbe should supply the host?

To answer, an optimality criterion is needed. One possibility is the multilevel microbial fitness, *w*. An example of the optimal altruistic effort, 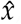, that maximizes *w* appears in Figure S23.

Next, do microbes with the optimal effort exclude all other microbe strains with a non-optimal effort? Suppose a mutant strain arises that is identical in all respects to the optimal strain except for expending a different altruistic effort, 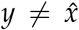. Can the optimal altruistic strain exclude all non-optimal altruistic strains? If it can, then 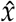 would represent a microbial counterpart to an evolutionarily stable strategy (ESS), which for nuclear genes, is a strategy that cannot be invaded by any other strategy (Maynard Smith 1974).

Indeed, consider the more general question of whether a selfish mutant can invade *any* established altruistic strain, including an optimally altruistic strain. Let the established strain exhibit some degree of altruism, not necessarily optimal, at *x >* 0. Can an otherwise identical but completely selfish mutant with *y* = 0 invade any altruistic strain with *x >* 0?

The Supplementary Material shows that the completely selfish mutant with *y* = 0 *always* increases when rare into a holobiont population fixed for a strain supplying *any* altruistic effort, *x >* 0, including 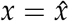. The reason is that a completely selfish mutant has a higher *k* than that of any altruism-providing strain because it does not incur any cost of altruism, assuming all else is equal. Therefore, when the mutant and established strain both colonize a holobiont together, the selfish strain always competitively excludes any established altruistic strain. Hence, the optimal degree of altruism, 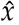, or indeed any altruism at all cannot be the counterpart of an evolutionarily stable strategy because it can always be invaded by a more selfish strategy. Instead, only the completely selfish microbe itself, *x* = 0, is an evolutionarily stable strategy. Figure S24 illustrates a selfish microbe with *y* = 0 excluding an altruistic microbe with 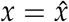.

#### Host Intervention

To obtain an evolutionarily stable altruistic effort by the microbes to the host, the host needs to discriminate against microbial selfishness through its production of antibodies. The quantity of antibodies a host makes results from a balance between the *direct* fitness benefit and cost of its antibody production. The optimal antibody production is a decreasing function of the amount of resources being supplied by the microbes because the more resources being supplied, the less deleterious (or even beneficial) the microbes are. An example of how the optimal antibody production declines as function of the microbe’s contribution to the host appears in Figure S28.

A “side effect” of the host’s antibody production is to lower the selfish microbe’s colonization parameter. It is a side effect because the host determines its antibody production solely from balancing the immediate fitness benefit and cost regardless of how the microbe population dynamics are subsequently affected.

The Supplementary Material shows that there is a minimum amount of altruism, *x*_*min*_(*k*), the microbe must supply for it to increase when rare into the holobiont population. This minimum is a decreasing function of the microbial *k* as shown in Figure S30. The lower the microbe’s *k*, the fewer microbes it is capable of releasing into the microbial source pool. Then these fewer microbes in turn need a higher colonization rate to sustain their population, and the only way to obtain a higher colonization parameter is to increase the amount of altruistic effort it supplies to the host.

However, from the host’s perspective, if the altruism supplied is greater than a certain maximum level, 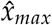, it will shutdown antibody production altogether because the benefit of making antibodies is no longer worth the cost. So, the possibility arises that a microbe’s *k* may be low enough that the altruistic effort it would need to increase when rare exceeds the level at which the host has already shut down antibody production. Increasing altruistic effort beyond 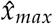 does nothing to improve colonization.

If a microbe cannot make it into the holobiont population when the host is not making any antibodies at all, then it is locked out of the holobiont population. Figure S30 shows 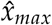 as a horizontal line. All microbes whose carrying capacity lie to the left of the intersection between the curve and the line are locked out from the holobiont population.

If a microbe’s carrying carrying capacity is to the right of the intersection between the curve and line in Figure S30, then the microbe is not locked out. Accordingly, if a microbe’s *k* is large enough that it can enter the holobiont population it must supply at least *x*_*min*_(*k*). However, the microbe also should not bother to supply more than what is necessary to shutdown the host’s antibody production, 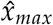. So, a microbe whose within-host carrying capacity is *k* needs to supply some level of altruistic effort between *x*_*min*_(*k*) and 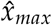.

Well, which is it? Should the microbe supply the minimum altruism needed to enter the holobiont population, *x*_*min*_(*k*), or enough altruism to shut down the host antibody production, 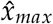? The Supplementary Material shows that the holobionts wind up consisting of microbes that have a bit more altruism than the bare minimum together with hosts that make a corresponding level of antibodies in response. This arguably imperfect outcome, which is worse than the best possible for both parties, nonetheless does represent host-microbiome integration.

#### Host-Orchestrated Species Sorting (HOSS)

The process whereby the host through its antibody production determines which microbes are eligible to colonize represents a kind of species sorting that is here termed as “host-orchestrated species sorting (HOSS).” Species sorting is well known in environmental microbiology. Baas-Becking (1934, *cf*. de Witand Bouvier. 2006) famously wrote “everything is everywhere, but the environment selects.” As Van der Gucht *et. al*. (2007) note, this phrase assumes that high dispersal rates for microorganisms are ubiquitous, an assumption receiving active investigation (Székely and Langenheder 2014, Wu *et. al*. 2018). Here, the idea of species sorting refers to hosts selecting, through their antibody production, which strains in the microbial source pool can colonize.

Metaphorically, the holobiont is a seller’s market for the host. The host sets up the price schedule—for each level of resource the microbes supply there is a corresponding level of antibodies produced, irrespective of any impact on the microbes. In contrast, the microbes must pay to play. They must provide enough resources to obtain a colonization parameter that permits successful colonization. From the microbes’ point of view, the situation is take-it-or-leave-it— provide enough resources to colonize and you’re in, otherwise you’re out. And even after paying the price for successful colonization, a microbial strain faces competition from other strains who may have paid less and still successfully colonized.

The combination of host-orchestrated species sorting followed by competition among the colonizing microbes leads to a conceptual model of holobiont assembly. A diagram illustrating the process of holobiont assembly appears in Figure 4. This holobiont assembly based on host-orchestrated species sorting followed by microbial competition, rather than coevolution or multilevel selection, is the likely cause of host-microbiome integration.

**Figure 4:**
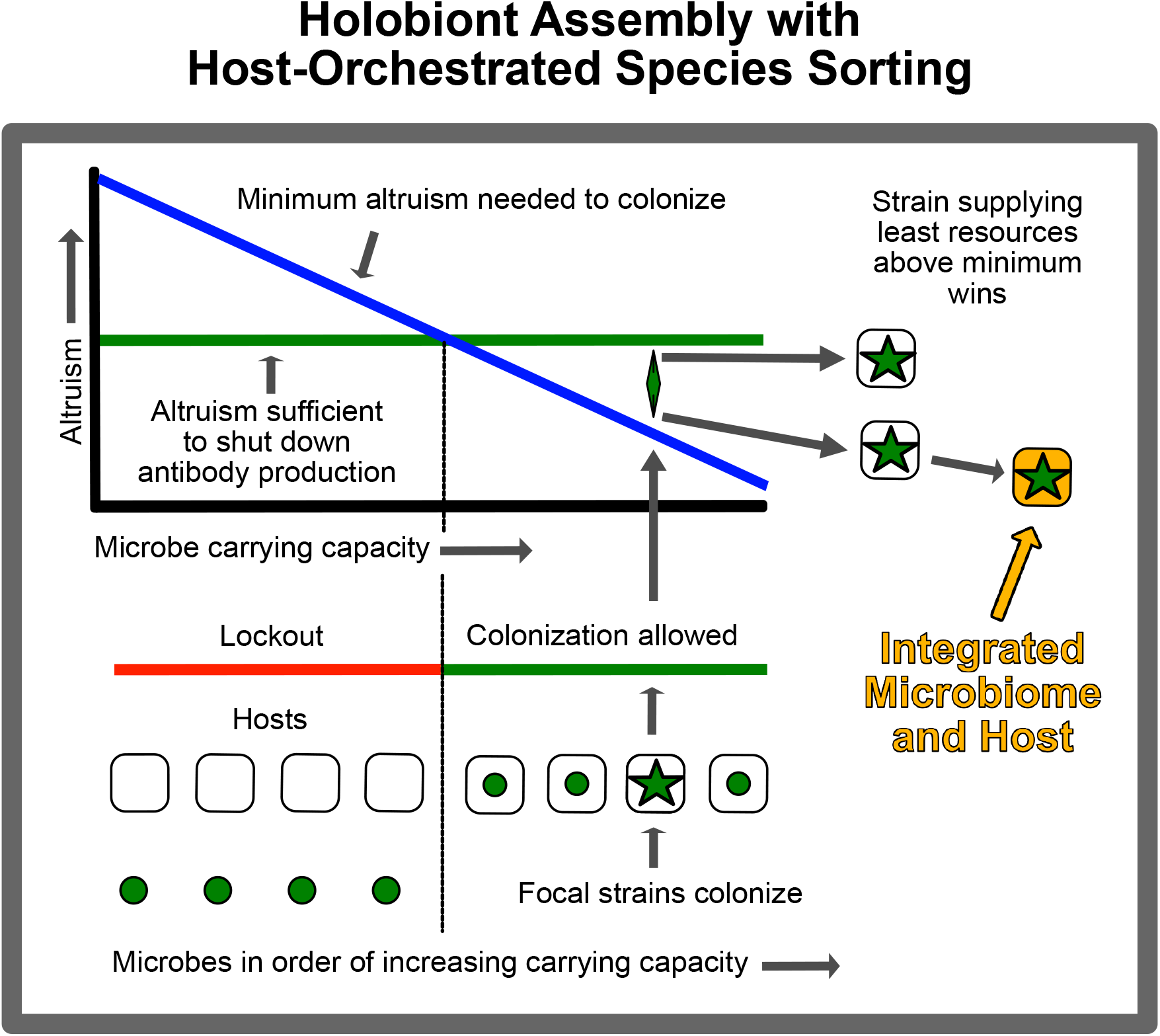
Holobiont assembly with host-orchestrated species sorting (HOSS). At top left, descending blue line indicates minimum altruism microbe must supply to colonize as function of the microbe’s carrying capacity. Horizontal green line indicates amount of altruism that shuts down host antibody production. Microbes required to supply more altruism than needed to shutdown antibody production are locked out. Demarcation between rejected and eligible microbes occurs at intersection of blue and green lines. Suite of microbes differing in carrying capacities depicted at bottom. Microbes with carrying capacities left of demarcation are locked out. Microbes right of demarcation can become incorporated into hosts. Thus, host’s altruism requirement “sorts” available microbes into those that can and cannot colonize. A focal microbe is shown with a star (*). Some strains with this carrying capacity supply the minimum altruism needed to colonize—their altruism level is near the blue line. Other strains supply more than the minimum—their altruism is near the green line. Minimum altruism strain excludes all other strains. Host-orchestrated species sorting followed by competition among strains to supply least altruism consistent with colonization results in holobionts that integrate altruism-supplying microbes with hosts that produce a corresponding level of antibodies.

The theory of HOSS was developed assuming the species sorting is accomplished by the host’s immune system. However, the logic of HOSS remains valid for other mechanisms of host selectivity. The host could employ toxic mucus, chemicals or physical deterrents like spines—any mechanism applied in proportion to the benefit that the microbes supply will suffice. Moreover, the host could facilitate the entry of benefit-producing microbes depending on the amount of benefit supplied. Host facilitation could be represented as a negative antibody, *i*.*e*., a “probody”.

## 6 Discussion

This article has developed mathematical theory for the population dynamics, evolution and assembly of holobionts.

Several population-dynamic outcomes for microbes and hosts have been found, including that a microbe might be excluded from the microbiome, a microbe might initially enter the microbiome only to be shed thereafter from the holobiont, a microbe might join the microbiome and coexist with the host, or a deleterious microbe might enter the microbiome culminating in its own extinction together with the host.

This article has also developed the idea that a horizontally transmitted microbiome constitutes a genetic system with “collective inheritance”. This form of inheritance supports Darwinian descent with modification but is inconsistent with neoDarwinism. With this form of inheritance, the microbial source pool in the environment is the counterpart for the microbiome of the gamete pool for nuclear genes. Poisson sampling of the microbial source pool is the counterpart of binomial sampling of the gamete pool. However, holobiont selection on the microbiome does not lead to a counterpart of the Hardy-Weinberg Law or directional selection that necessarily produces fixation of microbial genes conferring the highest holobiont fitness. The effectiveness of holobiont selection on microbial genes is limited compared to the effectiveness of natural selection on nuclear genes.

The article has further shown that in the absence of holobiont selection, a competitively inferior microbial strain may coexist with a superior strain in a colonization-extinction equilibrium. Alternatively, the presence of holobiont selection permits a polymorphism between altruistic and selfish microbial strains.

The article shows that although a microbe might strike an optimal balance between lowering its within-host fitness while increasing holobiont fitness it is replaced by otherwise identical microbes that contribute nothing to holobiont fitness. This replacement can be reversed by hosts that discriminate against colonization by non-helpful microbes—likely brought about by hosts producing an immune response against non-helpful microbes. This discrimination constitutes microbial species sorting orchestrated by the host immune system.

Holobiont assembly with host-orchestrated species sorting (HOSS) brought about by the host immune system followed by microbial competition extends existing discussion about how the immune system determines biological individuality (Pradeu 2010, 2012, 2016, Gilbert and Tauber 2016). Pradeu (2016) writes “The immune system constitutes a discrimination mechanism, accepting some entities in the organism and rejecting others, thus participating decisively in the delineation of the organism’s boundaries” (p. 803). Gilbert and Tauber (2016) write, the “immune system [is] the mediator of both defensive and assimilative environmental intercourse, where a balance of immune rejection and tolerance governs the complex interactions of the organism’s ecological relationships” (p. 839). This article shows that this discrimination by the immune system results in a composite entity, the holobiont, that possesses a degree of integrated function involving both the microbes and host, even though such integration is generally less than optimal.

This article shows that microbiome-host functional integration is not the result of coevolution because holobiont evolution is a multilevel process whereas coevolution is a single-level process. Moreover, the host-microbiome association is not mutualistic because each party is indifferent to the other’s welfare and neither reciprocally evolves mutual altruism. Furthermore, microbiome-host functional integration does not result from any of the commonly defined versions of multilevel selection. Finally, microbiome-host functional integration is not brought about solely by holobiont selection itself because the effectiveness of holobiont selection on microbial genes is limited. Thus, although a holobiont is a functional unit, it is not an evolutionary unit.

Instead, holobiont formation is a unique and simultaneous combination of both evolutionary and ecological processes. At the upper level, hosts evolve their immune responses by ordinary natural selection on host nuclear genes. At the lower level, microbes colonize hosts in an ecological community-assembly process. The processes at the two levels are coupled because the host’s immune response affects microbial community assembly, sorting the available microbes into those that can and those that cannot enter the holobiont. Competition between microbes that have entered the holobiont reduces the within-host microbial species diversity leaving only those that supply the least possible resources to the host consistent with being able to colonize successfully.

## Supporting information

Mathematical Appendix

## 7 Acknowledgments

The author thanks Seth Bordenstein, Ford Doolittle, Scott Gilbert, Jeremy Van Cleve, Forest Rohwer with members of the BioMath Reading Group at San Diego State University and Priscilla San Juan and Katherine Lagerstrom with members of the Stanford University biology department’s microbiome reading group for helpful comments on the manuscript. The author also thanks Daniel Bolnick and Michael Cortez for superb editing of the manuscript and three anonymous reviewers for extensive and constructive comments. This article is Contribution #1 from a project, The Theory of Holobiont Evolution, funded by the Gordon and Betty Moore Foundation through Grant GBMF10000 to the University of Hawaii.

## References

Amarasekare, P. and H. Possingham. 2001. “Patch Dynamics and Metapopulation Theory: the Case of Successional Species.” J. Theor. Biol. 209:333–344, doi:10.1006/jtbi.2001.2269

Andrewartha H. and L. C. Birch. 1954. The Distribution and Abundance of Animals. University of Chicago Press. Chicago Ill. USA

A. S. Amend, S. O. I. Swift, J. L. Darcy, M. Belcaid, C. E. Nelson, J. Buchanan, N. Cetraro, K. M.S. Fraiola, K. Frank, K. Kajihara, T. G. McDermot, M. McFall-Ngai, M. Medeiros, C. Mora, K. K. Nakayama, N. H. Nguyen, R. L. Rollins, P. Sadowski, W. Sparagon, M. A. Téfit, J. Y. Yew, D. Yogi, and N. A. Hynson. 2022. “A Ridge-to-Reef Ecosystem Microbial Census Reveals Environmental Reservoirs for Animal and Plant Microbiomes”. Proc Nat Acad Sci (USA) 119(33) e2204146119 https://doi.org/10.1073/pnas.2204146119

Atoda K. 1947. “The Larva and Postlarval Development of Some Reef-building Corals. I. Pocillopora damicornis cespitosa (Dana)”. Sci Reports Tohoku Univ. 18: 24–47.

Babcock, R. C., G. D. Bull, P. L. Harrison, A. J. Heyward, J. K. Oliver, C. C. Wallace and B. L. Willis. 1986. “Synchronous Spawnings of 105 Scleractinian Coral Species on the Great Barrier Reef.” Marine Biology 90:379–394.

Baas-Becking L. G. M. 1934. Geologie of Inleidning tot de Milieukunde. W. P. Van Stokum, The Hague, The Netherlands.

Bordenstein, S. R. and K. R. Theis. 2015. “Host Biology in Light of the Microbiome: Ten Principles of Holobionts and Hologenomes.” PLoS Biol. 13(8):e1002226.

Bruijning, M., L. Henry, S. K. G. Forsberg, C. J. E. Metcalf, J. F. Ayroles. 2021. “Natural selection for Imprecise Vertical Transmission in Host-microbiota Systems.” Nature Ecology and Evolution. https://doi.org/10.1038/s41559-021-01593-y

Burns, A., E. Miller, M. Agarwal, A. S. Rolig, K. Milligan-Myhre, S. Seredick, K. Guillemin, and B. J. M. Bohannan. 2017. “Interhost Dispersal Alters Microbiome Assembly and Can Overwhelm Host Innate Immunity in an Experimental Zebrafish Model”. Proc. Nat. Acad. Sci. USA 114(42):11181–11186. http://www.pnas.org/cgi/doi/10.1073/pnas.1702511114

Carmona, D., C. R. Fitzpatrick, and M. T. Johnson. 2015. “Fifty years of Co-evolution and Beyond: Integrating Co-evolution from Molecules to Species.” Mol. Ecol. 24:5315–5329.

Damuth, J. and L. Heisler. 1988. “Alternative Formulations of Multilevel Selection.” Biology and Philosophy 3:407–430.

de Wit, R. and T. Bouvier. 2006. “ ‘Everything is Everywhere, but, the Environment Selects’; What did Baas Becking and Beijerinck Really Say?” Environmental Microbiology 8:755–758. doi:10.1111/j.1462-2920.2006.01017.x

Dieckmann, U., and R. Law. 1996. “The Dynamical Theory of Coevolution: a Derivation from Stochastic Ecological Processes.” J. Math. Biol. 34:579–612.

Ellis, E., and M. Delbrück. 1939. “The Growth of Bacteriophage.” J. General Physiology 22:365– 384. doi: 10.1085/jgp.22.3.365

Fitzpatrick, B. 2014. “Symbiote Transmission and Maintenance of Extra-genomic Associations. Frontiers in Microbiology 5(46)1–15. doi:10.3389/fmicb.2014.00046

Foster, K. R., J. Schluter, K. Coyte, and S. Rakoff-Nahoum. 2017. “The Evolution of the Host Microbiome as an Ecosystem on a Leash”. Nature 548:43–51.

Gilbert, S. F., J. Sapp, and A. I. Tauber. 2012. “A Symbiotic View of Life: We Have Never Been Individuals.” Q. Rev. Biol. 87:325–341.

Gilbert, S. F., and A. I. Tauber. 2016. “Rethinking Individuality: the Dialectics of the Holobiont”. Biol. Philos. 31:839–853, DOI 10.1007/s10539-016-9541-3

Hanski, I. 1990. “Density Dependence, Regulation and Variability in Animal Populations”. Philosophical Transactions: Biological Sciences. 330:141–150

Harrison P. L. and C. C. Wallace 1990. “Reproduction, Dispersal and Recruitment of Scleractinian Corals”. In: Dubinsky Z, editor. Ecosystems of the World. Amsterdam: Elsevier Science; 1990. pp. 133–207.

Hirose, M. and M. Hidaka. 2006. “Early Development of Zooxanthella-containing Eggs of the Corals, Porites cylindirica and Montipora digitata: The Endodermal Localization of Zooxanthellae.” Zoological Science 23:873–881.

Hurst, G. D. D. 2017. “Extended Genomes: Symbiosis and Evolution.” Interface Focus 7: 20170001. http://dx.doi.org/10.1098/rsfs.2017.0001

Jennings, E. C. 2019. “A Holobiont Characterization of Reproduction in a Live-bearing Cockroach, Diploptera punctata.” Phd. diss. Retrieved from https://etd.ohiolink.edu.

Jorge, Fátima, Nolwenn M. Dheilly and Robert Poulin. 2020. “Persistence of a Core Microbiome Through the Ontogeny of a Multi-Host Parasite.” Front. Microbiol. 11:954. doi: 10.3389/fmicb.2020.00954

Karlin, S. 1975. “General Two Locus Selection Models: Some Objectives, Rules and Interpretations.”Theoretical Population Biology 7:364–398.

N. Knowlton and F. Rohwer. 2003. “Multispecies Microbial Mutualisms on Coral Reefs: The Host as a Habitat”. Amer Naturalist 162(Supplement):S51–S62.

Lederberg J, McCray AT. 2001. “Ome Sweet Omics”—a Genealogical Treasury of Words. Scientist. 15:8.

Levins, R. and D. Culver. 1971. “Regional Coexistence of Species and Competition between Rare Species.” Proc. Nat. Acad. Sci. (USA) 68:1246–1248.

Lewontin, R. C. 1970. “The Units of Selection.” Annual Review of Ecology and Systematics, 1:1–18.

Lim, S. J. and S. R. Bordenstein. 2020. “An Introduction to Phylosymbiosis” Proc. R. Soc. B 287: 20192900. http://dx.doi.org/10.1098/rspb.2019.2900.

Lindholm, A. K., K. A. Dyer, R. C. Firman, L. Fishman, W. Forstmeier, L. Holman, H. Johannesson, U. Knief, H. Kokko, A. M. Larracuente, A. Manser, C. Montchamp-Moreau, V.G. Petrosyan, A. Pomiankowski, D. C. Presgraves, L. D. Safronova, A. Sutter, R. L. Unckless, R. L. Verspoor, N. Wedell, G. S. Wilkinson, and T. A. R. Price. 2016. “The Ecology and Evolutionary Dynamics of Meiotic Drive”. Trends in Ecology & Evolution, 31(4) http://dx.doi.org/10.1016/j.tree.2016.02.001

Lipsitch, M. S. Siller, and M. Nowak. 1996. “The Evolution of Virulence in Pathogens with Vertical and Horizontal Transmission.” Evolution 50:1729–1741.

MacArthur, R. H. 1962. “Some Generalized Theorems of Natural Selection”. Proc. Natl. Acad. Sci. USA. 48(11):1893–1897. doi: 10.1073/pnas.48.11.1893

MacArthur R. H. and E. O. Wilson. 1963. “An Equilibrium Theory of Insular Zoogeography.” Evolution 17:373–387.

E. K. Mallott and K. R. Amato. 2021. “Host Specificity of the Gut Microbiome”. Nature Reviews: Microbiology 19:639–653. https://doi.org/10.1038/s41579-021-00562-3

Margulis, L. 1991. “Symbiosis as a Source of Evolutionary Innovation: Speciation and Morphogenesis.” In Symbiogenesis and Symbionticism, edited by L. Margulis and R. Fester, 1–14, Cambridge: MIT Press.

Maynard Smith, J. 1964. “Group Selection and Kin Selection.” Nature 201:1145–1147.

Maynard Smith, J. 1974. “The Theory of Games and the Evolution of Animal Conflicts”. J. theor. Biol. 47(1):209–221 https://doi.org/10.1016/0022-5193(74)90110-6

Mayo, D. and N. Gilinsky. 1987. “Models of Group Selection.” Philosophy of Science 54:515–538.

McFall-Ngai, M., M. G. Hadfield, T. C. Bosch, H. V. Carey, T. Domazet-Loo, A. E. Douglas, N. Dubilier, et al. 2013. “Animals in a Bacterial World, A New Imperative for the Life Sciences.” Proc. Natl. Acad. Sci. (USA) 110:3229–3236.

Moeller, A. H., A. Caro-Quintero, D. Mjungu,A. V. Georgiev, E. V. Lonsdorf, M. N. Muller, A. E. Pusey, M. Peeters, B. H. Hahn, and H. Ochman. 2016. “Cospeciation of gut microbiota with hominids” Science 353(6297):380–382.

Moran, P. A. P. 1964. “On the Non-Existence of Adaptive Topographies.” Annals of Human Genetics 27:383–393.

Moran N. A., and D. B. Sloan. 2015. “The Hologenome Concept: Helpful or Hollow?” PLoS Biol 13(12): e1002311. doi:10.1371/journal.pbio.1002311

Narayana, J. K., M. M. Aogááin, W. W. B. Goh, K. Xia, K. Tsaneva-Atanasova, S. H. Chotirmall. 2021. “Mathematical-based Microbiome Analytics for Clinical Translation.” Computational and Structural Biotechnology Journal. 19:6272–6281.

Nitschke, M. R. 2015. The Free-Living Symbiodinium reservoir and Scleractinian Coral Symbiont Acquisition. PhD. thesis, School of Biological Sciences. The University of Queensland.

Obeng, N., F. Bansept, M. Sieber, A. Traulsen, and H. Schulenburg. 2021. “Evolution of Microbiota-Host Associations: The Microbe’s Perspective”. Trends in Microbiology https://doi.org/10.1016/j.tim.2021.02.005

Okasha, S. 2006. Evolution and the Levels of Selection. Oxford UK: Oxford University Press

Osmanovic, D., D. A. Kessler, Y. Rabin, and Y. Soen. 2018. “Darwinian Selection of Host and Bacteria Supports Emergence of Lamarckian-like Adaptation of the System as a Whole.” Biology Direct. https://doi.org/10.1186/s13062-018-0224-7.

Papale, François. 2020. “Evolution by means of natural selection without reproduction: revamping Lewontin’s account.” Synthese https://doi.org/10.1007/s11229-020-02729-6

Powell J. E., V. G. Martinson, K. Urban-Mead, N. A. Moran. 2014. “Routes of Acquisition of the Gut Microbiota of the Honey Bee Apis mellifera. Appl Environ Microbiol. 80(23):7378–87. https://doi.org/10.1128/AEM.01861-14.

Pradeu, T. 2010. “What is an Organism? an Immunological Answer”. Hist. Philos. Life Sci. 32:247–268.

Pradeu, T. 2012. The Limits of the Self: Immunology and Biological Identity. Oxford University Press, New York.

Pradeu, T. 2016. “Organisms or Biological Individuals? Combining Physiological and Evolutionary Individuality”. Biol. Philos. 31:797–817, DOI 10.1007/s10539-016-9551-1

Prasetia, R., F. Sinniger, K. Hashizume, S. Harii 2017. “Reproductive Biology of the Deep Brooding Coral Seriatopora hystrix: Implications for Shallow Reef Recovery”. PLOS ONE https://doi.org/10.1371/journal.pone.0177034

Renelies-Hamilton, J., K. Germer, D. Sillam-Dussès, K. H. Bodawatta, M. Poulsena. 2021. “Disentangling the Relative Roles of Vertical Transmission, Subsequent Colonizations, and Diet on Cockroach Microbiome Assembly”. mSphere 6:e01023–20. https://doi.org/10.1128/mSphere.01023-20.

Ridley, M. 2004. Evolution. 3rd Edition, Blackwell Pub., Oxford.

Rosenberg, Eugene and Ilana Zilber-Rosenberg. 2018. “The Hologenome Concept of Evolution after 10 years.” Microbiome 6:78 https://doi.org/10.1186/s40168-018-0457-9

Roughgarden, J. 1971. “Density-dependent Natural Selection”. Ecology. 52(3):453–468. https://doi.org/10.2307/1937628

Roughgarden, J. 1983. “The Theory of Coevolution.” In Coevolution, edited by D. J. Futuyma and M. Slatkin, 33–64, Sunderland MA: Sinauer.

Roughgarden, J. 2017. “Model of Holobiont Population Dynamics and Evolution: a Preliminary Sketch.” In Landscapes of Collectivity in the Life Sciences, edited by S. B. Gissis, E. Lamm, and A. Shavit, pp. 325–350. London: The MIT Press.

Roughgarden, J., S. F. Gilbert, E. Rosenberg, I. Zilber-Rosenberg, and E. Lloyd. 2018. “Holobionts as Units of Selection and a Model of Their Population Dynamics and Evolution.” Biological Theory 13:44–65. https://doi.org/10.1007/s13752-017-0287-1.

Roughgarden, J. 2020. “Holobiont Evolution: Mathematical Model with Vertical vs. Horizontal Microbiome Transmission.” Philosophy, Theory and Practice in Biology, 12:2. http://dx.doi.org/10.3998/ptpbio.16039257.0012.002.

Sale, P. 1977. Maintenance of High Diversity in Coral Reef Fish Communities. Amer. Natur. 111:337–359.

Sandler, L. and K. Golic. 1985. “Segregation Distortion in Drosophila”. Trends in Genetics 1:181– 185. https://doi.org/10.1016/0168-9525(85)90074-5

Shapiro, J. and P. E. Turner. 2014. “The Impact of Transmission Mode on the Evolution of Benefits Provided by Microbial Symbionts”. Ecology and Evolution 4: 3350–3361.

Strong, D. 1986. “Density-Vague Population Change”. Trends in Ecology and Evolution. 1(2):39–42.

Su, Q., Q. Wang, X. Mu, H. Chen, Y. Meng, X. Zhang, L. Zheng, X. Hu, Y. Zhai and H Zheng. 2021. “Strain-level Analysis Reveals the Vertical Microbial Transmission During the Life Cycle of Bumblebee”. Microbiome 9:216 https://doi.org/10.1186/s40168-021-01163-1

Székely, A. J. and S. Langenheder. 2014. “The Importance of Species Sorting Differs Between Habitat Generalists and Specialists in Bacterial Communities.” FEMS Microbiol Ecol 87:102–112. DOI: 10.1111/1574-6941.12195

Taylor, D. R. and P. Ingvarsson. 2003. “Common Features of Segregation Distortion in Plants and Animals.” Genetica 117:27–35.

Theis K. R., N. M. Dheilly, J. L. Klassen,R. M. Brucker, J. F. Baines, T. C. G. Bosch, J. F. Cryan, S F. Gilbert, C. J. Goodnight, E. A. Lloyd, J. Sapp, P. Vandenkoornhuyse, I. Zilber-Rosenberg, E. Rosenberg, S. R. Bordenstein. 2016. Getting the Hologenome Concept Right: an Eco-Evolutionary Framework for Hosts and Their Microbiomes. mSystems 1(2):e00028–16. doi: 10.1128/mSystems.00028-16.

Trench, R. K. 1993. “Microalgal-Invertebrate Symbioses: A Review.” Endocytobiosis Cell Res. 9:135–175.

Unzueta-Martínez, A., H. Welch and J. L. Bowen. 2022. “Determining the Composition of Resident and Transient Members of the Oyster Microbiome.” Front. Microbiol. 12:828692. doi: 10.3389/fmicb.2021.828692

van Vilet, S. and M. Doebeli. 2019. “The Role of Multilevel Selection in Microbiome Evolution.” Proc. Nat. Acad. Sci. (USA) http://www.pnas.org/cgi/doi/10.1073/pnas.1909790116.

Van der Gucht, K., K. Cottenie, K. Muylaert, N. Vloemans, S. Cousin, S. Declerck, E. Jeppesen, J-M. Conde-Porcuna, K. Schwenk, G. Zwart, H. Degans, W. Vyverman, and L. De Meester. 2007. “The Power of Species Sorting: Local Factors Drive Bacterial Community Composition Over a Wide Range of Spatial Scales.” Proc. Natl. Acad. Sci. (USA) 104:20404–20409. doi/10.1073/pnas.0707200104

Wang, G. H., B. M. Berdy, O. Velasquez, N. Jovanovic, S. Alkhalifa, K. P. C. Minbiole, and R. M. Brucker. 2020. “Changes in Microbiome Confer Multigenerational Host Resistance after Sub-toxic Pesticide Exposure.” Cell Host & Microbe 27, 213–224 https://doi.org/10.1016/j.chom.2020.01.009

Wang, G. H., J. Dittmer, B. Douglas, L. Huang, R. M. Brucker. 2021. “Coadaptation between Host Genome and Microbiome under Long-term Xenobiotic-induced Selection.” Sci. Adv. 7:eabd4473 pp1–15.

Wright, S. 1931. “Evolution in Mendelian populations.” Genetics 16:97–159. The ISME Journal 12:485–494. doi:10.1038/ismej.2017.183.

Wu, W., HP. Lu, A. Sastri, YC. Yeh, GC. Gong, WC. Chou and CH. Hsieh. 2018. “Contrasting the Relative Importance of Species Sorting and Dispersal Limitation in Shaping Marine Bacterial versus Protist Communities.” The ISME Journal 12:485–494; doi:10.1038/ismej.2017.183

Xiong, X., S. Loo, and M. Tanaka. 2022. “Gut mutualists can persist in host populations despite low fidelity of vertical transmission.” Evolutionary Human Sciences. DOI: 10.1017/ehs.2022.38.

Yamashita, H., G. Suzuki, S. Kai, T. Hayashibara, K. Koike. 2014. “Establishment of Coral–Algal Symbiosis Requires Attraction and Selection” PLoS ONE 9(5): e97003. doi:10.1371/journal.pone.0097003

Zeng, Q., S. Wu, J. Sukumaran, and A. Rodrigo. 2017. “Models of Microbiome Evolution Incorporating Host and Microbial selection.” Microbiome 5(1):127. doi:10.1186/s40168-017-0343-x.

Zilber-Rosenberg, I. and E. Rosenberg. 2008. “Role of Microorganisms in the Evolution of Animals and Plants: the Hologenome Theory of Evolution.” FEMS Microbiology Reviews 32:723– 735. https://doi.org/10.1111/j.1574-6976.2008.00123.x

