## Supplementary material for "Holobiont Evolution: Population Theory for the Hologenome": Mathematical Appendix

**Joan Roughgarden**[1]

**Manuscript as of September 29, 2022**

### Holobiont With One Microbial Strain

#### Population Dynamic Model

This section develops equations for the population dynamics of a holobiont with single-strain microbiome. The equations predict the holobiont population size through time including its host and microbial components together with the conditions for microbe-host coexistence.

The life cycle of a holobiont with a microbiome consisting of one horizontally transmitted microbial strain is diagrammed in the left panel of Figure 2 in the main article. A macro time step begins with the state variables, $H(t)$ and $G(t)$, which are the number of empty hosts in the host source pool and the number of microbes in the microbial source pool at time $t$, respectively. The hologenotypes are $\{0\}$ and $\{k\}$ indicating holobionts containing 0 and $k$ microbes in them respectively. The frequencies of these hologenotypes at time $t$ are $h_0(t)$ and $h_1(t)$, where $h_0(t) + h_1(t) \equiv 1$.

The microbes colonize the empty hosts according to a Poisson distribution with density parameter $\mu(t) \equiv d\,g(t)$, where $g(t) \equiv G(t)/H(t)$ is the ratio of microbes to hosts in their source pools at time, $t$, and $d$ is an important parameter between 0 and 1. A low $d$ indicates a dilute microbial source pool relative to the host source pool, and a high $d$ indicates a dense microbial source pool relative to the host source pool. The $d$ might arise from physical mixing processes in the water column that contains the microbial and host source pools. For example, a low $d$ might arise if the host source pool is concentrated near one spot on the benthos while the microbial source pool is broadly distributed in the water column. The $d$ might also be interpreted as a coefficient of transmission such as found in models of disease dynamics in epidemiology. For example, a low $d$ might describe a microbe transmitted only through physical contact and a high $d$ might describe a microbe transmitted through aerosols. Furthermore, the $d$ might be interpreted as a recruitment coefficient such as found in marine-biology models whereby a high $d$ indicates a high recruitment rate by microbes to empty hosts. The $d$ may also be a host trait indicating the

[1]Hawaii Institute of Marine Biology, University of Hawaii and Department of Biology, Stanford University.

degree to which the host accepts or rejects the microbes. Depending on context, $d$ may be called the dilution factor, the transmission parameter, or the colonization parameter.

According to a Poisson distribution, the fraction of empty hosts that are not colonized by any microbes is $e^{-\mu(t)}$ and the fraction that are colonized by one or more microbes is $(1 - e^{-\mu(t)})$. To simplify notation, put $P(t) \equiv (1 - e^{-\mu(t)})$, which is the probability a host is colonized by one or more microbes, and $(1 - P(t))$ is the probability that a host is not colonized by any microbes.

After the microbiome colonization phase,

$$\begin{aligned} H'(t,0) &= (1 - P(t))\, H(t) \\ H'(t,\bullet) &= P(t)\, H(t) \end{aligned}$$

where $H(t,n)$ is the number of holobionts with $n$ microbes at time $t$. The raised dot ($\bullet$) indicates that one or more microbes are present. Thus, $H'(t,0)$ indicates the number of hosts that were not colonized by any microbes while $H'(t,\bullet)$ indicates the number that were colonized by one or more microbes.

Once the juvenile hosts have been colonized, the microbes proliferate and come to an equilibrium microbiome community structure. Attaining the within-host equilibrium erases the initial conditions with which the hosts were colonized. The equilibrium abundance of the microbes is $k$. Therefore, after the microbiome proliferation phase, the raised dot ($\bullet$) is replaced with $k$, leaving

$$\begin{aligned} H''(t,0) &= (1 - P(t))\, H(t) \\ H''(t,k) &= P(t)\, H(t) \end{aligned}$$

Next, the holobiont fitness depends on the state of the microbiome, its hologenotype, namely, whether the microbe abundance is 0 or $k$. Let $W_0$ be the holobiont fitness with an empty microbiome and $W_1$ be the holobiont fitness with a microbiome abundance of $k$. Now, each holobiont contributes juvenile hosts to the host source pool based on these fitnesses,

$$H(t+1) = W_0\, H''(t,0) + W_1\, H''(t,k)$$

Also, each holobiont contributes microbes to the microbe source pool based on the holobiont fitness *and* on the abundance of microbes within each host,

$$G(t+1) = k\, W_1\, H''(t,k)$$

Substituting for $H''(t,0)$ and $H''(t,k)$ yields

$$\begin{aligned} H(t+1) &= (W_0\, (1 - P(t)) + W_1\, P(t))\, H(t) \\ G(t+1) &= k\, W_1\, P(t)\, H(t) \end{aligned}$$

Now, define the microbe multilevel fitness per holobiont as

$$w_1 \equiv k\, W_1 \tag{1}$$

The microbe multilevel fitness per holobiont includes both its within-holobiont success ($k$) and between-holobiont success ($W_1$). (The microbes undergo $K$-selection within their hosts, and $r$-selection between hosts.) Also, define the net microbe multilevel fitness per holobiont as

$$w(t) = P(t)\, w_1 \tag{2}$$

The $w(t)$ is the microbial multilevel fitness per holobiont taking into account the microbe colonization probability. Finally, define the holobiont mean fitness, $\overline{W}(t)$, as

$$\overline{W}(t) = W_0\,(1 - P(t)) + W_1\, P(t) \tag{3}$$

where the average is respect to the probability distribution of being colonized by zero or by more than zero microbes.

So, substituting the mean holobiont fitness, $W(t)$ and the net microbe multilevel fitness per holobiont, $w(t)$, yields the basic population-dynamic model for the interaction between a single strain of microbes and their host, assuming Poisson microbial colonization:

$$\begin{aligned} H(t+1) &= \overline{W}(t)\, H(t) && (4) \\ G(t+1) &= w(t)\, H(t) && (5) \end{aligned}$$

Furthermore, given $H(t)$ and $G(t)$, the ratio of microbes to hosts in their source pools, $g(t)$, and the proportion of hosts that either remain uncolonized or are colonized during the colonization phase of the life cycle, $h_0(t)$ and $h_1(t)$, are recorded as

$$\begin{aligned} g(t) &= G(t)/H(t) && (6) \\ h_0(t) &= 1 - P(t) && (7) \\ h_1(t) &= P(t) && (8) \end{aligned}$$

These quantities, together with $H(t)$ and $G(t)$ are included in the subsequent figures.

One possible example of a computer iteration with this model yields trajectories illustrated in Figure S1 for a beneficial microbe with $W_1 > W_0 > 1$. In this example, the parameters are arbitrarily set to $d = 0.01$, $W_0 = 5/4$, $W_1 = 2$, and $k = 75$. For this example, the figure shows first, that both $H(t)$ and $G(t)$ asymptotically approach geometric growth with a geometric growth factor each time step, $R_o$, that computes to about 1.373; second, that $g(t)$ asymptotically approaches a constant, $g_o$, indicating that about 17.918 microbes exist in the microbial source pool for each empty host in the host source pool; and third that the hologenotype frequencies, which

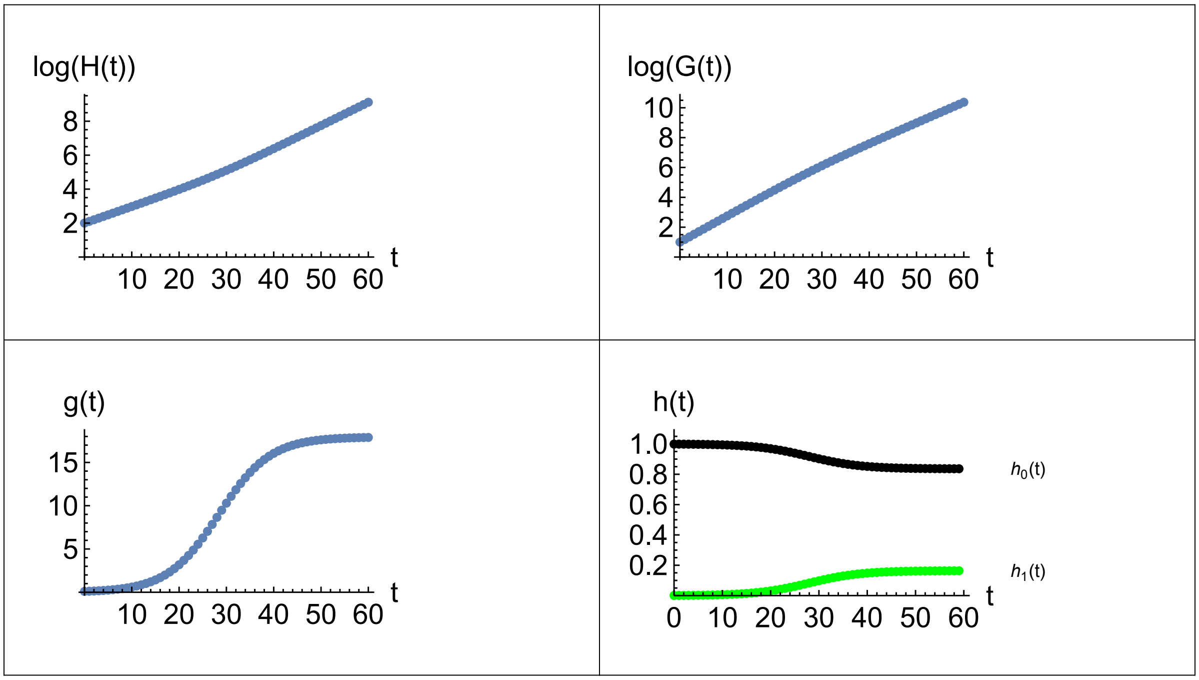


Figure S1: *Trajectories of* $\log_{10}(H(t))$, $\log_{10}(G(t))$ *and* $g(t)$ *through time for a beneficial microbe. Here,* $H(0) = 100$, $G(0) = 10$, $d = 0.01$, $W_0 = 5/4$, $W_1 = 2$, $k = 75$ *and* $t_{\max} = 60$. *Asymptotically,* $R_{oH} \approx 1.373$, $R_{oG} \approx 1.374$, $W_o = 1.373$, $g_o = 17.918$, $h_{0o} = 0.836$ *and* $h_{1o} = 0.164$ *where* $R_{oH} = H(t_{\max})/H(t_{\max}-1)$ *and* $R_{oG} = G(t_{\max})/G(t_{\max}-1)$.

are the frequencies of holobionts with 0 microbes or $k$ microbes, $h_0(t)$ and $h_1(t)$, asymptotically approach 0.836 and 0.164 respectively. The asymptotic Poisson density parameter, $\mu_o = d\, g_o$, is 0.17918. Hence, after colonization, the fraction of hosts that remain empty, $h_{0o} = e^{-\mu_o}$, is 0.836 and the fraction of hosts that was colonized by one or more microbes and that now has $k$ microbes in them, $h_{1o} = 1 - e^{-\mu_o}$, is 0.164, as already indicated. According to the properties of a Poisson distribution, the average number of colonizing microbes per host, averaged over both the empty and non-empty hosts, is $\mu_o$, namely, 0.17918. The average number of colonizing microbes in the non-empty hosts is then $\mu_o/(1-e^{-\mu_o}) = 1.092$.

#### Asymptotic Properties

To find equations for the holobionts' asymptotic geometric growth factor, $R_o$, and the asymptotic ratio of microbes to hosts in their source pools, $g_o$, put $H(t+1) = W_o H(t)$ on the left hand side of Eq. 4, divide out the $H(t)$, substitute for $P_0$ and $P$ explicitly, leaving

$$W_o = W_0\, e^{-\mu_o} + W_1\,(1 - e^{-\mu_o}) \tag{9}$$

Turning now to Eq. 5, put $G(t+1) = W_o\, G(t)$ on the left hand side, divide both sides by $H(t)$, and identify $G(t)/H(t)$ on the left hand side as $g_o$. Then divide both sides by $g_o$, leaving

$$W_o = \frac{k\, W_1\, (1 - e^{-\mu_o})}{g_o} \tag{10}$$

Setting these two equations for $W_o$ equal to each other, and putting $\mu_o = d\, g_o$, yields a single equation for $g_o$

$$W_0\, e^{-d\, g_o} + W_1\, (1 - e^{-d\, g_o}) = \frac{k\, W_1\, (1 - e^{-d\, g_o})}{g_o}$$

which is best rewritten as

$$g_o = \frac{k\, W_1\, (1 - e^{-d\, g_o})}{W_0\, e^{-d\, g_o} + W_1\, (1 - e^{-d\, g_o})} \tag{11}$$

The right hand side (RHS) of Eq. 11 is a concave increasing function of $g_o$. If the slope of the RHS is greater than 1 at $g_o = 0$, then the curve for the RHS initially rises above the line for $g_o$ and then eventually bends down and intersects the line for $g_o$. The value of $g_o$ at the intersection point is the desired answer. The condition for a solution to exist is found from

$$\frac{\partial\, RHS}{\partial\, g_o}\Big|_{g_o=0} = d\, k\, \frac{W_1}{W_0}$$

Therefore, the condition for the host and microbe to coexist is

$$d\, k\, \frac{W_1}{W_0} > 1 \tag{12}$$

To use Eq. 11 with this example, the FindRoot function in *Mathematica* (with a seed of 50) immediately returns the answer as $g_o = 17.938$. Then evaluating either Eq. 9 or Eq. 10 with this value of $g_o$ returns $W_o = 1.373$. These values of $g_o$ and $W_o$ analytically confirm those previously obtained by the numerical iteration of Eqs. 4 and 5 as illustrated in Figure S1.

A special case is where the microbe has no effect on the host, so that $W_0 = W_1 = W$. Then Eq. 11 simplifies to

$$g_o = k\, (1 - e^{-d\, g_o})$$

The root to this transcendental equation, in *Mathematica* notation, is

$$g_o|_{W_0=W_1} = \frac{d\, k + \text{ProductLog}[-d\, k\, e^{-d\, k}]}{d}$$

The $g_o$ can be directly evaluated and plotted in *Mathematica* with this formula based on the model parameters. The $g_o$ is an increasing function of both $d$ and $k$.

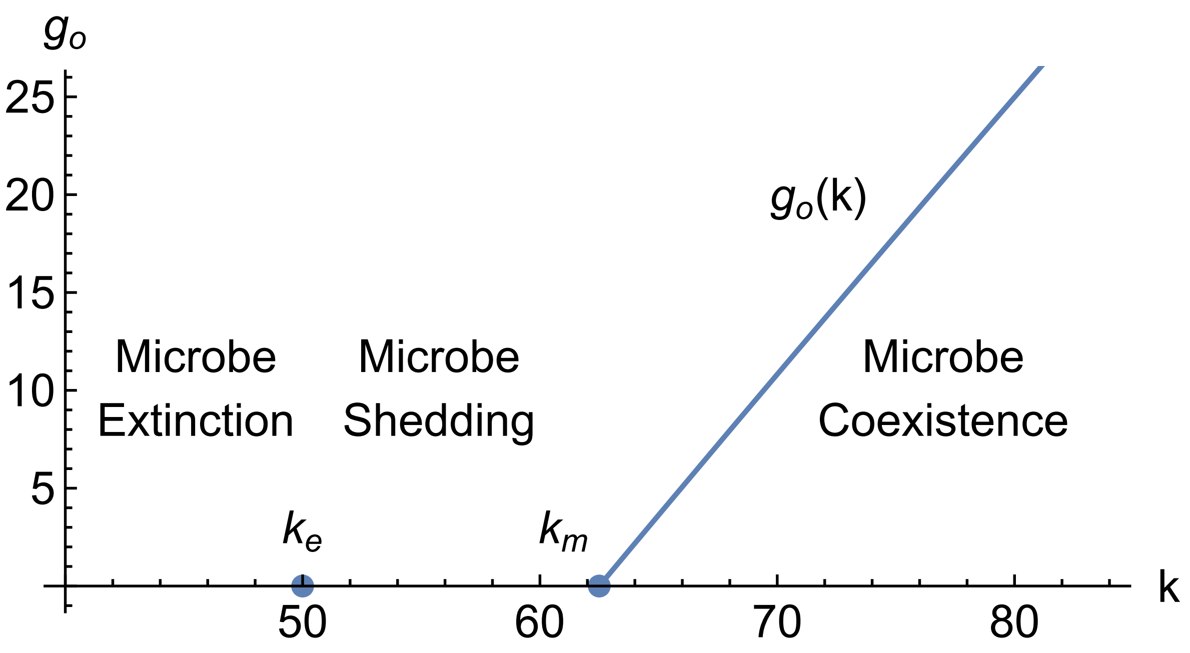


Figure S2: *Beneficial microbe*, $d = 0.01$, $W_0 = 5/4$, $W_1 = 2$.

#### Conditions for Increase When Rare

To analyze the conditions for microbiome-host coexistence, one criterion is the ability of each party to increase when the other is rare. To determine if the microbes can increase when rare upon introduction into a population of uncolonized hosts, rewrite Eq. 5 as

$$\begin{aligned} \Delta G &\equiv G(t+1) - G(t) \\ &= k\, W_1\, (1 - e^{-d\,\frac{G}{H}})\, H - G \end{aligned}$$

The $\Delta G$ is zero at $G = 0$ because no microbes have yet been introduced. To see if $\Delta G$ becomes positive with the addition of some microbes, check that the derivative of $\Delta G$ with respect to $G$ is positive when $G = 0$. So, differentiate $\Delta G$ with respect to $G$ and evaluate at $G = 0$, yielding

$$\frac{\partial\, \Delta G}{\partial\, G}|_{G=0} = d\, k\, W_1 - 1$$

Hence the microbes can enter the holobiont system if

$$d\, k\, W_1 > 1 \tag{13}$$

This requirement means that to enter the uncolonized host population and form holobionts, the microbes must be able to do better than to merely replace themselves taking into account the abundance realized within their hosts, the holobiont fitness and the dilution factor. Eq. 13

illustrates that the microbe's dilution factor, $d$, and its multi-level fitness per holobiont, $w_1$, often appear together indicating an equivalence of sorts between these two parameters. That is, the microbes can enter a population of empty hosts either with a high enough transmissibility, $d$, or a high enough multilevel fitness per holobiont, $w_1$, or both. And for completeness, it is worth noting that a population of empty hosts cannot increase unless $W_0 > 1$.

From Eq. 13, for a given $d$ and $W_1$, the within-host carrying capacity, $k$, must be greater than $k_e$ for microbes to enter the holobiont population, where

$$k > k_e \equiv \frac{1}{d\,W_1} \tag{14}$$

The example in Figure S1 satisfies this condition.

Although Eq. 14 above has just supplied a condition for the microbes to enter the holobiont population, recall that Eq. 12 earlier supplied a condition for the microbe and host to coexist. From Eq. 12, for a given $d$, $W_0$, and $W_1$, the within-host carrying capacity, $k$, must be greater than $k_m$ for microbes and host to coexist, where

$$k > k_m \equiv W_0\,\frac{1}{d\,W_1} = W_0\,k_e \tag{15}$$

The $k_m$ is greater than $k_e$ if $W_0 > 1$, which is automatically true if a population of empty hosts can increase in the absence of microbes. If $k$ is between $k_e$ and $k_m$, microbes can enter the holobiont population but cannot coexist with the host, as now discussed.

#### Coexistence: Host with Beneficial Microbes

Although Figure S1 illustrates an example where a beneficial microbe and host coexist, such coexistence is not inevitable and depends on the relative kinetics of microbial and host population growth.

The function, $g_o(k)$ is found by solving Eq. 11 for various values of $k$. This function is depicted in Figure S2 as a (slightly) curved line running up from the $x$-axis. The curve for $g_o(k)$ hits the $x$-axis at $k = k_m$, which is 62.5 in this example. If $k > k_m$ then the host and beneficial microbes can coexist. This situation was shown previously in Figure S1 with $k = 75$.

If $k$ is in the interval, $(k_e, k_m)$, then the beneficial microbes can enter an empty host population, but their reproductive rate cannot keep up with the reproductive rate of the hosts, leading to the hosts shedding the microbes. That is, both the microbes and host populations increase through time, but the fraction of hosts with microbes in them declines to zero. This situation is shown in Figure S3 with $k = 55$.

Finally, if $k < k_e$, which is 50 in this example, then the beneficial microbes cannot enter the empty host population to begin with. Figure S4 illustrates this case where $k = 35$ and where the microbe source pool is initially seeded with a microbe population size of 10.

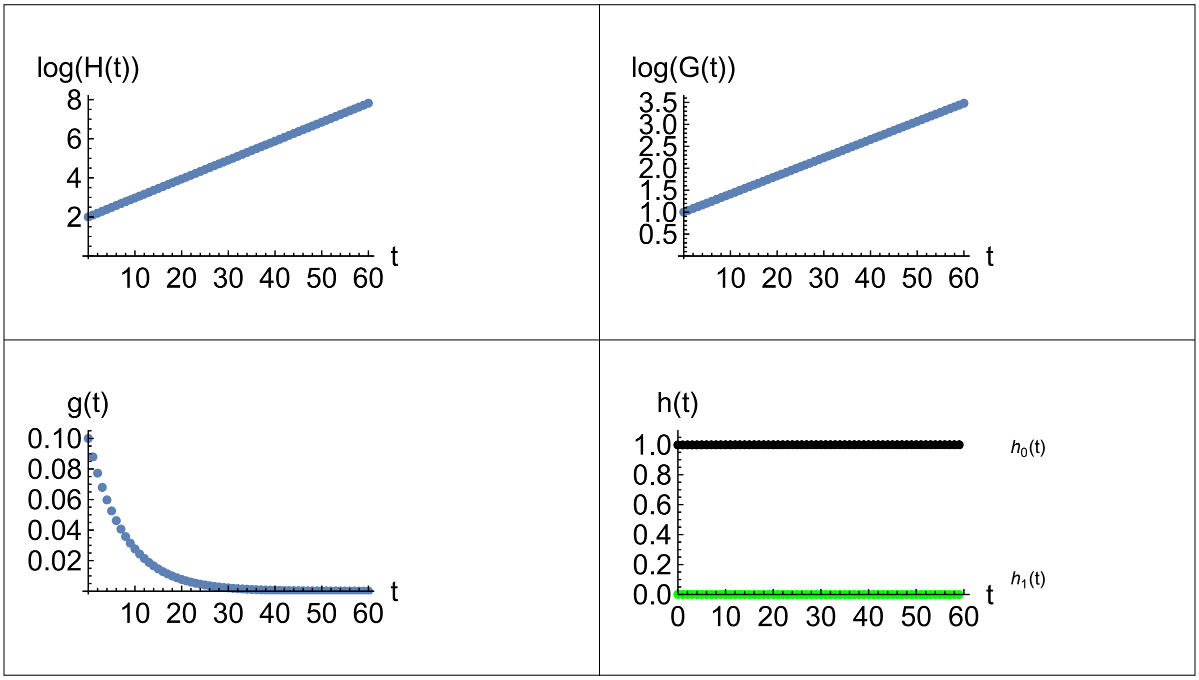


Figure S3: *Trajectories of* $\log_{10}(H(t))$, $\log_{10}(G(t))$ *and* $g(t)$ *through time for a beneficial microbe. Here,* $H(0) = 100$, $G(0) = 10$, $d = 0.01$, $W_0 = 5/4$, $W_1 = 2$, $k = 55$ *and* $t_{\max} = 60$. *Asymptotically,* $R_{oH} \approx 1.25$, $R_{oG} \approx 1.100$, $g_o \to 0$, $h_{0o} \to 1$ *and* $h_{1o} \to 0$ *where* $R_{oH} = H(t_{\max})/H(t_{\max} - 1)$ *and* $R_{oG} = G(t_{\max})/G(t_{\max} - 1)$.

#### Coexistence: Host with Deleterious Microbes.

Turning now to deleterious microbes. The condition for the holobiont to persist with deleterious microbes is found from Eq. 4 which is written as a function of $g_o$ and with an explicit $P$:

$$\overline{W}(g_o) = W_0\, e^{-d\, g_o} + W_1\, (1 - e^{-d\, g_o}) > 1 \tag{16}$$

If there are no microbes, $g_o = 0$, then the holobiont fitness is $W_0$, which is $> 1$ by assumption. As $g_o$ increases above 0, then $\overline{W}$ becomes an average of $W_0$ and $W_1$.

If the microbes are deleterious, then $W_1 < 1 < W_0$. In this case, $\overline{W}$ is a decreasing function of $g_o$. Therefore at some critical value of $g_o$, say $g_{oc}$,

$$\overline{W}(g_{oc}) = W_0\, e^{-d\, g_{oc}} + W_1\, (1 - e^{-d\, g_{oc}}) = 1 \tag{17}$$

The root, $g_{oc}$, to this equation is the critical value of the microbe to host ratio that allows coexistence of the host with a deleterious microbe. If $g_o$ is less than $g_{oc}$ then the host may coexist with the deleterious microbes. But if $g_o$ is greater than $g_{oc}$, then the microbes colonize the hosts and proceed to drive the holobiont population to extinction.

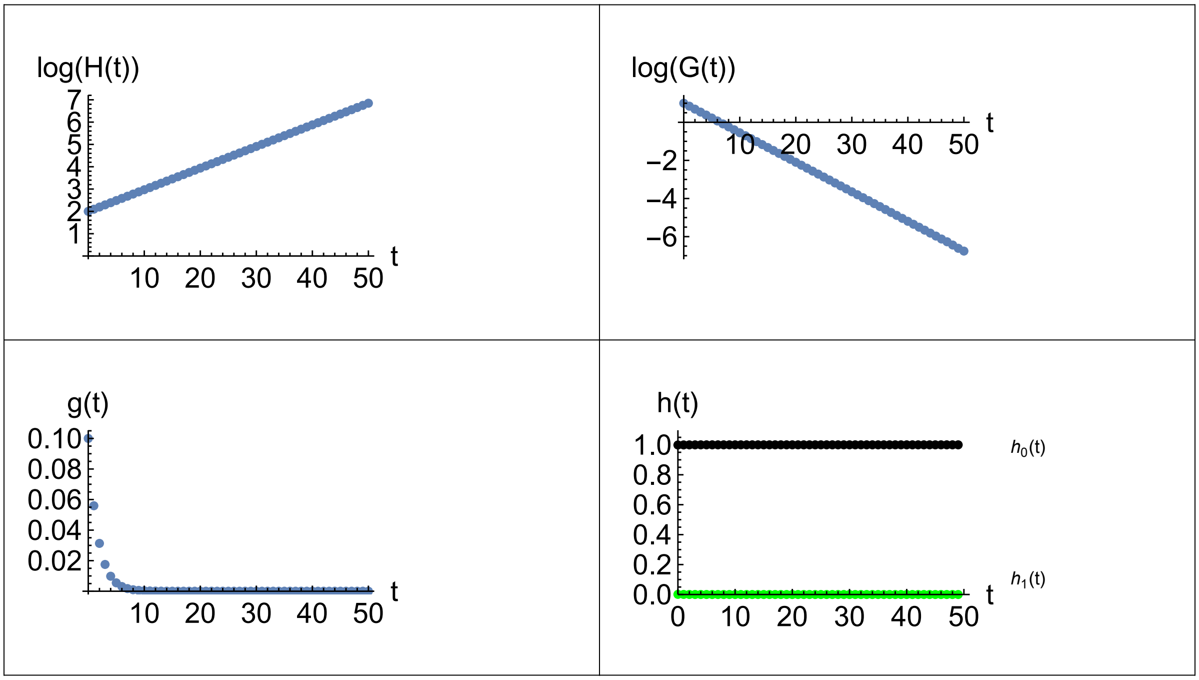


Figure S4: *Trajectories of* $\log_{10}(H(t))$, $\log_{10}(G(t))$ *and* $g(t)$ *through time for a beneficial microbe. Here,* $H(0) = 100$, $G(0) = 10$, $d = 0.01$, $W_0 = 5/4$, $W_1 = 2$, $k = 35$ *and* $t_{\max} = 50$. *Asymptotically,* $R_{oH} \approx 1.25$, $R_{oG} \approx 0.714$, $g_o \to 0$, $h_{0o} \to 1$ *and* $h_{1o} \to 0$ *where* $R_{oH} = H(t_{\max})/H(t_{\max} - 1)$ *and* $R_{oG} = G(t_{\max})/G(t_{\max} - 1)$.

The $g_{oc}$ depends on $d$, $W_0$ and $W_1$, whereas the condition for the microbes to invade when rare depends on $d$, $k$, and $W_1$. So, given $d$, $W_0$, and $W_1$, the parameter $k$ can be freely varied allowing examples to be constructed of holobionts that coexist with deleterious microbes *vs.* holobionts that are driven to extinction by deleterious microbes.

Four possible cases exist with deleterious microbes as diagrammed in Figure S5. For this example, the same parameters are used as in Figure S1 except that here $W_0 > 1 > W_1$, and specifically, $W_0$ remains equal to 5/4, but $W_1$ is now dropped from 2 down to 3/4. With these parameters in Eq. 13, the microbes can enter the population if its $k > k_e = 133.33$. Next, the $g_{oc}$ works out to be 69.315 and is depicted as a horizontal line in the figure. Also, the function, $g_o(k)$ is found by solving Eq. 11 for various values of $k$. This function is depicted in Figure S5 as a curved line running up from the $x$-axis at $k_m$ and intersecting the $g_{oc}$ horizontal line. This intersection point occurs at a critical $k_c$ which works out to equal 184.839 in this example.

If $k > k_c$, then the deleterious microbes can enter a population of empty hosts starting with a $g$ near zero that increases through time until the $g$ exceeds the critical $g_{oc}$ whereupon both the host and microbe populations jointly decline to extinction. This situation is shown in Figure S6 with $k = 200$.

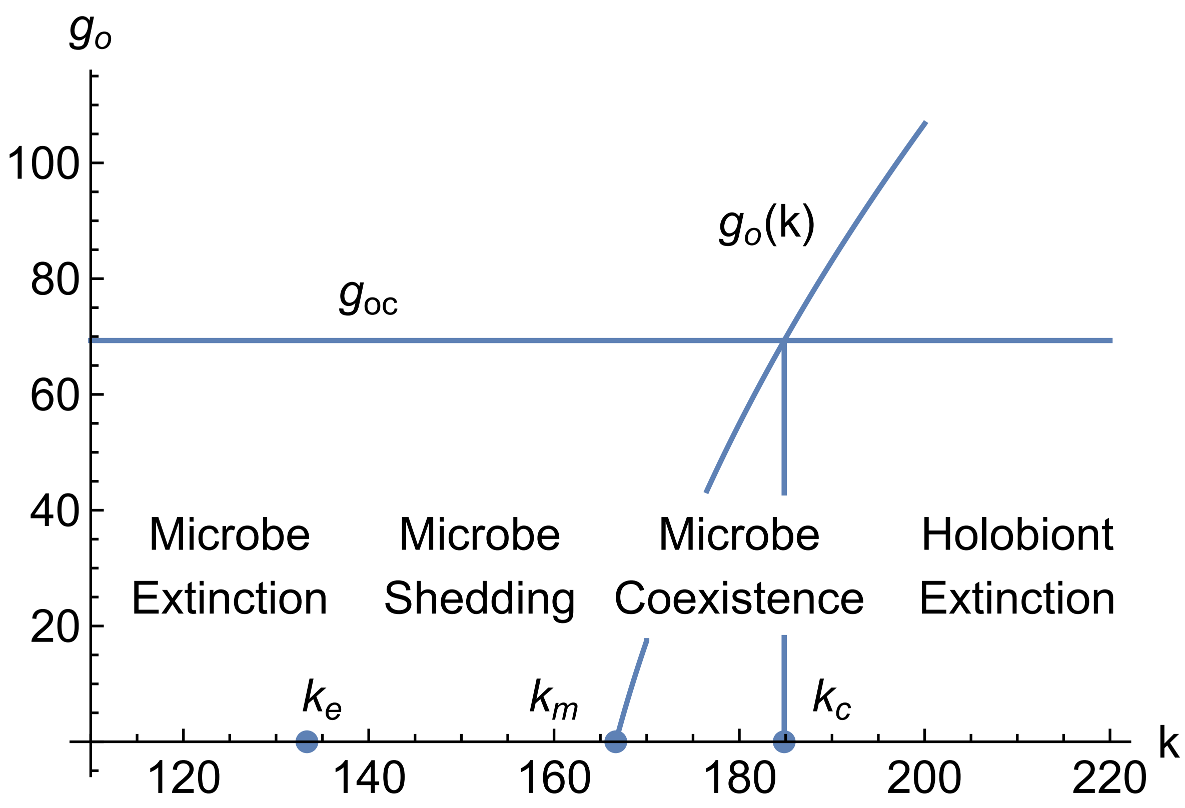


Figure S5: *Deleterious microbe,* $d = 0.01$, $W_0 = 5/4$, $W_1 = 3/4$

171 The curve for $g_o(k)$ hits the $x$-axis at $k = k_m$, which is 166.666. If $k$ is in the interval, $(k_m, k_c)$, then the host and deleterious microbes can coexist as shown in Figure S7 with $k = 175$.

If $k$ is in the interval, $(k_e, k_m)$, then the deleterious microbes can enter an empty host population, but its reproductive rate cannot keep up with the reproductive rate of the hosts, leading to the hosts shedding the microbes as shown in Figure S8 with $k = 150$.

Finally, if $k < k_e$, which is 133.33 in this example, then the deleterious microbes cannot enter the empty host population to begin with. Figure S9 illustrates this case where $k = 125$ and where the microbe source pool is initially seeded with a microbe population size of 10.

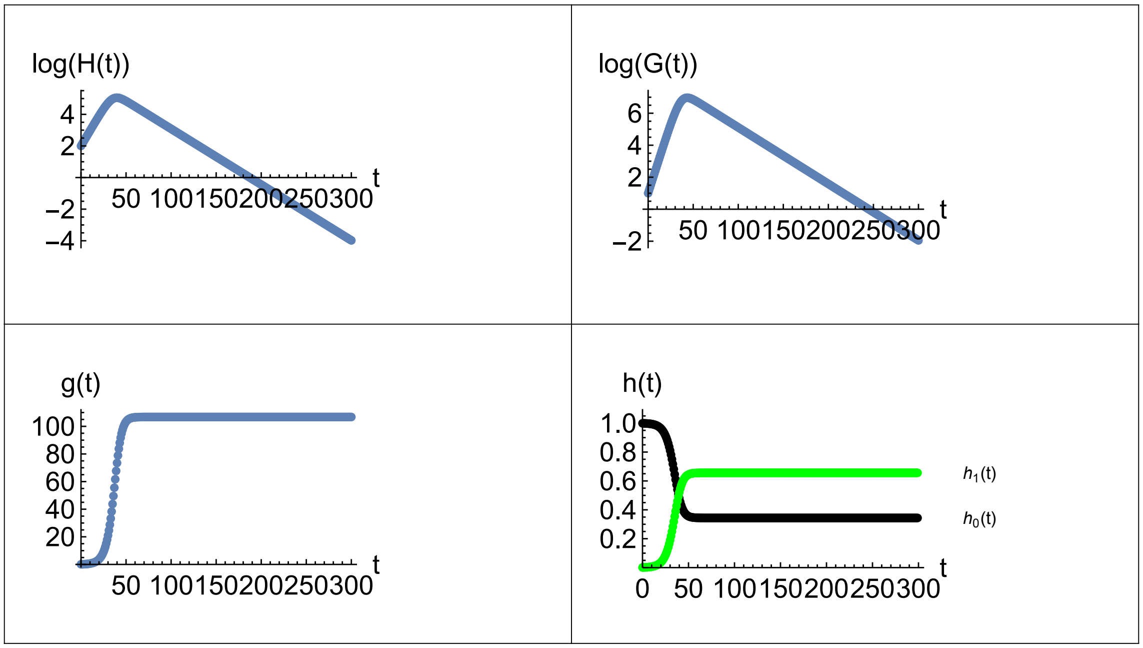


Figure S6: *Trajectories of* $\log_{10}(H(t))$, $\log_{10}(G(t))$ *and* $g(t)$ *through time for a deleterious microbe. Here,* $H(0) = 100$, $G(0) = 10$, $d = 0.01$, $W_0 = 5/4$, $W_1 = 3/4$, $k = 200$ *and* $t_{\max} = 300$. *Asymptotically,* $R_o = 0.922$, $g_o = 106.766$, $h_{0o} = 0.344$ *and* $h_{1o} = 0.656$ *where* $R_{oH} = H(t_{\max})/H(t_{\max} - 1)$ *and* $R_{oG} = G(t_{\max})/G(t_{\max} - 1)$.

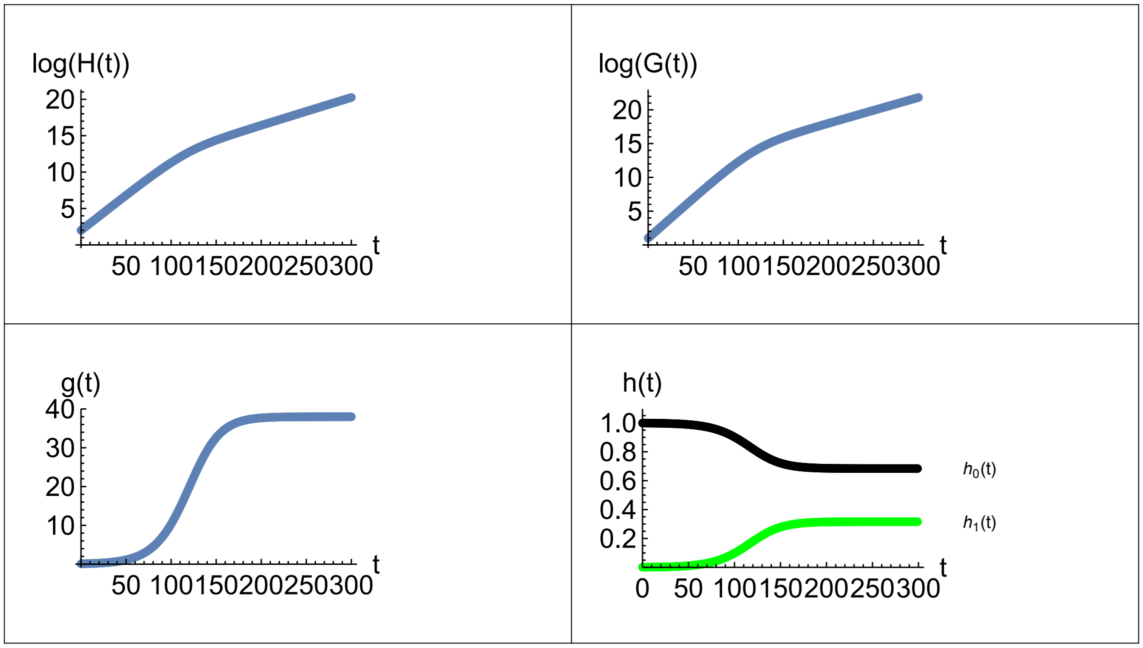


Figure S7: *Trajectories of* $\log_{10}(H(t))$, $\log_{10}(G(t))$ *and* $g(t)$ *through time for a deleterious microbe. Here,* $H(0) = 100$, $G(0) = 10$, $d = 0.01$, $W_0 = 5/4$, $W_1 = 3/4$, $k = 175$ *and* $t_{\max} = 300$. *Asymptotically,* $R_o = 1.091$, $g_o = 37.997$, $h_{0o} = 0.684$ *and* $h_{1o} = 0.326$ *where* $R_{oH} = H(t_{\max})/H(t_{\max} - 1)$ *and* $R_{oG} = G(t_{\max})/G(t_{\max} - 1)$.

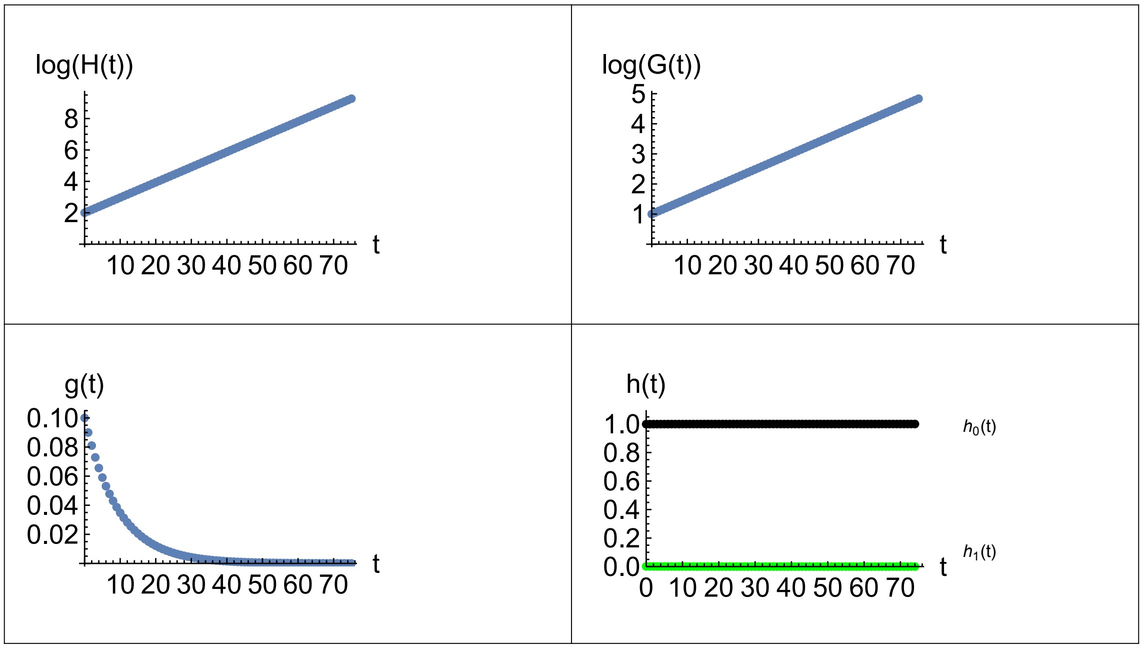


Figure S8: *Trajectories of* $\log_{10}(H(t))$, $\log_{10}(G(t))$ *and* $g(t)$ *through time for a deleterious microbe. Here,* $H(0) = 100$, $G(0) = 10$, $d = 0.01$, $W_0 = 5/4$, $W_1 = 3/4$, $k = 150$ *and* $t_{\max} = 75$. *Asymptotically,* $R_{oH} \approx 1.251$, $R_{oG} \approx 1.125$, $g_o \to 0$, $h_{0o} \to 1$ *and* $h_{1o} \to 0$ *where* $R_{oH} = H(t_{\max})/H(t_{\max} - 1)$ *and* $R_{oG} = G(t_{\max})/G(t_{\max} - 1)$.

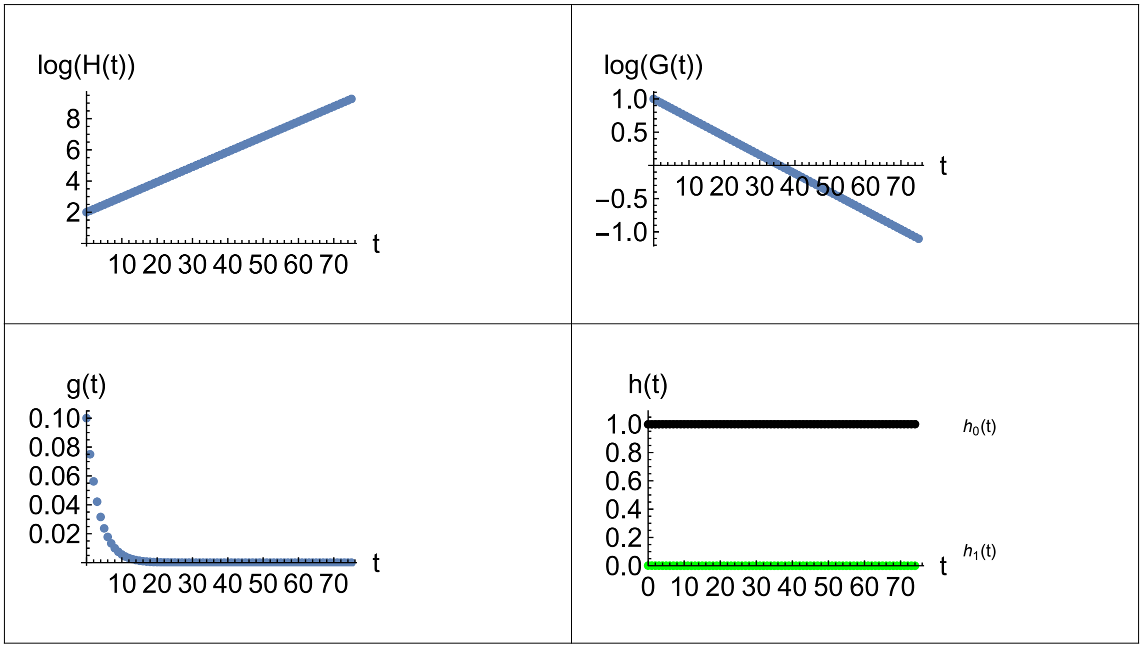


Figure S9: *Trajectories of* $\log_{10}(H(t))$, $\log_{10}(G(t))$ *and* $g(t)$ *through time for a deleterious microbe. Here,* $H(0) = 100$, $G(0) = 10$, $d = 0.01$, $W_0 = 5/4$, $W_1 = 3/4$, $k = 125$ *and* $t_{\max} = 75$. *Asymptotically,* $R_{oH} \approx 1.25$, $R_{oG} \approx 0.938$, $g_o \to 0$, $h_{0o} \to 1$ *and* $h_{1o} \to 0$ *where* $R_{oH} = H(t_{\max})/H(t_{\max} - 1)$ *and* $R_{oG} = G(t_{\max})/G(t_{\max} - 1)$.

### Holobiont with Two Microbial Strains

#### Population Dynamic Model

This section develops equations for the population dynamics for a holobiont with a two-strain microbiome. The microbes of each strain are understood as supplying "genes" for the holobiont and thus comprise its "hologenome". The abundance of a microbial strain is understood to be the "copy number" of the gene from that strain. The "hologenotypes" consist of {0,0}, $\{k_1, 0\}$, $\{0, k_2\}$, and $\{n_1, n_2\}$ copies of the genes in both strains, where $k_1$ and $k_2$ are the within-host equilibrium abundances of single-strain microbiomes consisting of strain-1 and of strain-2 respectively. The $n_1$ and $n_2$ are the within-host equilibrium abundances in a two-strain microbiome obtained from a within-host population-dynamic model such as the Lotka-Volterra competition equations. The frequencies of these hologenotypes are $h_{00}(t)$, $h_{10}(t)$, $h_{02}(t)$ and $h_{12}(t)$ for the hologenotypes, {0,0}, $\{k_1, 0\}$, $\{0, k_2\}$, and $\{n_1, n_2\}$ respectively, where $h_{00}(t) + h_{10}(t) + h_{02}(t) + h_{12}(t) \equiv 1$. The equations developed in this section predict these hologenotype frequencies through time as a result of holobiont selection together with the microbial and host population dynamics.

The life cycle of a holobiont with a microbiome consisting of two horizontally transmitted microbial strains is diagrammed in right panel of Figure 2 in the main article. A macro time step begins with the state variables $H(t)$, $G_1(t)$ and $G_2(t)$ where $H(t)$ is number of empty hosts in the host pool at time $t$, and $G_1(t)$ and $G_2(t))$ are the number of microbes of strain-1 and strain-2 in the microbial pool at time $t$. The total number of microbes in the microbial source pool at time $t$ is $G(t) \equiv G_1(t) + G_2(t)$. The ratio of strain-1 microbes to hosts in their source pools is $g_1(t) \equiv G_1(t)/H(t)$, and similarly $g_2(t) \equiv G_2(t)/H(t)$. The ratio of both microbe strains combined to hosts in their source pools is $g(t) \equiv G(t)/H(t)$. The frequencies of strain-1 and strain-2 in the microbial pool at time $t$ are $p_1(t) \equiv G_1(t)/G(t)$ and $p_2(t) \equiv G_2(t)/G(t)$ where $p_1(t) + p_2(t) \equiv 1$. The Poisson density parameters for strain-1 and strain-2 at time $t$ are $\mu_1(t) \equiv d_1 g_1(t)$ and $\mu_2(t) \equiv d_2 g_2(t)$ where $d_1$ is the dilution factor for strain-1 and $d_2$ is the dilution factor for strain-2.

The two strains independently colonize the empty hosts according to a Poisson distribution. To simplify the subsequent notation, put the probability of a host being colonized by one or more microbes of strain-1 as $P_1(t) \equiv (1 - e^{-\mu_1(t)})$ and for strain-2 as $P_2(t) \equiv (1 - e^{-\mu_2(t)})$. Accordingly, the fraction of empty hosts not colonized by any microbes of strain-1 is $(1 - P_1(t))$ and the fraction not colonized by any microbes of strain-2 is $(1 - P_2(t))$. Therefore, after the microbiome colonization phase,

$$
\begin{aligned}
H'(t,0,0) &= (1-P_1(t))\,(1-P_2(t))\,H(t) \\
H'(t,\bullet,0) &= P_1(t)\,(1-P_2(t))\,H(t) \\
H'(t,0,\bullet) &= (1-P_1(t))\,P_2(t)\,H(t) \\
H'(t,\bullet,\bullet) &= P_1(t)\,P_2(t)\,H(t)
\end{aligned}
$$

where $H(t,n_1,n_2)$ is the number of holobionts with a hologenotype consisting of $n_1$ microbes of strain-1 and $n_2$ microbes of strain-2 at time $t$. As before, the raised dot (•) indicates that one or more microbes are present. Thus, $H'(t,0,0)$ indicates the number of hosts that were not colonized by either species of microbes, and $H'(t,\bullet,\bullet)$ indicates the number that were colonized by both strains, and similarly for the number of hosts that were colonized by only one strain.

Once the juvenile hosts have been initially populated, the microbes proliferate within their hosts, coming to an equilibrium microbiome community structure. The process of attaining the population-dynamic equilibrium within the hosts erases the initial conditions with which the hosts were colonized. Thus, all hosts colonized with one or more of microbes of any particular strain come to the same equilibrium abundance characteristic of that strain. Hence, the equilibrium abundances of single-strain microbiomes consisting of only strain-1 or strain-2 are $k_1$ and $k_2$ respectively, regardless of the number of colonizing microbes, provided the colonizing microbes were all from the same strain. Similarly, the equilibrium abundances in a dual-strain microbiome consisting of strain-1 and strain-2 are $n_1$ and $n_2$ regardless of the number of colonizing microbes provided that both strains are included among the colonists in each host. The $n_1$ and $n_2$ are the equilibrium within-host microbe population sizes from a species-interaction model such as the Lotka-Volterra (LV) competition equations or other appropriate species-interaction model. Therefore, after the microbiome proliferation phase, the number of holobionts of each hologenotype are

$$
\begin{aligned}
H''(t,0,0) &= (1-P_1(t))\,(1-P_2(t))\,H(t) \\
H''(t,k_1,0) &= P_1(t)\,(1-P_2(t))\,H(t) \\
H''(t,0,k_2) &= (1-P_1(t))\,P_2(t)\,H(t) \\
H''(t,n_1,n_2) &= P_1(t)\,P_2(t)\,H(t)
\end{aligned}
$$

Next, the holobiont fitnesses depend on the hologenotype. Let the holobiont fitness of the hologenotypes, $(0,0)$, $(k_1,0)$, $(0,k_2)$ and $(n_1,n_2)$ be $W_0$, $W_1$, $W_2$ and $W_{12}$, respectively.

$$\begin{aligned}
H'''(t,0,0) &= W_0\,(1-P_1(t))\,(1-P_2(t))\,H(t)\\
H'''(t,k_1,0) &= W_1\,P_1(t)\,(1-P_2(t))\,H(t)\\
H'''(t,0,k_2) &= W_2\,(1-P_1(t))\,P_2(t)\,H(t)\\
H'''(t,n_1,n_2) &= W_{12}\,P_1(t)\,P_2(t)\,H(t)
\end{aligned}$$

These holobionts then release their microbiomes into the microbial source pool and their empty juvenile hosts into the host source pool.

Accordingly, the number of empty juvenile hosts in the host source pool at $t+1$, $H(t+1)$, is

$$\begin{aligned}
H(t+1) &= H'''(t,0,0) + H'''(t,k_1,0) + H'''(t,0,k_2) + H'''(t,n_1,n_2)\\
&= \Big( W_0\,(1-P_1(t))\,(1-P_2(t)) + W_1\,P_1(t)\,(1-P_2(t)) +\\
&\quad W_2\,(1-P_1(t))\,P_2(t) + W_{12}\,P_1(t)\,P_2(t) \Big)\,H(t)\\
&= \overline{W}(t)\,H(t) \qquad (18)
\end{aligned}$$

where

$$\overline{W}(t) = (1-P_1(t))\,(1-P_2(t))\,W_0 + P_1(t)\,(1-P_2(t))\,W_1 + (1-P_1(t))\,P_2(t)\,W_2 + P_1(t)\,P_2(t)\,W_{12} \qquad (19)$$

is the mean holobiont fitness after microbe colonization averaged over all possible hologenotypes.

Turning now to the microbes, consider the number of each strain within the microbiome. As before, the products of within-host microbe population sizes and the holobiont fitnesses can be combined into measures of multilevel microbe fitness per holobiont. These measures represent simultaneous success in both the within-holobiont $K$-selection and the between-holobiont $r$-selection. The multilevel microbe fitnesses per holobiont are

$$\begin{aligned}
w_{1,1} &\equiv k_1\,W_1\\
w_{1,12} &\equiv n_1\,W_{12}\\
w_{2,12} &\equiv n_2\,W_{12}\\
w_{2,2} &\equiv k_2\,W_2 \qquad (20)
\end{aligned}$$

These coefficients refer to the multilevel success of a specific microbe strain within a specific microbiome. Thus, $w_{1,1}$ is the multilevel fitness per holobiont of a strain-1 microbe in a single-strain microbiome consisting only of strain-1. The $w_{1,12}$ is the multilevel fitness per holobiont of a strain-1 microbe in a dual strain microbiome, and so forth. The notation convention is that upper case $W$'s refer to holobiont fitnesses and lower case $w$'s refer to microbe fitnesses. Next, the average multilevel microbe fitness per holobiont of strain-1 and strain-2 respectively, are

$$\begin{aligned}\overline{w_1}(t) &= w_{1,1}\,(1-P_2(t)) + w_{1,12}\,P_2(t)\\ \overline{w_2}(t) &= w_{2,12}\,P_1(t) + w_{2,2}\,(1-P_1(t))\end{aligned} \tag{21}$$

The $\overline{w_1}(t)$ is the multilevel fitness per holobiont of a strain-1 microbe averaged over the probability of being in a host by itself, $w_{1,1}$, and the probability of being in a host together with strain-2, $w_{1,12}$, and conversely for $\overline{w_2}(t)$. Finally, the net multilevel fitness per holobiont, taking into account the probability of colonization, is

$$\begin{aligned}w_1(t) &= P_1(t)\,\overline{w_1}(t)\\ w_2(t) &= P_2(t)\,\overline{w_2}(t)\end{aligned} \tag{22}$$

And the net microbe fitness per holobiont of both strains combined is

$$w(t) = w_1(t) + w_2(t) \tag{23}$$

The number of strain-1 microbes in the microbial source pool at $t+1$, $G_1(t+1)$, is

$$\begin{aligned}G_1(t+1) &= k_1\,H'''(t,k_1,0) + n_1\,H'''(t,n_1,n_2)\\ &= \Big(k_1\,W_1\;P_1(t)\,(1-P_2(t)) + n_1\,W_{12}\;P_1(t)\,P_2(t)\Big)\,H(t)\\ &= \Big(P_1(t)\,(w_{1,1}\,(1-P_2(t)) + w_{1,12}\,P_2(t))\Big)\,H(t)\\ &= P_1(t)\,\overline{w_1}(t)\,H(t)\\ &= w_1(t)\,H(t)\end{aligned} \tag{24}$$

Similarly,

$$G_2(t+1) = w_2(t)\,H(t) \tag{25}$$

Thus, the basic population-dynamic model for a holobiont with two microbial strains is given by three dynamical equations: Eqs. 18, 24 and 25. Furthermore, given $H(t)$, $G_1(t)$, and $G_2(t)$, the following quantities can be recorded:

$$G(t) = G_1(t) + G_2(t) \tag{26}$$

$$\begin{aligned}g(t) &= G(t)/H(t)\\ g_1(t) &= G_1(t)/H(t)\\ g_2(t) &= G_2(t)/H(t)\end{aligned} \tag{27}$$

$$\begin{aligned}h_{00}(t) &= (1-P_1(t))\,(1-P_2(t))\\ h_{10}(t) &= P_1(t)\,(1-P_2(t))\\ h_{02}(t) &= (1-P_1(t))\,P_2(t)\\ h_{12}(t) &= P_1(t)\,P_2(t)\end{aligned} \tag{28}$$

$$p_1(t+1) = w_1(t)/w(t)$$
$$p_2(t+1) = w_2(t)/w(t) \quad (29)$$

These quantities, together with $H(t)$, $G_1(t)$, and $G_2(t)$ are included in the subsequent figures.

#### No Selective Neutrality

A fundamental feature of classical population genetics is the possibility of selective neutrality provided the mating system consists of random union of gametes (or random mating). Recall, for example, the classic model for selection on a diploid locus with two alleles, $A_1$ and $A_2$. If the fitnesses for the three genotypes, $A_1A_1$, $A_1A_2$ and $A_2A_2$ are equal, i.e., $W_{11} = W_{12} = W_{22}$, then any initial allele frequency, $p$, persists unchanged. That is, the random union of gametes process does not itself alter the allele frequencies—only "forces" of evolution such as natural selection, drift, mutation and so forth can cause allele frequency change. This result is the familiar Hardy-Weinberg Law.

The process whereby empty hosts in the host source pool are colonized by microbes from the microbial source pool is, for the hologenome, the conceptual analogue of the mating system for the nuclear genome. The colonization process produces a random assortment of microbe strains in the hosts according to Poisson sampling analogous to how the mating system produces a random assortment of alleles in the nucleus according to Binomial sampling. However, the Poisson colonization process is *not* neutral and it produces changes in the frequency of the microbial strains by itself even if the microbe strains are functionally identical. There is no analogue of the Hardy-Weinberg Law for the hologenome.

Figure S10 illustrates the iteration of Eqs. 18, 24, and 25. The properties of both strains are identical. However, the initial frequency of strain-1 is 0.8 and of strain-2 is 0.2. After 20 generations, both strain frequencies converge to 0.5. Thus, the colonization process itself influences the strain frequencies. Therefore, the effect of any within-holobiont $K$-selection between the microbial strains and of any between-holobiont $r$-selection on the entire holobionts both play out upon the non-neutral stage supplied by the colonization process. The Poisson colonization process supplies a strong pull to the center, and the effect of any selection will be combined with this central pull to yield a net result.

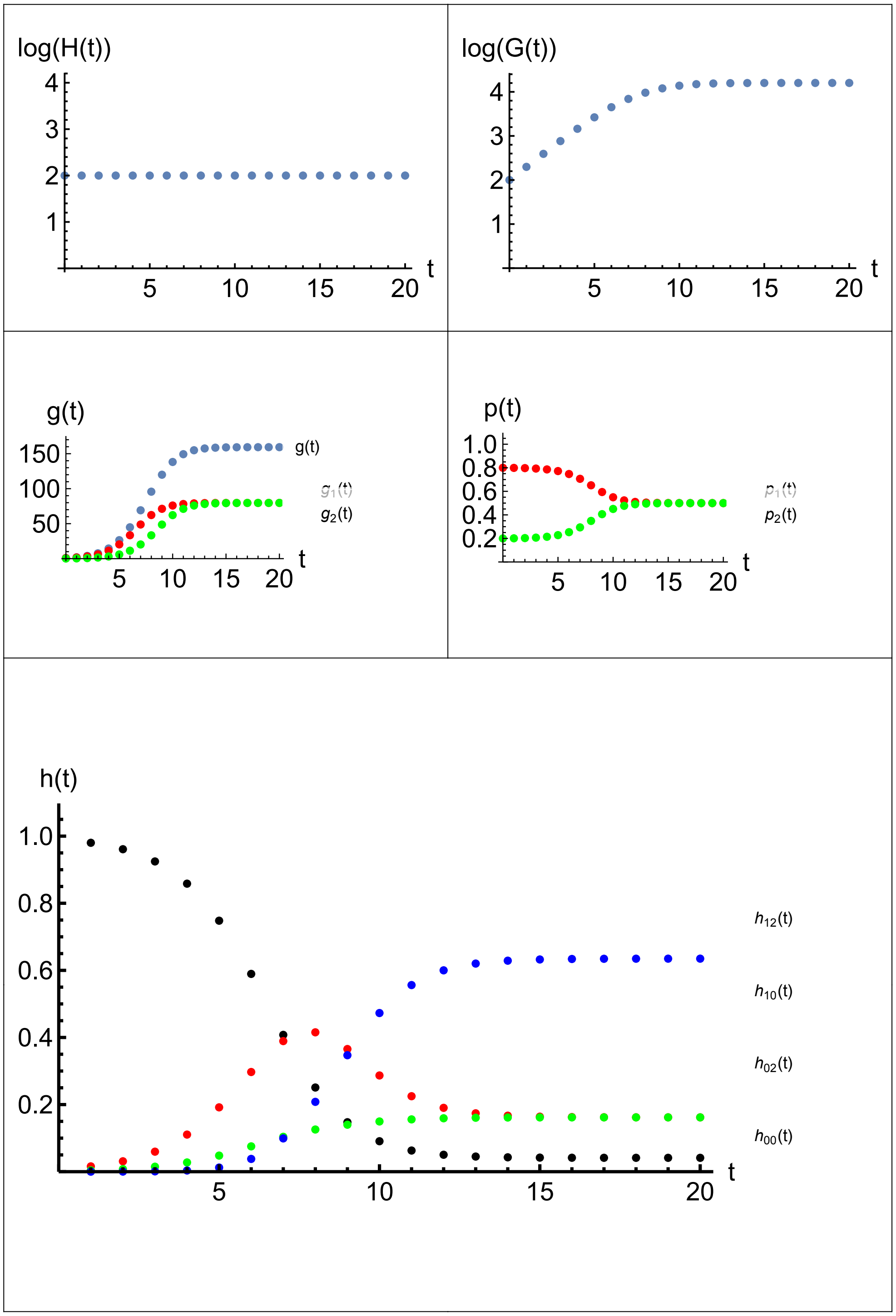


Figure S10: *Host with two identical strains of microbes. The strains start from different initial conditions.* $d_1 = .02$, $d_2 = .02$, $k_1 = 100$, $k_2 = 100$, $a_{12} = 0$, $a_{21} = 0$, $W_0 = 1$, $W_1 = 1$, $W_2 = 1$, $W_{12} = 1$, $H_{\text{init}} = 100$, $G_{1\text{init}} = 80$, $G_{2\text{init}} = 20$, $t_{\max} = 20$. *Asymptotically,* $R_{oH} \approx 1.00$, $R_{oG} \approx 1.00$, $g_o = 159.36$, $g_{1o} = 79.68$, $g_{2o} = 79.68$, $p_{1o} = 0.5$, $p_{2o} = 0.5$, $h_{00o} = 0.04$, $h_{10o} = 0.16$, $h_{02o} = 0.16$, *and* $h_{12o} = 0.63$ *where* $R_{oH} = H(t_{\max})/H(t_{\max}-1)$ *and* $R_{oG} = G(t_{\max})/G(t_{\max}-1)$.

#### Asymptotic Properties

Figure S10 shows that the holobiont population size and total number of microbes approach exponential growth after about 10 generations at which time the ratios of microbes to hosts in the source pools, the frequency of the strains in the microbial source pool, and the hologenotype frequencies all approach equilibrium. The figure's caption details the asymptotic outcome obtained by numerically iterating Eqs. 18, 24, and 25.

To obtain mathematical formulae for the outcome revealed in Figure S10 for a holobiont with a two-strain microbiome, an approach can be developed that parallels Eqs. 9–11 for a holobiont with a single strain of microbiome. Specifically, the two-strain counterpart to Eq. 9 is

$$W_o(g_{1o}, g_{2o}) = h_{00o}\, W_0 + h_{10o}\, W_1 + h_{02o}\, W_2 + h_{12o}\, W_{12} \tag{30}$$

where

$$\begin{aligned} \mu_{1o} &= d_1\, g_{1o} \\ \mu_{2o} &= d_2\, g_{2o} \\ P_{1o} &= 1 - e^{-\mu_{1o}} \\ P_{2o} &= 1 - e^{-\mu_{2o}} \end{aligned}$$

$$\begin{aligned} h_{00o} &= (1 - P_{1o})\,(1 - P_{2o}) \\ h_{10o} &= P_{1o}\,(1 - P_{2o}) \\ h_{02o} &= (1 - P_{1o})\,P_{2o} \\ h_{12o} &= P_{1o}\,P_{2o} \end{aligned}$$

The two-strain counterparts of Eq. 10 leading to the equilibrium solution

$$W_o(g_{1o}, g_{2o}) = \frac{P_{1o}\overline{w_{1o}}}{g_{1o}} \tag{31}$$

$$W_o(g_{1o}, g_{2o}) = \frac{P_{2o}\overline{w_{2o}}}{g_{2o}} \tag{32}$$

where

$$
\begin{aligned}
n_1 &= \frac{k_1 - a_{12}\,k_2}{1 - a_{12}\,a_{21}} \\
n_2 &= \frac{k_2 - a_{21}\,k_1}{1 - a_{12}\,a_{21}} \\
w_{1,1} &= k_1\,W_1 \\
w_{2,2} &= k_2\,W_2 \\
w_{1,12} &= n_1\,W_{12} \\
w_{2,12} &= n_2\,W_{12} \\
\overline{w_{1o}} &= w_{1,1}\,(1 - P_{2o}) + w_{1,12}\,P_{2o} \\
\overline{w_{2o}} &= w_{2,2}\,(1 - P_{1o}) + w_{2,12}\,P_{1o}
\end{aligned}
$$

For the two-strain counterpart of the equilibrium solution, Eq. 11, put Eq. 30 equal to Eq. 31, and Eq. 30 equal to Eq. 32, yielding two simultaneous equations for $g_{1o}$ and $g_{2o}$

$$g_{1o} = \frac{P_{1o}\overline{w_{1o}}}{h_{00o}\,W_0 + h_{10o}\,W_1 + h_{02o}\,W_2 + h_{12o}\,W_{12}} \tag{33}$$

$$g_{2o} = \frac{P_{2o}\overline{w_{2o}}}{h_{00o}\,W_0 + h_{10o}\,W_1 + h_{02o}\,W_2 + h_{12o}\,W_{12}} \tag{34}$$

Using the parameters from Figure S10 for Eqs. 33 and 34, the FindRoot function in *Mathematica* (with a seed of {100,100}) immediately returns the answer as $g_{1o} = 79.68$ and $g_{2o} = 79.68$, in agreement with the results previously obtained by numerical iteration. Given this solution for for $g_{1o}$ and $g_{2o}$, all the remaining quantities can be calculated. Specifically, $W_o = 1$, $h_{00o} = 0.04$, $h_{10o} = 0.16$, $h_{02o} = 0.16$, $h_{12o} = 0.63$, $p_{1o} = 0.5$ and $p_{2o} = 0.5$.

#### Conditions for Increase When Rare

Although a simultaneous solution to Eqs. 33 and 34 indicates a possible equilibrium, whether this equilibrium is stable depends on whether the strains can each increase when rare, given that the other strain is established. Paralleling the derivation of Eq. 13 for a single strain, the condition for increase when rare for strain-1 is found by first developing the formula for $\Delta G_1$ from Eq. 24 as

$$
\begin{aligned}
\Delta G_1 &\equiv G_1(t+1) - G_1(t) \\
&= w_1\,H - G_1
\end{aligned}
$$

where $w_1$ is defined in Eqs. 20–22. With $P_1$ and $P_2$ written explicitly the equation for $\Delta G_1$ becomes

$$\Delta G_1 = \left(k_1\,W_1\,e^{-d_2\frac{G_2}{H}} + n_1\,W_{12}(1 - e^{-d_2\frac{G_2}{H}})\right)\,(1 - e^{-d_1\frac{G_1}{H}})\,H - G_1$$

To see if $\Delta G_1$ becomes positive with the addition of some strain-1 microbes to a holobiont population where strain-2 is already established, check that the derivative of $\Delta G_1$ with respect to $G_1$ is positive when $G_1 = 0$ and $G_2/H = \hat{g}_2$ where $\hat{g}_2$ is the value of $g_2$ when it is established in the absence of strain-1—this is found from the single strain formula, Eq. 11. So, differentiate $\Delta G_1$ with respect to $G_1$ and evaluate at $G_1 = 0$ and $G_2/H = \hat{g}_2$, yielding

$$\frac{\partial \Delta G_1}{\partial G_1}|_{G_1=0,\, G_2/H=\hat{g}_2} = d_1 \, \left(k_1 \, W_1 \, e^{-d_2 \hat{g}_2} + n_1 \, W_{12} \, (1 - e^{-d_2 \hat{g}_2})\right) - 1$$

Hence, strain-1 when rare can enter the holobiont population in which strain-2 is already established at $g_2 = \hat{g}_2$ if

$$d_1 \, \left(k_1 \, W_1 \, e^{-d_2 \hat{g}_2} + n_1 \, W_{12} \, (1 - e^{-d_2 \hat{g}_2})\right) > 1 \tag{35}$$

Similarly, strain-2 when rare can enter the holobiont population in which strain-1 is already established at $g_1 = \hat{g}_1$ if

$$d_2 \, \left(k_2 \, W_2 \, e^{-d_1 \hat{g}_1} + n_2 \, W_{12} \, (1 - e^{-d_1 \hat{g}_1})\right) > 1 \tag{36}$$

Eqs. 35 and 36 are the two-strain counterparts of Eq. 13 for a single strain. These requirements means that for each strain to enter the holobiont population in which the other strain is established, each strain of microbes must be able to do better than to merely replace themselves taking into account their dilution factor and both their average abundance realized within their hosts and their average holobiont fitness as encountered upon entry.

Returning to the case illustrated in Figure S10 where the microbes are identical, $k_1 = k_2 = n_1 = n_2 = k$, $d_1 = d_2 = d$, and $W_0 = W_1 = W_2 = W_{12} = W$, Eqs. 35 and 36 both reduce down to

$$d \, k \, W > 1,$$

which is the counterpart to Eq. 13. In Figure S10, $d = 0.02$, $k = 100$, and $W = 1$, so $d \, k \, W = 2 > 1$, so both strains can increase when the other is rare, as the figure indeed illustrates.

#### Hologenome Polymorphism from Microbial Dynamics

Hologenome polymorphism can result from microbial dynamics alone without any selection at the holobiont level. To illustrate, suppose the two strains of microbes compete with each other within the host but each strain has the same effect on the host. From the host's point of view, the strains are identical, but from the microbes' point of view, one strain is superior to the other—say, strain-1 out-competes strain-2 whenever the two strains are within the same host. This assumption requires strong competitive asymmetry which, with Lotka-Volterra competition, implies that $a_{12} < k_1/k_2$ and $a_{21} > k_2/k_1$. Can strain-2 persist in the holobiont population despite losing in competition to strain-1 whenever both occur together?

To model this case, let $W_0 = W_1 = W_2 = W_{12} = W$. Then, Eq. 30 reduces to

$$W_o = W$$

Also, if strain-1 always eliminates strain-2 when both colonize the same host, then $n_1 = k_1$ and $n_2 = 0$. Hence

$$\begin{aligned} w_{1,1} &= k_1 W \\ w_{2,2} &= k_2 W \\ w_{1,12} &= k_1 W \\ w_{2,12} &= 0 \end{aligned}$$

Therefore, in this case

$$\begin{aligned} \overline{w_{1o}} &= k_1 W \\ \overline{w_{2o}} &= (1 - P_{1o})\, k_2 W \end{aligned}$$

and the coexistence conditions for strain-1 and strain-2, Eqs. 33 and 34, reduce to

$$\begin{aligned} g_{1o} &= P_{1o}\, k_1 \\ g_{2o} &= P_{2o}\, (1 - P_{1o})\, k_2 \end{aligned}$$

Putting

$$\begin{aligned} P_{1o} &= 1 - e^{-d_1 g_{1o}} \\ P_{2o} &= 1 - e^{-d_2 g_{2o}} \end{aligned}$$

yields the counterpart of Eq. 33 and 34 for this case

$$g_{1o} = (1 - e^{-d_1 g_{1o}})\, k_1 \tag{37}$$

$$g_{2o} = (1 - e^{-d_2 g_{2o}})\, e^{-d_1 g_{1o}}\, k_2 \tag{38}$$

These are simultaneous equations for $g_{1o}$ and $g_{2o}$. Eq. 37 is a function of only $g_{1o}$, so this equation can be solved for $g_{1o}$ and the answer plugged into Eq. 38 to yield $g_{2o}$.

Turning now to the conditions for invasion when rare, the condition for strain-1, Eq. 35, reduces to

$$d_1\, k_1\, W > 1 \tag{39}$$

irrespective of $\hat{g}_2$. Similarly, the condition for strain-2, Eq. 36, reduces to

$$d_2\, k_2\, W\, e^{-d_1 \hat{g}_1} > 1 \tag{40}$$

Because $e^{-d_1 \hat{g}_1}$ is the probability that a host has *not* been colonized by strain-1, Eq. 40 shows that the condition for increase when rare for strain-2 depends on its success in the hosts available to it, namely those uncolonized by strain-1 from which it is excluded. In particular, Eq. 40 implies that a critical $k_2$, say $k_{2c}$ exists

$$k_{2c} = \frac{e^{d_1 \hat{g}_1}}{d_2 W} \tag{41}$$

such that if $k_2 < k_{2c}$ then the competitively superior strain-1 microbe can exclude the competitively inferior strain-2 microbe from the holobiont population, whereas if $k_2 > k_{2c}$ strain-1 cannot exclude strain-2 from the holobiont population even though it can exclude strain-2 whenever both colonize the same host. In Eq. 41 , $\hat{g}_1$, from Eq. 11 where $W_0 = W_1 = W$, is explicitly

$$\hat{g}_1 = \frac{d_1 k_1 + \text{ProductLog}[-d_1 k_1 e^{-d_1 k_1}]}{d_1}$$

The reason that the strains can coexist despite the competitive asymmetry, provided $k_2 > k_{2c}$, is that the colonization process provides some empty hosts that end up being colonized only by strain-2, thereby providing a statistical refuge for the competitively inferior strain from the competitively superior strain. However, the production of strain-2 microbes in these refuge hosts must be high enough to compensate for their inability to produce anything in those hosts where strain-1 is also present. This result is an instance of a metapopulation model for patch dynamics featuring a colonization-extinction equilibrium.

Figures S11 and S12 illustrate the the dynamics of the host with a two-strain microbiome, assuming that strain-1 competitively excludes strain-2 whenever both colonize the same host. For the parameters of the illustration, the critical value, $k_{2c}$ works out to be 246. In Figure S11, $k_2$ is less than $k_{2c}$, and strain-1 drives strain-2 to extinction in the holobiont population because strain-2 does not produce enough where it is alone in a host to compensate for the loss of production from hosts that both strains colonized. In Figure S12, $k_2$ is greater than $k_{2c}$, and strain-1 and strain-2 coexist in the holobiont population at $g_{1o} = 79.68$ and $g_{2o} = 20.51$. Here, strain-2 does produce enough in hosts where it is alone to compensate for the loss of production in hosts where both strains colonize.

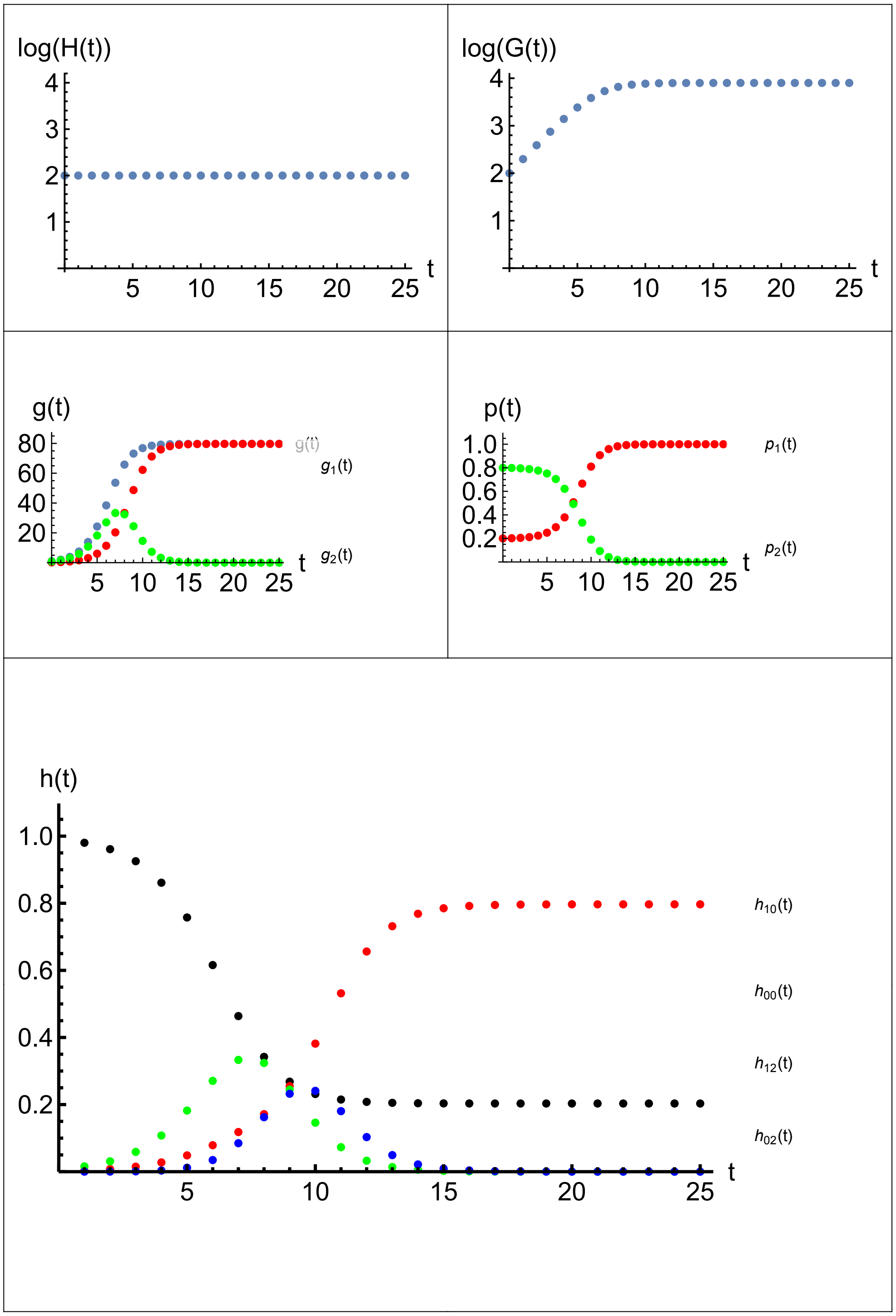


Figure S11: *Host with competitively superior strain-1 and competitively inferior strain-2. Here,* $k_2 < k_{2c} = 246$. *Also* $d_1 = .02$, $d_2 = .02$, $k_1 = 100$, $k_2 = 100$, $W = 1$, $H_{\text{init}} = 100$, $G_{1\text{init}} = 20$, $G_{2\text{init}} = 80$, $t_{\text{max}} = 25$, $a_{12} < 1$ *and* $a_{21} > 1$. *Asymptotically,* $R_{oH} \approx 1.00$, $R_{oG} \approx 1.00$, $g_o = 79.68$, $g_{1o} = 79.68$, $g_{2o} = 0$, $p_{1o} = 1$, $p_{2o} = 0$, $h_{00o} = 0.20$, $h_{10o} = 0.80$, $h_{02o} = 0$, *and* $h_{12o} = 0$ *where* $R_{oH} = H(t_{\text{max}})/H(t_{\text{max}} - 1)$ *and* $R_{oG} = G(t_{\text{max}})/G(t_{\text{max}} - 1)$.

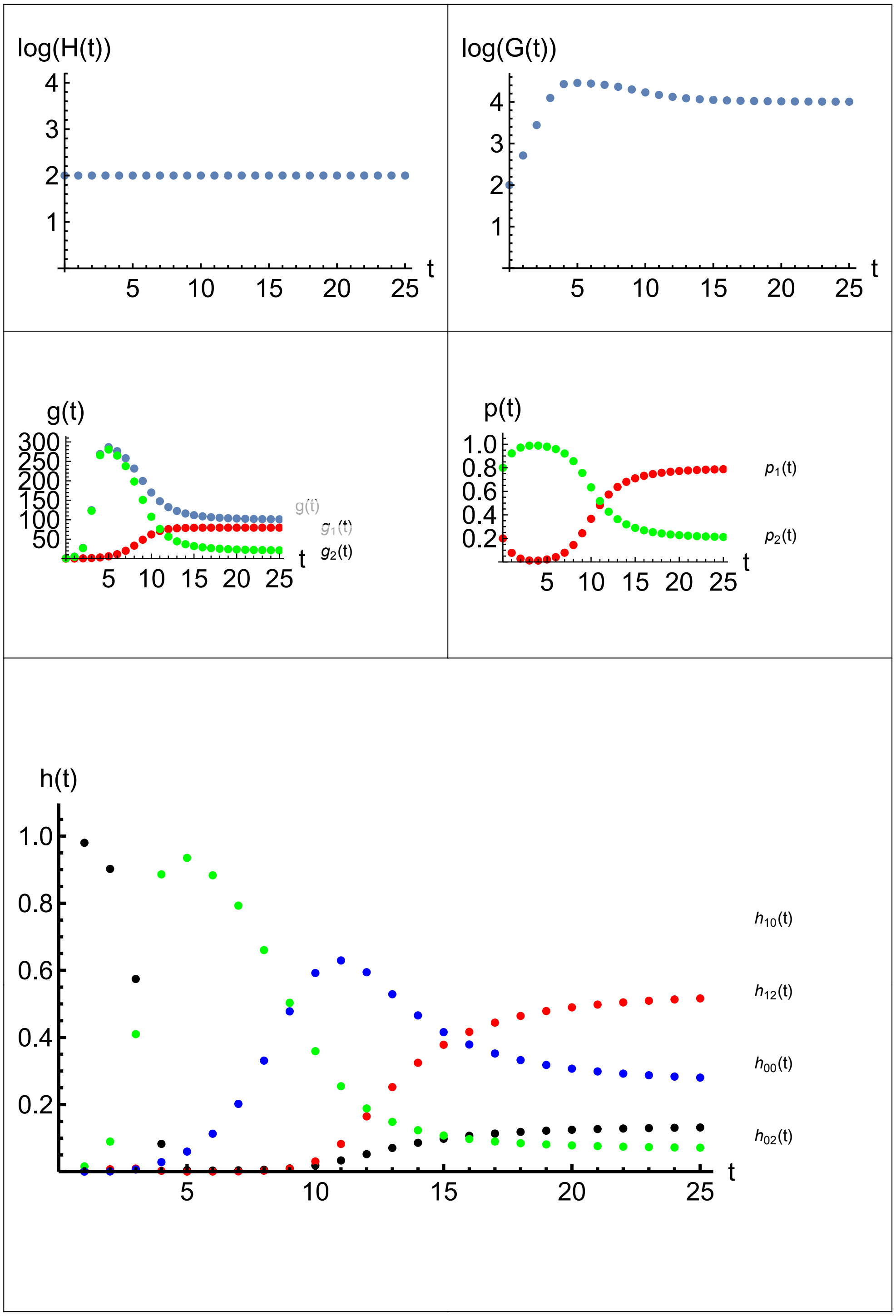


Figure S12: *Host with competitively superior strain-1 and competitively inferior strain-2. Here,* $k_2 > k_{2c} = 246$. *Also* $d_1 = .02$, $d_2 = .02$, $k_1 = 100$, $k_2 = 300$, $W = 1$, $H_{\text{init}} = 100$, $G_{1\text{init}} = 20$, $G_{2\text{init}} = 80$, $t_{\max} = 25$, $a_{12} < 1/3$ *and* $a_{21} > 3$. *Asymptotically,* $R_{oH} \approx 1.00$, $R_{oG} \approx 1.00$, $g_o = 101.13$, $g_{1o} = 79.68$, $g_{2o} = 21.45$, $p_{1o} = 0.79$, $p_{2o} = 0.21$, $h_{00o} = 0.13$, $h_{10o} = 0.52$, $h_{02o} = 0.07$, *and* $h_{12o} = 0.28$ *where* $R_{oH} = H(t_{\max})/H(t_{\max} - 1)$ *and* $R_{oG} = G(t_{\max})/G(t_{\max} - 1)$.

#### Hologenome Polymorphism from Holobiont Selection

Hologenome polymorphism can result from selection at the holobiont level, without any contribution from microbial dynamics. To illustrate, suppose the two strains of microbes are identical and transparent to each other. They do differ however, in their impact on holobiont fitness. That is, $k_1 = k_2 = k$, $d_1 = d_2 = d$, and $a_{12} = a_{21} = 0$, so that $n_1 = k$ and $n_2 = k$.

This scenario for horizontal microbial transmission corresponds to the standard case of vertical nuclear gene transmission in classical population genetics. For example, with one diploid locus and two alleles that have fitnesses, $W_{11}$, $W_{12}$, and $W_{22}$, the well-known condition for polymorphism with random mating is heterozygote superiority, i.e., both $W_{12} > W_{11}$ and $W_{12} > W_{22}$. Conversely, directional selection selection results in fixation of say, allele $A_1$, if both $W_{11} > W_{12}$ and $W_{12} > W_{22}$. Now, suppose that the same two alleles from the nucleus are instead found in the two strains of the microbiome—microbes are the vehicles delivering genes to the host instead of the nucleus. If so, are there any analogues of the conditions for polymorphism and directional selection for genes carried by the microbiome compared to the conditions for polymorphism and fixation of genes carried by the nucleus?

To model this case, Eq. 30 becomes

$$\begin{aligned} W_o(g_{1o}, g_{2o}) &= e^{-d\,g_{1o}}\, e^{-d\,g_{2o}}\, W_0 + (1 - e^{-d\,g_{1o}})\, e^{-d\,g_{2o}}\, W_1 \\ &+ e^{-d\,g_{1o}}\, (1 - e^{-d\,g_{2o}})\, W_2 + (1 - e^{-d\,g_{1o}})\, (1 - e^{-d\,g_{2o}})\, W_{12} \end{aligned} \tag{42}$$

Also,

$$\begin{aligned} w_{1,1} &= k\, W_1 \\ w_{2,2} &= k\, W_2 \\ w_{1,12} &= k\, W_{12} \\ w_{2,12} &= k\, W_{12} \end{aligned}$$

Hence,

$$\begin{aligned} \overline{w_{1o}}(g_{2o}) &= k\, \Big( W_1\, e^{-d\,g_{2o}} + W_{12}\, (1 - e^{-d\,g_{2o}}) \Big) \\ \overline{w_{2o}}(g_{1o}) &= k\, \Big( W_2\, e^{-d\,g_{1o}} + W_{12}\, (1 - e^{-d\,g_{1o}}) \Big) \end{aligned}$$

Therefore the simultaneous equations for the polymorphic solution corresponding to Eqs. 33 and 34 in this case become

$$g_{1o} = \frac{k\, (1 - e^{-d\,g_{1o}})\, \big( W_1\, e^{-d\,g_{2o}} + W_{12}\, (1 - e^{-d\,g_{2o}}) \big)}{W_o(g_{1o}, g_{2o})} \tag{43}$$

$$g_{2o} = \frac{k\, (1 - e^{-d\,g_{2o}})\, \big( W_2\, e^{-d\,g_{1o}} + W_{12}\, (1 - e^{-d\,g_{1o}}) \big)}{W_o(g_{1o}, g_{2o})} \tag{44}$$

The conditions for invasion when rare corresponding to Eqs. 35 and 36 in this case become

$$d\,k\,\Big(W_1\,e^{-d\,\hat{g}_2} + W_{12}\,(1 - e^{-d\,\hat{g}_2})\Big) \;>\; 1 \tag{45}$$
$$d\,k\,\Big(W_2\,e^{-d\,\hat{g}_1} + W_{12}\,(1 - e^{-d\,\hat{g}_1})\Big) \;>\; 1 \tag{46}$$

where $\hat{g}_1$ and $\hat{g}_2$ are the roots, from Eq. 11, of

$$\hat{g}_1 \;=\; \frac{k\,W_1\,(1 - e^{-d\,\hat{g}_1})}{W_0\,e^{-d\,\hat{g}_1} + W_1\,(1 - e^{-d\,\hat{g}_1})} \tag{47}$$
$$\hat{g}_2 \;=\; \frac{k\,W_2\,(1 - e^{-d\,\hat{g}_2})}{W_0\,e^{-d\,\hat{g}_2} + W_2\,(1 - e^{-d\,\hat{g}_2})} \tag{48}$$

Furthermore, for $\hat{g}_1$ and $\hat{g}_2$ to be feasible it is necessary, from Eq. 14, that

$$k \;>\; k_{e1} \equiv \frac{1}{d\,W_1} \tag{49}$$
$$k \;>\; k_{e2} \equiv \frac{1}{d\,W_2} \tag{50}$$

and it is sufficient, from Eq. 15, that

$$k \;>\; k_{m1} \equiv \frac{W_0}{d\,W_1} \tag{51}$$
$$k \;>\; k_{m2} \equiv \frac{W_0}{d\,W_2} \tag{52}$$

to ensure that each strain can individually coexist with the host to begin with.

The expressions in large parentheses in both Eqs. 45 and 46 are averages. In Eq. 45 the expression represents an average between $W_1$ and $W_{12}$ and in Eq. 46, an average between $W_2$ and $W_{12}$. In Eq. 45 , the average shifts to $W_1$ as $\hat{g}_2$ approaches 0 and to $W_{12}$ as $\hat{g}_2$ becomes large. Furthermore, from Eq. 49, $d\,k\,W_1 > 1$ because this condition is necessary for strain-1 to coexist with the host to begin with. Therefore, if $\hat{g}_2$ is low enough, then Eq. 45 is automatically satisfied because of satisfying Eq. 49, whereas as $\hat{g}_2$ increases, then $W_1$ is being averaged in with the $W_{12}$. The same considerations apply to Eq. 46, the condition for strain-2 to increase when rare. The $d\,k\,W_2 > 1$ because this condition is necessary for strain-2 to coexist with the host to begin with. Therefore, if $\hat{g}_1$ is low enough, then Eq. 46 is automatically satisfied because of satisfying Eq. 50, whereas as $\hat{g}_1$ increases, then $W_2$ is being averaged in with the $W_{12}$. Thus, the effectiveness of holobiont selection depends on the size of $\hat{g}_1$ and $\hat{g}_2$. If these are low, then many hosts wind up empty of microbes, thereby denying the holobionts any exposure to holobiont selection. But if $\hat{g}_1$ and $\hat{g}_2$ are high, then most hosts are colonized by microbes and holobiont selection can discriminate among the holobionts depending on their hologenotypes.

The expressions in Eqs. 45 and 46 compare $d$ and the multilevel fitnesses with the number, 1, and not with each another. In contrast, in classical population genetics, whether or not an allele can increase when rare depends on comparing the genotype fitnesses with each other.

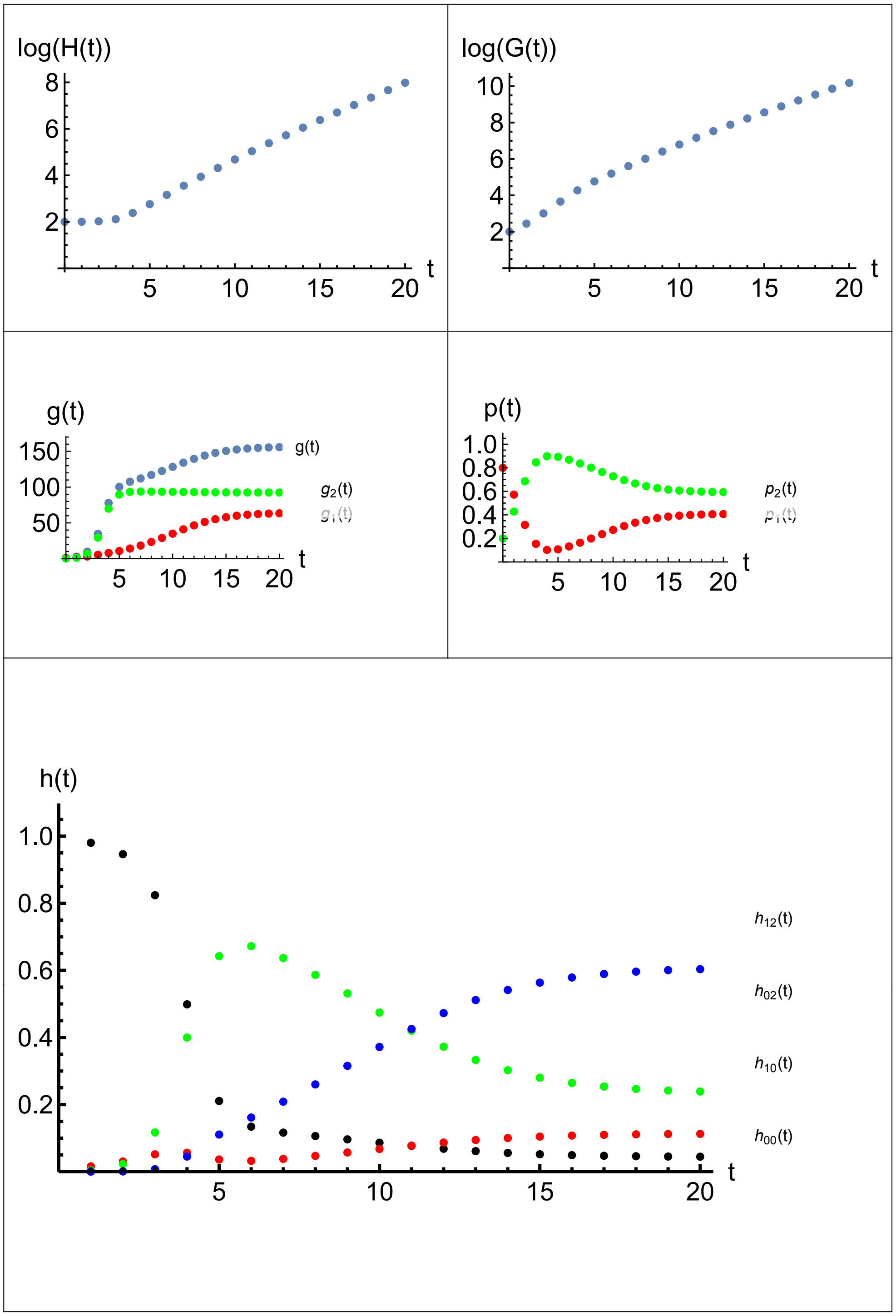


Figure S13: *Holobiont selection favoring strain-2. Here,* $W_0 = 1$, $W_1 = 1$, $W_{12} = 2$, $W_2 = 3$. *Also,* $d_1 = 0.02$, $d_2 = 0.02$, $k_1 = 100$, $k_2 = 100$, $a_{12} = 0$, $a_{21} = 0$, $H_{\text{init}} = 100$, $G_{1\text{init}} = 20$, $G_{2\text{init}} = 80$, $t_{\text{max}} = 20$. *Asymptotically,* $R_{oH} \approx 2.08$, $R_{oG} \approx 2.08$, $g_o = 155.86$, $g_{1o} = 63.41$, $g_{2o} = 92.44$, $p_{1o} = 0.407$, $p_{2o} = 0.593$, $h_{00o} = 0.045$, $h_{10o} = 0.113$, $h_{02o} = 0.239$, *and* $h_{12o} = 0.604$ *where* $R_{oH} = H(t_{\text{max}})/H(t_{\text{max}} - 1)$ *and* $R_{oG} = G(t_{\text{max}})/G(t_{\text{max}} - 1)$.

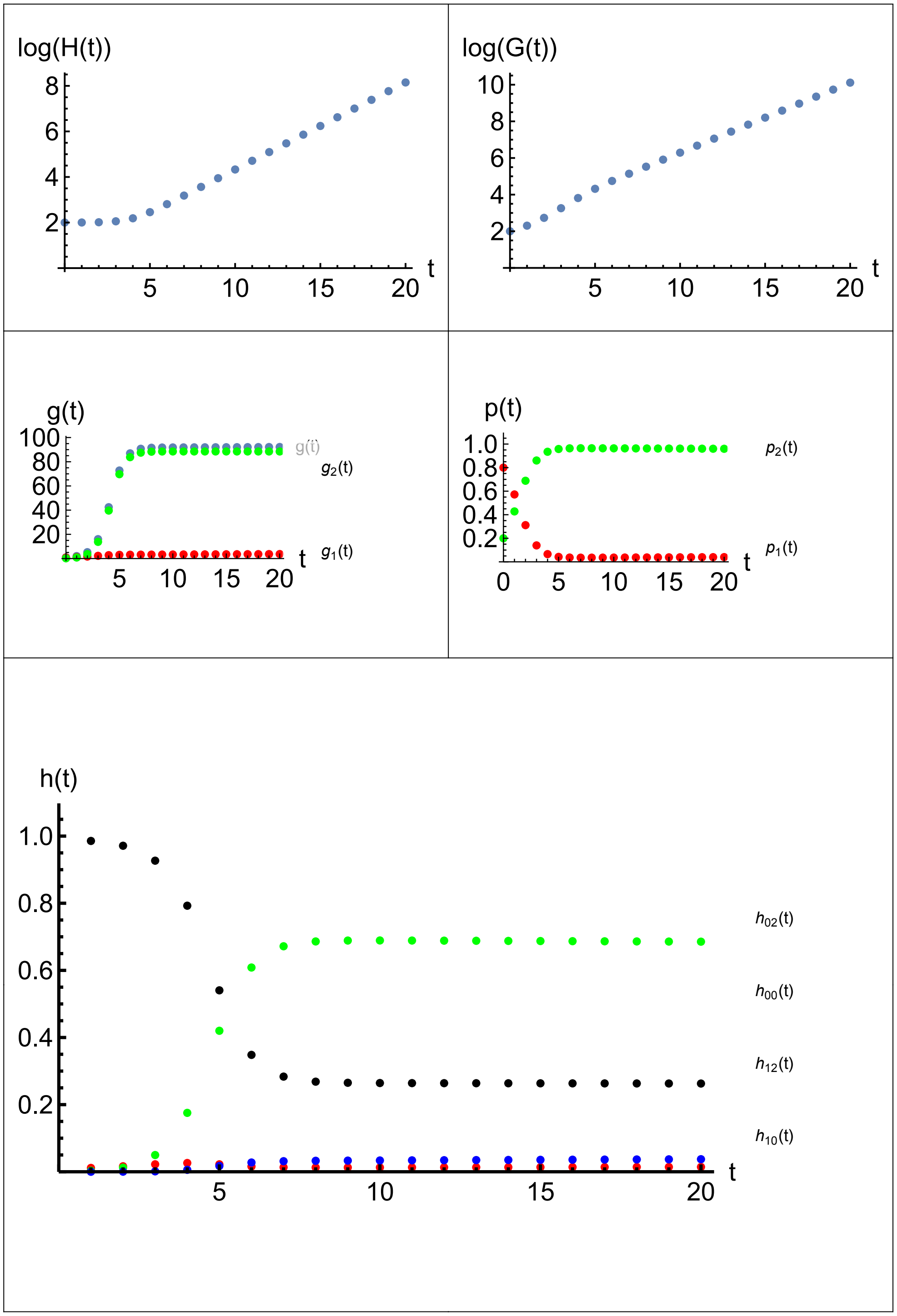


Figure S14: *Holobiont selection favoring strain-2. Here,* $W_0 = 1$, $W_1 = 1$, $W_{12} = 2$, $W_2 = 3$. *Also,* $d_1 = 0.0145$, $d_2 = 0.0145$, $k_1 = 100$, $k_2 = 100$, $a_{12} = 0$, $a_{21} = 0$, $H_{\text{init}} = 100$, $G_{1\text{init}} = 20$, $G_{2\text{init}} = 80$, $t_{\max} = 20$.

**Directional Selection.** Consider a numerical illustration of directional holobiont selection. Suppose as in classical population genetics that two nuclear alleles lead to the genotype fitnesses, $W_{11} = 1$, $W_{12} = 2$, and $W_{22} = 3$. Then $A_1$ cannot increase when rare and the gene pool is fixed for $A_2$, a case of directional selection favoring $A_2$ over $A_1$. Suppose instead the same alleles are lodged in microbial strains leading to the same hologenotype fitnesses, $W_1 = 1$, $W_{12} = 2$, and $W_2 = 3$. Also, let $W_0 = 1$. Then, the LHS of Eq. 45 is a number between $d\,k\,W_1$ and $d\,k\,W_{12}$ depending on the magnitude of $\hat{g}_2$. Because $d\,k\,W_1 > 1$ and $W_{12} > W_1$, the LHS of Eq. 45 is automatically satisfied. Similarly, the LHS of Eq. 46 is a number between $d\,k\,W_{12}$ and $d\,k\,W_2$ depending on $\hat{g}_1$. Both of these are $> 1$ too. Thus, both LHS's are greater than 1 regardless of the size of $\hat{g}_1$ and $\hat{g}_2$. Hence, because Eq. 45 and 46 are both satisfied, a polymorphism is indicated. While some directional selection occurred, a selective sweep did not occur.

This case is illustrated in Figure S13. For this figure, the equilibrium strain frequencies are $p_{1o} = 0.410$ and $p_{2o} = 0.590$ from Eqs. 43 and 44. The equilibrium reveals the strong pull to the center caused by the Poisson colonization dynamics, as was first illustrated in Fig. S10. The selection in favor of strain-2 pulls the frequency of strain-2 above the center at 0.5 and pushes the frequency of strain-1 below the center of 0.5. In another example, not illustrated, the strength of selection favoring strain-2 over strain-1 is increased to $W_1 = 1$, $W_{12} = 4$, and $W_2 = 6$ with $W_0 = 1$. As a consequence. $p_{1o} = 0.396$ and $p_{2o} = 0.604$, showing that increasing the strength of selection does shift the net result further away from the center.

The power of the colonization process to override the holobiont selection is controlled by the colonization parameter, $d$. If $d$ is lowered, then the directional selection is able to drive the frequency of the inferior strain-1 lower. In Figure S13, $d = 0.020$. In contrast, Figure S14 uses $d = 0.0145$. In this figure, the equilibrium strain frequencies now are $p_1 = 0.040$ and $p_2 = 0.960$, showing that directional selection has a much larger effect than in Figure S13.

Figure S15 shows how the equilibrium frequency of strain-1 depends on the colonization parameter. In the example, if $d$ is greater than 0.014, polymorphism results. However, as $d$ is increased much beyond 0.014, the equilibrium frequency of strain-1 quickly approaches 0.5, indicating that the colonization process completely overpowers the directional selection against strain-1 and favoring strain-2. Conversely, if the the colonization process is weak enough, then directional selection does fix the favored allele and eliminates the other, as in classical population genetics.

The colonization process matters because $d$ controls whether the strains are able to express their fitness differences. If $d$ is high, then the strains often co-occur in the same host. Figure S13 (bottom) shows that the two-strain holobiont frequency, $h_{12}(t)$ is the highest frequency. In this situation, both strains often find themselves having the same fitness, $W_{12}$, because they are both together in the same holobiont. Hence, neither can have an advantage over the other. Conversely,

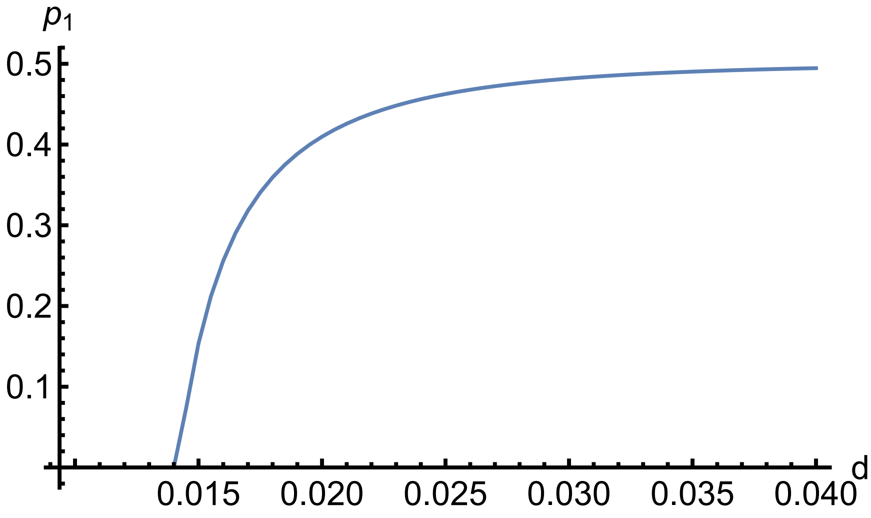


Figure S15: *Equilibrium frequency of strain-1 as a function of the colonization parameter, d. Directional holobiont selection is against strain-1 and favors strain-2. Directional holobiont selection is effective for low d but not high d. Here,* $k = 100$, $W_0 = 1$, $W_1 = 1$, $W_{12} = 2$, $W_2 = 3$

if $d$ is low, then the strains often occupy different hosts by themselves. In Figure S14 (bottom), the strain-2 holobiont frequency, $h_{02}(t)$, is the highest frequency and $h_{12}(t)$ is very low. In this situation the fitness differences between the strains, $W_2 > W_1$, are realized and strain-2 can benefit from its advantage over strain-1. Thus, the hologenotypic variation for holobiont selection to operate on is lessened if $d$ is high. High colonization rates homogenize the hologenotypes across the juvenile hosts. Conversely, a low $d$ allows hologenotypic variation to form that is then acted upon by holobiont selection leading to fixation of the selectively favored microbial gene.

**Two-Strain Superiority.** Turn now to the case where the two-strain microbiome is better than either single strain. In Eq. 45, if $W_{12}$ is greater than $W_1$ then the condition for strain-1 to increase when rare is strengthened. Similarly, in Eq, 46, if $W_{12}$ is greater than $W_2$ then the condition for strain-2 to increase when rare is strengthened. Thus two-strain superiority contributes to strain polymorphism beyond that already possible with directional selection.

**Two-Strain Inferiority.** If $W_{12}$ is less than $W_1$ then the condition for strain-1 to increase when rare is weakened. And indeed, if $d\,k\,W_{12} < 1$ then a value of $\hat{g}_2$ may exist which, if exceeded, implies that strain-1 can no longer increase when rare and becomes excluded from the hologenome. Similarly, a value of $\hat{g}_1$ may exist which, if exceeded, implies that strain-2 can no longer increase when rare and becomes excluded from the hologenome. If neither strain can increase when rare, an unstable polymorphism would be indicated, similar to the case of heterozygote inferiority

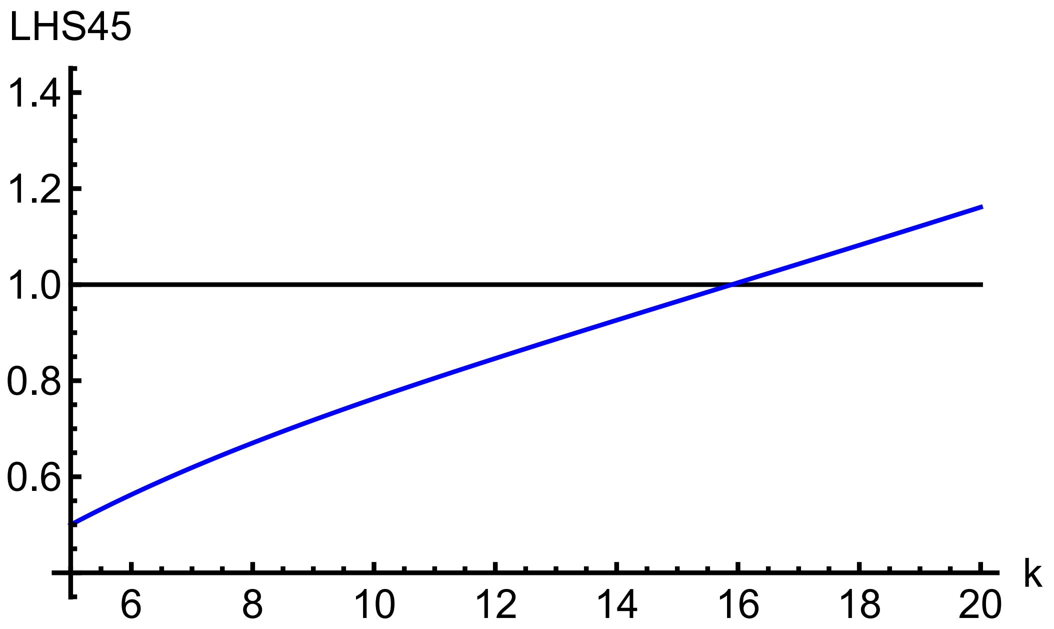


Figure S16: *Left hand side of Eq. 45 as a function of k.* $W_0 = 1$, $W_1 = 1$, $W_{12} = 1/2$, $W_2 = 2$, $d = 0.1$. *According to the figure, strain-1 can increase when rare if* $k > 15.95$ *and cannot increase when rare if* $k < 15.95$.

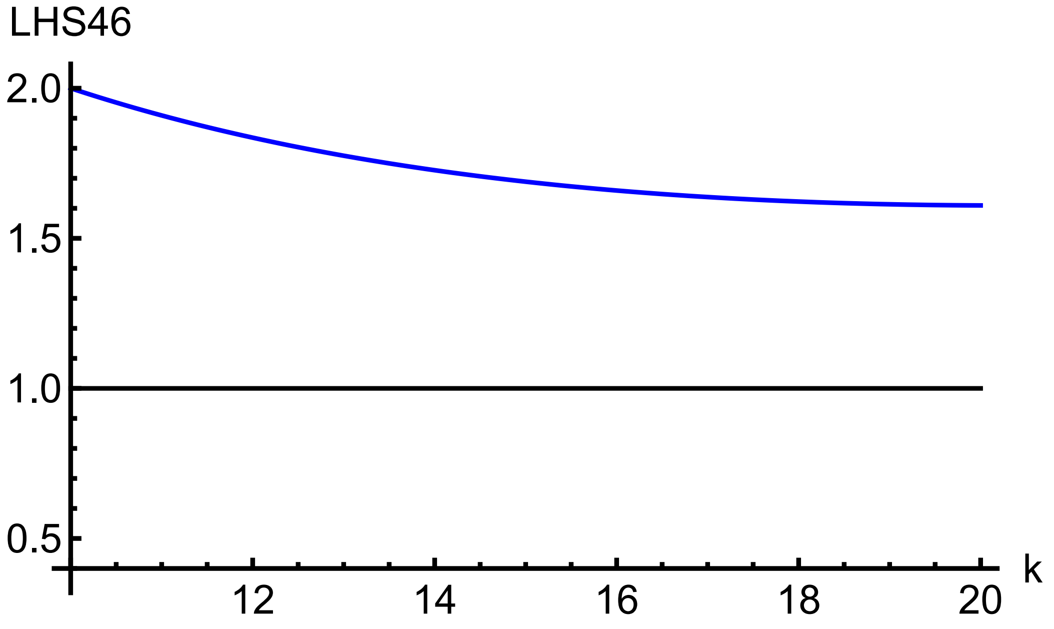


Figure S17: *Left hand side of Eq. 46 as a function of k.* $W_0 = 1$, $W_1 = 1$, $W_{12} = 1/2$, $W_2 = 2$, $d = 0.1$. *According to the figure, strain-2 can always increase when rare.*

in classical population genetics. An unstable equilibrium would represent a "priority effect" in the sense that the first strain to occupy the holobiont population precludes a subsequent strain from entering.[2] However, to date analysis has not uncovered an example of an unstable polymorphism with two-strain inferiority. Instead, in the examples examined so far, the most favored single strain fixes and the least favored single strain is eliminated.

Consider an example of two-strain inferiority where $W_0 = 1$, $W_1 = 1$, $W_{12} = 1/2$, and $W_2 = 2$. These fitness values would lead to an unstable polymorphism with nuclear genes. Figure S16 shows the condition for increase when rare of strain-1 from the left hand side of Eq. 45. When the curve for the left hand side of Eq. 45 is less than one, strain-1 cannot increase when rare and when greater than one strain-1 can increase when rare. The figure presents the left hand side of Eq. 45 as a function of $k$ where $\hat{g_2}(k)$ was obtained by solving Eq. 48 for $\hat{g_2}$ for various values of $k$ between $k_{m2}$ from Eq. 52 and $k_{max}$ which is the root, $k$, of $(d\,k\,W_{12} = 1)$. The left hand side of Eq. 45 crosses the horizontal line at $k \approx 15.95$. Accordingly, strain-1 can increase when rare if $k > 15.95$ and cannot increase when rare if $k < 15.95$.

Figure S17 shows the condition for increase when rare of strain-2 from the left hand side of Eq. 46. When the curve for the left hand side of Eq. 46 is less than one, strain-2 cannot increase when rare and when greater than one strain-2 can increase when rare. The figure presents the left hand side of Eq. 46 as a function of $k$ where $\hat{g_1}(k)$ was obtained by solving Eq. 47 for $\hat{g_1}$ for various values of $k$ between $k_{m1}$ from Eq. 51 and $k_{max}$ which is the root, $k$, of $(d\,k\,W_{12} = 1)$. According to the figure, strain-2 can always increase when rare because the left hand side of Eq 46 is always greater than 1 for $k$ between $k_{m1}$ and $k_{max}$.

According to Figure S16, strain-1 should not be able to increase when rare if $k < 15.95$. Hence, in this case strain-1 is excluded from the holobiont population, whereas strain-2 can increase when rare. Therefore, the hologene pool should fix for strain-2. Figure S18 illustrates this case with $k = 13$.

Also, according to Figure S16, strain-1 should be able to increase when rare if $k > 15.95$. Hence, in this case both strain-1 and strain-2 can increase when rare leading to a polymorphism. However, it is apparent that again the hologene pool fixes for strain-2 and that no polymorphism is indicated. What is happening is that strain-1 does increase when rare but at a low rate leading to its being effectively shed from the holobiont population.

Figure S20 shows the trajectory of $G_1$ from the iterations portrayed in Figures S18 and S19. The figure shows that $G_1$ does not increase when rare if $k = 13$ but does increase when rare if $k = 18$. However, even though $G_1$ does increase when rare if $k = 18$, it only increases from 80 to 1230 during the 20 iterations. During this same period, $G_2$ increases from 20 to $1.38 \times 10^7$,

[2]This priority effect would pertain to the entire holobiont population, not to whether a priority effect occurs between microbe strains within any particular host.

thus dwarfing the increase in $G_1$. Therefore, strain-1 is shed from the holobiont population, even though it can increase when rare. This case is similar to the microbe shedding previously shown in Figures S3 and S8 pertaining to whether a single strain can coexist with its host.

#### Polymorphism between Altruistic and Selfish Microbes

To consider an example where both microbial dynamics and holobiont selection jointly occur, suppose that strain-1 always excludes strain-2 in a host where both are present, as previously considered in Figs. S11 and S12, but now also allow directional holobiont selection in favor of strain-2. That is, strain-2 (the altruistic microbe) sacrifices its competitive ability with respect to strain-1 (the selfish microbe), but receives a higher fitness at the holobiont level. Can holobiont selection favoring strain-2 keep it from going extinct? Specifically, if $k_2 < k_{2c}$ from Eq. 41 such that strain-2 is excluded in the absence of holobiont selection, can directional holobiont selection in favor of strain-2 retain it in the holobiont population. The directional selection favoring strain-2 is implemented by assuming $W_2 > W_{12} = W_1 = W_0$.

Eq. 30 is again

$$W_o(g_{1o}, g_{2o}) = h_{00o}\, W_0 + h_{10o}\, W_1 + h_{02o}\, W_2 + h_{12o}\, W_{12}$$

where

$$\begin{aligned} h_{00o} &= (1 - P_{1o})\,(1 - P_{2o}) \\ h_{10o} &= P_{1o}\,(1 - P_{2o}) \\ h_{02o} &= (1 - P_{1o})\,P_{2o} \\ h_{12o} &= P_{1o}\,P_{2o} \end{aligned}$$

and

$$\begin{aligned} P_{1o} &= 1 - e^{-d_1\, g_{1o}} \\ P_{2o} &= 1 - e^{-d_2\, g_{2o}} \end{aligned}$$

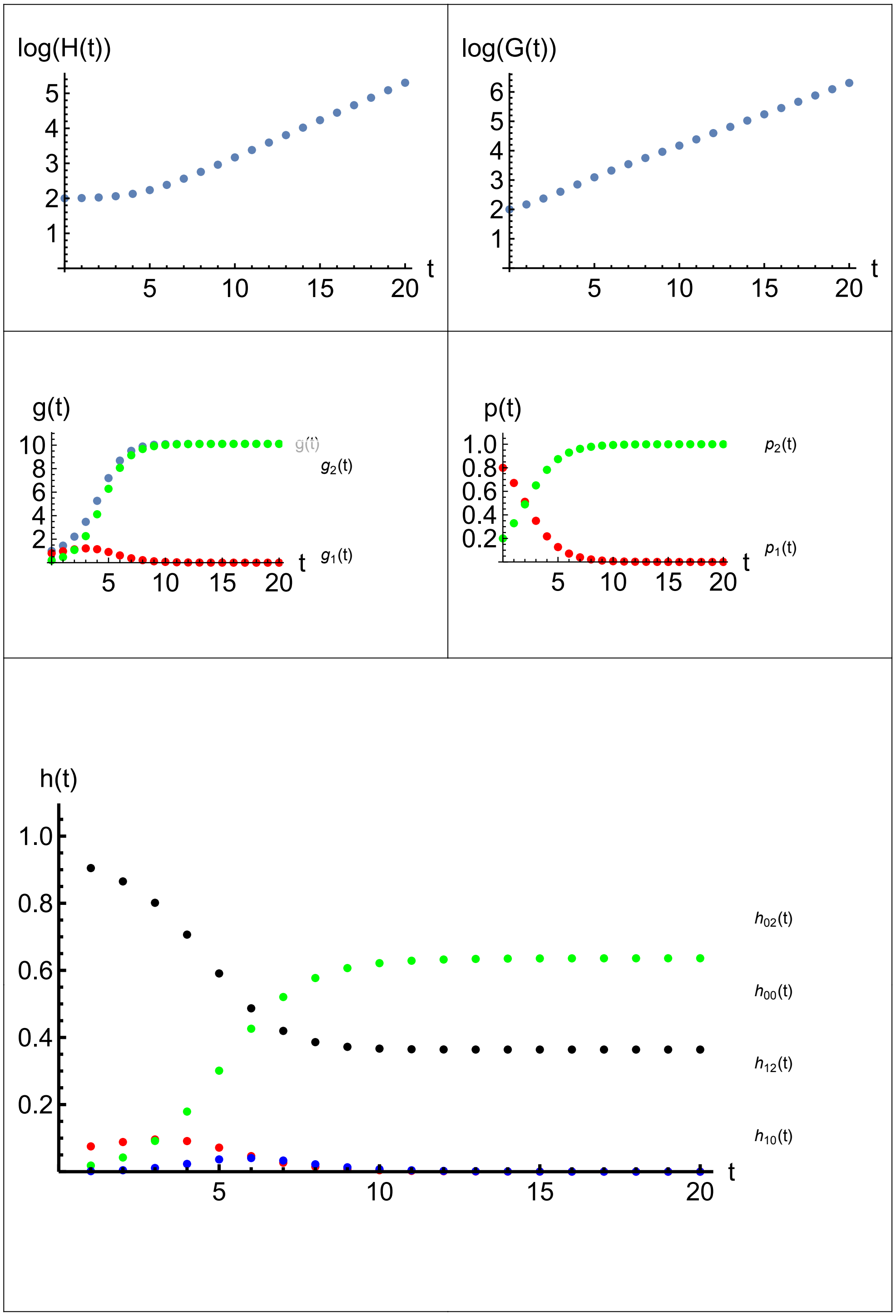


Figure S18: *Two-strain Inferiority—strain-1 cannot increase when rare.* $W_0 = 1$, $W_1 = 1$, $W_{12} = 1/2$, $W_2 = 2$. *Also,* $d_1 = 0.1$, $d_2 = 0.1$, $k_1 = 13$, $k_2 = 13$, $a_{12} = 0$, $a_{21} = 0$, $H_{\text{init}} = 100$, $G_{1\text{init}} = 20$, $G_{2\text{init}} = 80$, $t_{\max} = 20$. *Asymptotically,* $R_{oH} \approx 1.64$, $R_{oG} \approx 1.64$, $g_o = 10.10$, $g_{1o} = 0.00$, $g_{2o} = 10.11$, $p_{1o} = 0.00$, $p_{2o} = 1.00$, $h_{00o} = 0.364$, $h_{10o} = 0.000$, $h_{02o} = 0.636$, *and* $h_{12o} = 0.000$ *where* $R_{oH} = H(t_{\max})/H(t_{\max} - 1)$ *and* $R_{oG} = G(t_{\max})/G(t_{\max} - 1)$.

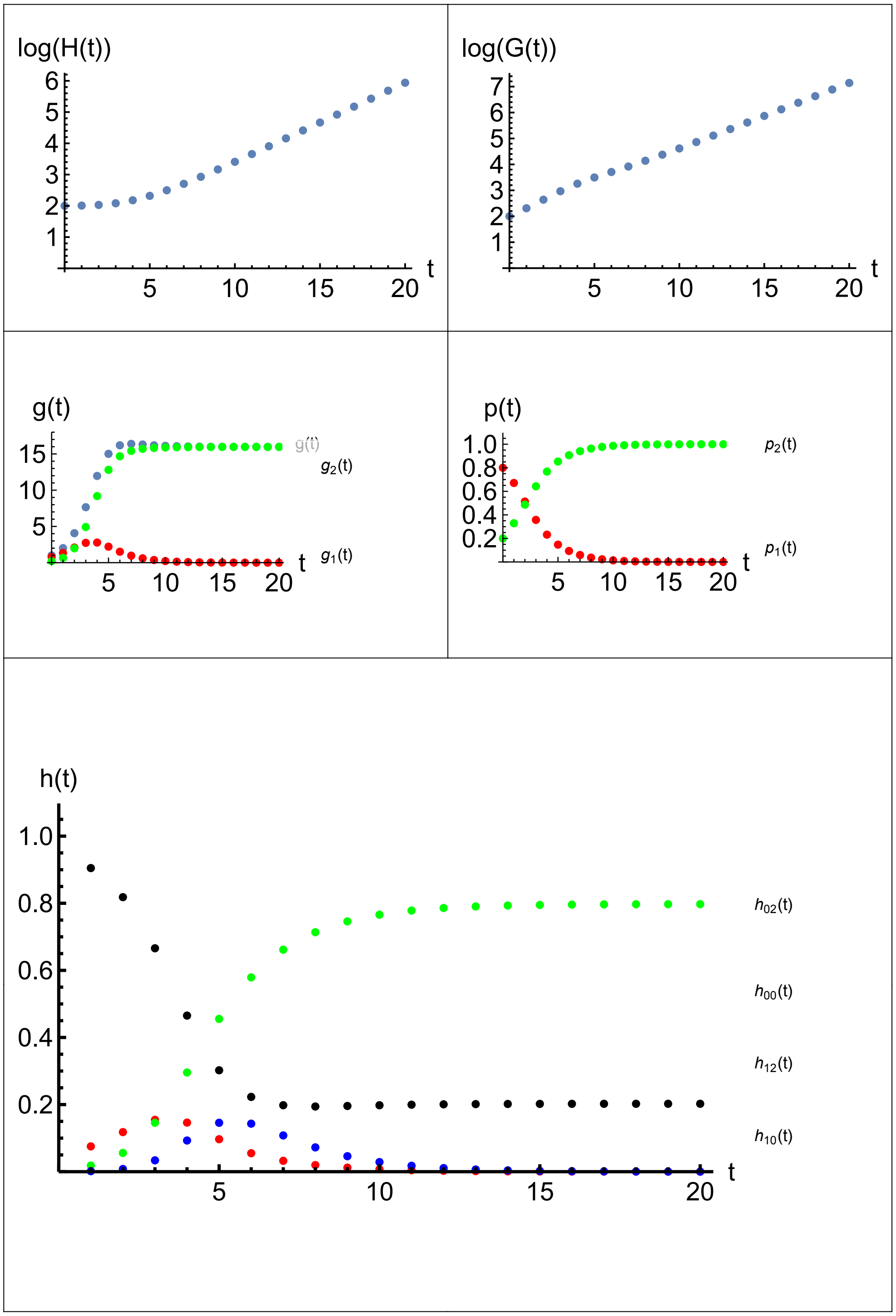


Figure S19: *Two-strain Inferiority—strain-1 is shed.* $W_0 = 1$, $W_1 = 1$, $W_{12} = 1/2$, $W_2 = 2$. *Also,* $d_1 = 0.1$, $d_2 = 0.1$, $k_1 = 18$, $k_2 = 18$, $a_{12} = 0$, $a_{21} = 0$, $H_{\text{init}} = 100$, $G_{1\text{init}} = 20$, $G_{2\text{init}} = 80$, $t_{\max} = 20$. *Asymptotically,* $R_{oH} \approx 1.80$, $R_{oG} \approx 1.80$, $g_o = 15.97$, $g_{1o} = 0.00$, $g_{2o} = 15.97$, $p_{1o} = 0.000$, $p_{2o} = 1.000$, $h_{00o} = 0.202$, $h_{10o} = 0.000$, $h_{02o} = 0.797$, *and* $h_{12o} = 0.000$ *where* $R_{oH} = H(t_{\max})/H(t_{\max} - 1)$ *and* $R_{oG} = G(t_{\max})/G(t_{\max} - 1)$.

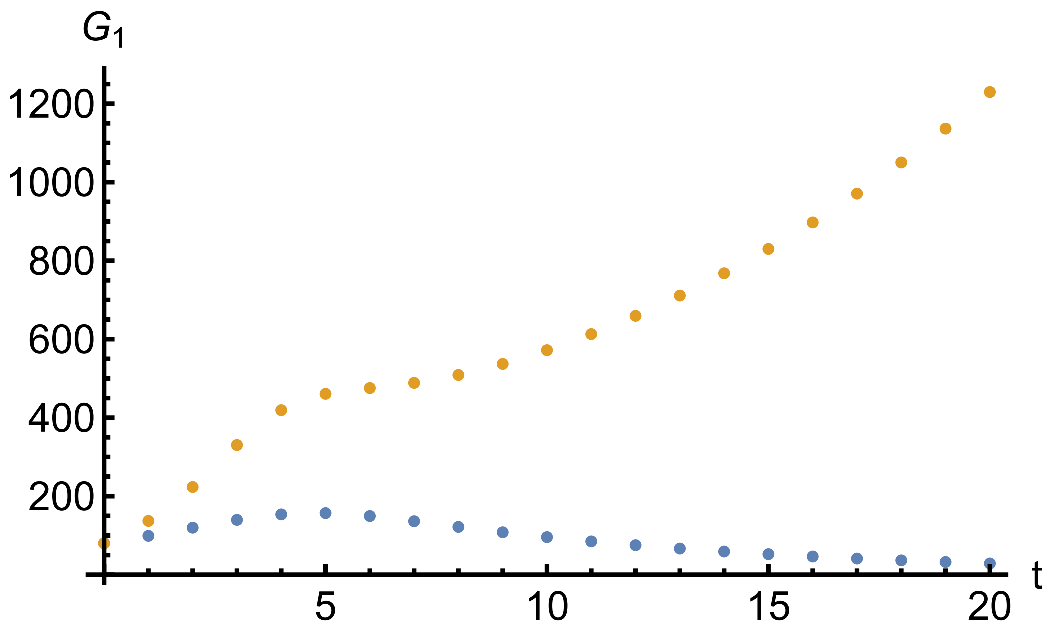


Figure S20: *Trajectory of $G_1$ from the iterations portrayed in Figures S18 (bottom) and 19 (top). $G_1$ does not increase when rare if $k = 13$ (bottom), but does increase when rare if $k = 18$ (top).*

Also, if strain-1 always eliminates strain-2 when both colonize the same host, then $n_1 = k_1$ and $n_2 = 0$. Hence

$$\begin{aligned} w_{1,1} &= k_1 W_1 \\ w_{2,2} &= k_2 W_2 \\ w_{1,12} &= k_1 W_1 \\ w_{2,12} &= 0 \end{aligned}$$

Therefore

$$\begin{aligned} \overline{w_{1o}} &= k_1 W_1 \\ \overline{w_{2o}} &= (1 - P_{1o}) k_2 W_2 \end{aligned}$$

and the equilibrium conditions for strain-1 and strain-2, Eqs. 33 and 34, reduce to

$$g_{1o} = \frac{P_{1o} k_1 W_1}{W_o(g_{1o}, g_{2o})} \quad (53)$$

$$g_{2o} = \frac{P_{2o} (1 - P_{1o}) k_2 W_2}{W_o(g_{1o}, g_{2o})} \quad (54)$$

Figure S21 presents an illustration of the roots of Eqs. 53 and 54, $g_{1o}$ and $g_{2o}$, as a function of $W_2$. The selfish strain-1 always exists in the holobiont population for the illustrated values of

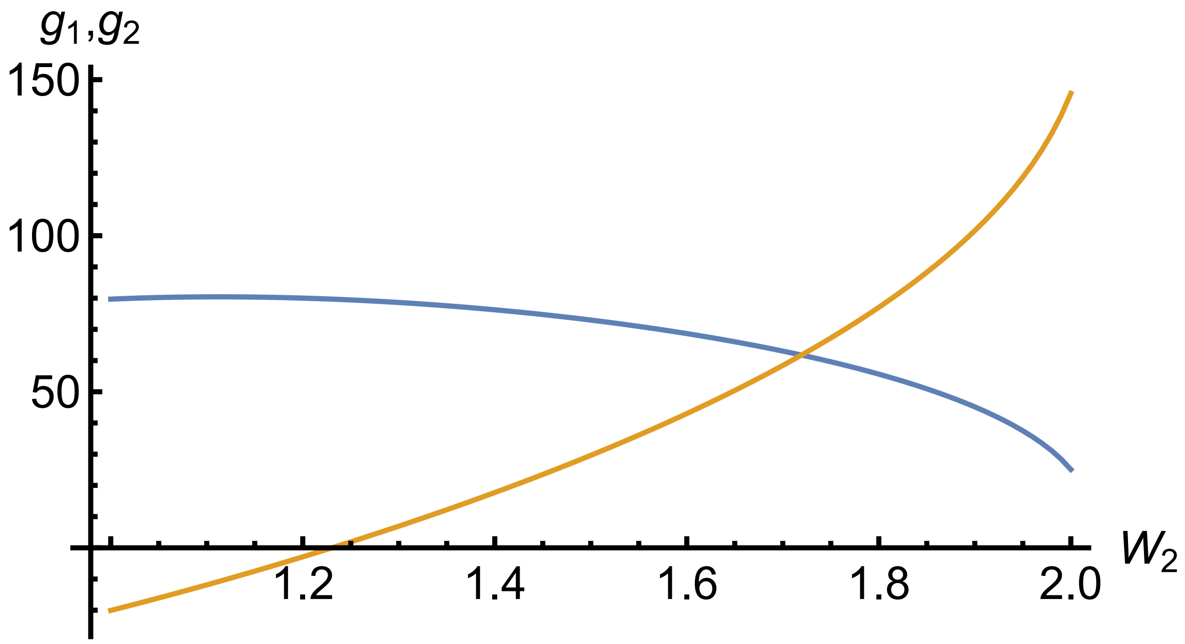


Figure S21: *Equilibrium relative abundances of strain-1 (selfish) and strain-2 (altruistic), $g_{1o}$ and $g_{2o}$, as a function of $W_2$, the holobiont selection in favor of strain-2. Strain-1 is the upper curve that decreases with $W_2$—this strain always exists in the holobiont population. Strain-2 is the lower curve that increases with $W_2$—this strain can enter the holobiont population and coexist with strain-1 if $W_2$ is greater than $\approx 1.23$. The other parameters are $d_1 = 0.02$, $d_2 = 0.02$, $k_1 = 100$, $k_2 = 200$, $W_0 = 1$, $W_1 = 1$, and $W_{12} = 1$. Also, $a_{12} < 1/2$ and $a_{21} > 2$ to indicate the competitive asymmetry that causes strain-1 to exclude strain-2 in the holobionts that are colonized by both strains.*

$W_2$. According to the figure, the altruistic strain-2 can enter the holobiont population and coexist with strain-1 if $W_2$ is greater than $\approx 1.23$.

Figure S22 presents trajectories illustrating the coexistence of altruistic and selfish microbes. Here, $W_2 = 1.5$, which is above the value of 1.23 that Figure S21 indicated as the minimum $W_2$ permitting coexistence. Thus, if the benefit a microbe confers on holobiont fitness is large enough, it persists in the holobiont population despite its otherwise being eliminated by competition with an alternative microbe.

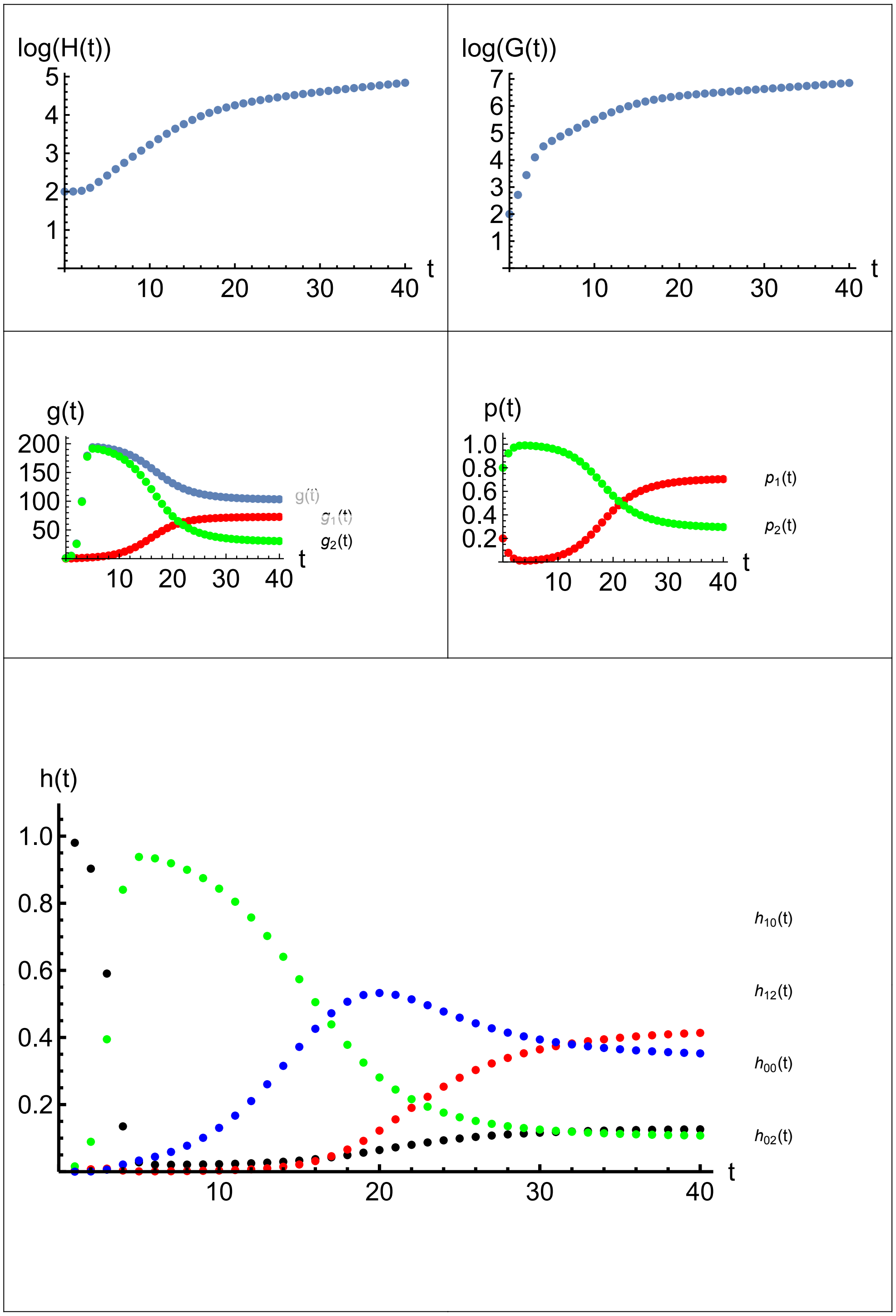


Figure S22: *Coexistence of altruistic and selfish microbes. Strain-1 is selfish and strain-2 is altruistic.* $d_1 = 0.02$, $d_2 = 0.02$, $k_1 = 100$, $k_2 = 200$, $W_0 = 1$, $W_1 = 1$, $W_{12} = 1$ *and* $W_2 = 1.5$. *Also,* $a_{12} < 1/2$ *and* $a_{21} > 2$. $H_{\text{init}} = 100$, $G_{1\text{init}} = 20$, $G_{2\text{init}} = 80$, $t_{\max} = 40$. *Asymptotically,* $R_{oH} \approx 1.05$, $R_{oG} \approx 1.05$, $g_o = 103.33$, $g_{1o} = 72.70$, $g_{2o} = 30.63$, $p_{1o} = 0.704$, $p_{2o} = 0.296$, $h_{00o} = 0.126$, $h_{10o} = 0.414$, $h_{02o} = 0.108$, *and* $h_{12o} = 0.352$ *where* $R_{oH} = H(t_{\max})/H(t_{\max} - 1)$ *and* $R_{oG} = G(t_{\max})/G(t_{\max} - 1)$.

#### Microbiome-Host Integration

**Optimal Microbial Altruism.** The next paragraphs investigate an optimal tradeoff by a microbe between its within-host carrying capacity, $k$, and the between-host fitness of the holobiont it inhabits, $W$. The optimal tradeoff is expressed in terms of a phenotypic variable, $x$, that is a measure of the "altruistic effort" by the microbe. The altruistic effort might refer to the amount of a valuable metabolite transferred from the microbe to the host. The $x$ measures how much of a microbe's $k$ it foregoes to improve $W$. The optimal altruistic effort by the microbe is assumed here to be the $x$ that maximizes its multilevel fitness $w$.

Next, consider an explicit form for the phenotypic tradeoffs.

$$\begin{aligned} k(x) &= k_o - c\,x \ \text{ for } \ x \leq x_{\text{afford}} \\ &= 0 \ \text{ for } \ x > x_{\text{afford}} \end{aligned} \tag{55}$$

where the maximum altruism the microbe can afford while maintaining a positive $k_o$ is

$$x_{\text{afford}} = k_o/c \tag{56}$$

and $c$ is the cost per unit effort paid by the microbe for the altruistic effort it supplies to the host and $k_o$ is the baseline $k$ without any altruistic effort.

Similarly, suppose

$$W(x) = W_o + b\,x \tag{57}$$

where $b$ is the benefit per unit effort received by the host from the altruistic effort supplied by the microbe and $W_o$ is the baseline host fitness without any altruism from the microbe. The multilevel fitness, $w(x) = k(x)\,W(x)$, is therefore a quadratic function of $x$ and its maximum occurs at

$$\hat{x} = \frac{b\,k_o - c\,W_o}{2\,b\,c} \tag{58}$$

For a positive $\hat{x}$, the benefit to cost ratio must satisfy

$$\frac{b}{c} > \frac{W_o}{k_o} \tag{59}$$

For a given $x$, the microbe can exist with the host at a relative abundance, $\hat{g}(x)$ from Eq. 11, as the root of

$$\hat{g}(x) = \frac{k(x)\,W(x)\,(1 - e^{-d\,\hat{g}(x)})}{W_o\,e^{-d\,\hat{g}(x)} + W(x)\,(1 - e^{-d\,\hat{g}(x)})} \tag{60}$$

where the fitness of hosts without any microbes is assumed to be $W_o$ and the fitness of hosts with the microbes is $W(x)$. The $\hat{g}(x)$ is feasible, according to Eq. 15, provided $k(x)$ is greater than $k_m(x)$ where

$$k_m(x) \equiv \frac{W_o}{d\ W(x)} \tag{61}$$

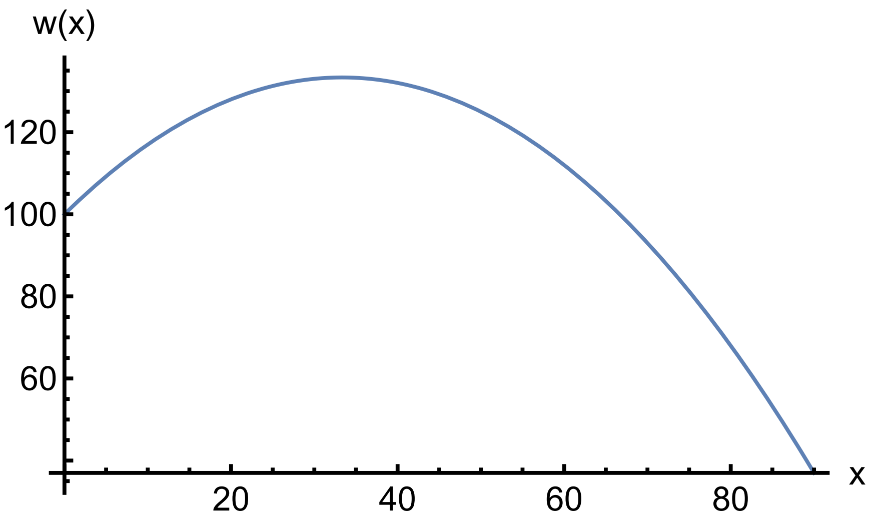


Figure S23: *Multilevel microbial fitness, w, as a function of the altruistic effort, x. The optimal effort,* $\hat{x} = 33.333$. *The maximum altruism consistent with microbe host coexistence,* $x_{max} = 90$ *and the maximum altruism the microbe can afford is* $x_{\text{afford}} = 100$. *Here,* $d = 0.1$, $k_o = 100$, $W_o = 1$, $b = 0.03$, $c = 1$.

If $x = 0$, then $W(x) = W_o$. Therefore, $k_o$, must be greater than $1/d$ because $k(x)$ declines with $x$. Therefore, the maximum $x$ allowed is $x_{max}$ which is the root of

$$\frac{1}{d} = k(x_{max}) \tag{62}$$

That is, if the microbe contributes more altruistic effort than $x_{max}$, then the microbe can no longer coexist with the host. So, the optimal altruistic effort, $\hat{x}$, must lie in the interval $[0, x_{max}]$. The $x_{max}$ works out to be

$$x_{max} = \frac{d\,k_o - 1}{c\,d} = k_o/c - 1/d \tag{63}$$

The $x_{max}$ is less than $x_{\text{afford}}$ as required. Thus, the microbe could afford a higher $x$ than $x_{max}$ without incurring a negative $k_o$, but doing so would cause its not being able to coexist with the host. In summary, $b/c$ must be large enough according to Eq. 58 for a positive $\hat{x}$ to exist, provided $k_o > 1/d$. The $\hat{x}$ must be less than $x_{max}$ from Eq. 62 for the microbe to coexist with its host. An example is provided in Figure S23.

**Selfish Invades.** Once an optimal altruistic effort, $\hat{x}$ exists, the further question arises of whether microbes with the optimal effort exclude all other microbe strains with a non-optimal effort. Suppose a mutant strain arises that is identical in all respects to the optimal strain except for expending a different altruistic effort, $y \neq \hat{x}$. By this assumption, the competition coefficients, $a_{12}$ and $a_{21}$ both equal 1 and the other parameters, $d$, $W_o$, $k_o$, $b$ and $c$ of the altruistic and mutant strains are the same for both. All that differs between the strains is their altruistic efforts, $x$ and $y$. Can the optimal altruistic strain exclude all non-optimal altruistic strains?

Indeed, consider the more general question of whether a selfish mutant can invade *any* established altruistic strain, including an optimally altruistic strain. Let the established strain exhibit some degree of altruism, not necessarily optimal, at $x > 0$. Can a completely selfish mutant with $y = 0$ invade any altruistic strain with $x > 0$?

According to Eq. 36, a mutant strain, when rare, cannot enter the holobiont population in which the altruistic strain is already established at $g = \hat{g}(x)$ provided

$$d \left( k(y)\, W(y)\, e^{-d\,\hat{g}(x)} + n_m\, W_{am}\, (1 - e^{-d\,\hat{g}(x)} \right) < 1 \tag{64}$$

where $n_m$ is abundance of the mutant strain in a dual-strain microbiome and $W_{am}$ is the host fitness with a dual-strain microbiome containing the altruist and the selfish mutant. And considering the optimal altruist in particular, if Eq. 63 were satisfied for any $y \neq \hat{x}$, then the $\hat{x}$ would represent a microbial counterpart to an evolutionarily stable strategy, which for nuclear genes, is a strategy that cannot be invaded by any other strategy.

If $y < x$, *i.e.*, the mutant supplies less than the existing altruistic effort, then $k(y) > k(x)$. In this situation, the two strains cannot coexist together within the hosts they both colonize given that the reciprocal competition coefficients equal 1. Hence the altruistic strain is excluded from all such hosts because of having the lower $k$. Accordingly, $W_{am}$ is the same as $W(y)$ and $n_m$ is the same as $k(y)$. Thus, Eq. 63 boils down to

$$d\, k(y)\, W(y) < 1 \tag{65}$$

Substituting the phenotypic tradeoffs, Eqs. 55 and 56, yields

$$d\,(k_o - y)\,(W_o + b\,y) < 1 \tag{66}$$

And for a completely selfish mutant, $y = 0$, Eq. 65 becomes

$$d\, k_o\, W_o < 1 \tag{67}$$

Because $k_o > 1/d$ according to Eq. 60, and also $W_o \geq 1$, Eq. 66 is never satisfied. Therefore, the completely selfish mutant with $y = 0$ *always* increases when rare into a holobiont population fixed for a strain supplying *any* altruistic effort, including $x = \hat{x}$. Hence, $\hat{x}$ is not the counterpart of an evolutionarily stable strategy because it can always be invaded by a more selfish strategy. Instead, only the completely selfish microbe, $x = 0$, is an evolutionarily stable strategy. Figure S24 illustrates a selfish microbe with $y = 0$ excluding an altruistic microbe with $x = \hat{x}$.

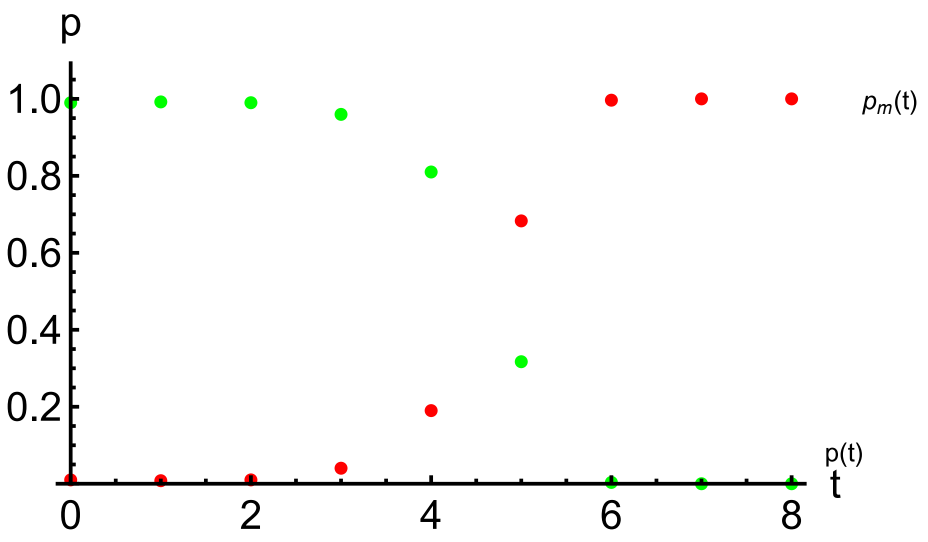


Figure S24: *Frequency of altruistic microbe, $p$, and selfish mutant, $p_m$, through time without host intervention. Host fitness with altruistic microbe, $W = W_o + b\,x$. Altruistic microbe carrying capacity, $k = k_o - c\,x$, Host fitness with selfish mutant, $W_m = W_o + b\,y$, Selfish microbe carrying capacity, $k_m = k_o - c\,y$. Here, $d = 0.1$, $W_o = 1$, $k_o = 100$, $b = 0.03$, $c = 1$, $x = 33.3333$, $y = 0$, $H_{init} = 100$, $G_{a,init} = 99$, $G_{m,init} = 1, t_{max} = 8$.*

**Host Intervention.** To obtain some degree of evolutionarily stable altruistic effort by the microbes to the host, host intervention is needed to discriminate against microbial selfishness. This intervention can be achieved by assuming $d_m \neq d$ and allowing the host to lower the colonization coefficient, $d_m$, for the mutant strain relative to $d$. In all other respects, the altruistic and selfish strains are the same—they solely differ in their degrees of altruism and in their colonization coefficients.

Suppose the mutant, $y$, is more selfish than the existing altruist, $x \leq \hat{x}$. Can the selfish mutant invade? In this situation, the condition for the selfish mutant, $y$, to be excluded, Eq. 65, becomes

$$d_m\,(k_o - c\,y)\,(W_o + b\,y) < 1 \tag{68}$$

If $y = 0$, the Eq. 67 is

$$d_m\,k_o\,W_o < 1 \tag{69}$$

which is satisfied provided

$$d_m < k_o\,W_o \tag{70}$$

So the host must reduce the colonization parameter of the mutant low enough to satisfy Eq. 69 to prevent a completely selfish mutant from invading. Furthermore, suppose $d_m$ is such that the

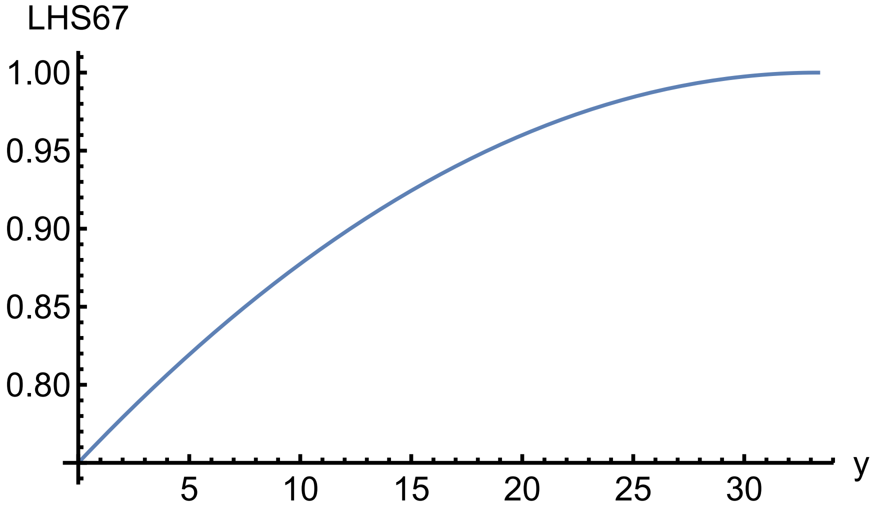


Figure S25: *Left hand side of Eq. 67, as a function of the mutant's altruistic effort,* $y < \hat{x}$. *The selfish microbe cannot invade if LHS67 is less than one. The optimal effort,* $\hat{x} = 33.333$. *Here,* $d_m = \hat{d}_m(\hat{x}) = 0.0075$, $d = 0.1$, $k_o = 100$, $W_o = 1$, $b = 0.03$, $c = 1$.

LHS of Eq. 67 equals 1 when $y$ equals $x \leq \hat{x}$. That value of $d_m$ is

$$\hat{d}_m = \frac{1}{(k_o - c\,x)\,(W_o + b\,x)} \tag{71}$$

The derivative of the LHS of Eq. 67 with $\hat{d}_m$ is

$$\hat{d}_m\,(b\,k_o - c\,W_o) \tag{72}$$

This derivative is positive if

$$\frac{b}{c} > \frac{W_o}{k_o} \tag{73}$$

which is identical to the condition, Eq, 58, for an optimal degree of altruism, $\hat{x}$, to be positive. Figure S25 shows the LHS of Eq. 67 as a function of the mutant's altruistic effort, $y$, for $y$ between 0 and $\hat{x}$, for $d_m = \hat{d}_m(\hat{x})$. The figures shows that the LHS of Eq. 67 is less than 1 for $y < \hat{x}$, implying that any mutant whose altruistic effort is less than $\hat{x}$ cannot invade.

Thus, for any $x \leq \hat{x}$, if the host lowers $d$ to $\hat{d}_m(x)$ then that altruistic effort, $x$, is stable to invasion by any strain with a lower altruistic effort, $y < x$, including an altruistic effort of 0. And in particular, if $x$ happens to equal $\hat{x}$, then the optimal altruistic effort is stable to invasion by any strain with a lower altruistic effort. However, the host does not "know" the optimal altruistic effort from the microbe's point of view. It would be a coincidence if $d_m$ happened to stabilize the microbe's optimal altruistic effort rather than some lesser effort. Nonetheless, the introduction of host intervention guarantees that at least some altruism by the microbes occurs. Figure S26 illustrates the altruistic strain excluding the selfish strain as a result of host intervention.

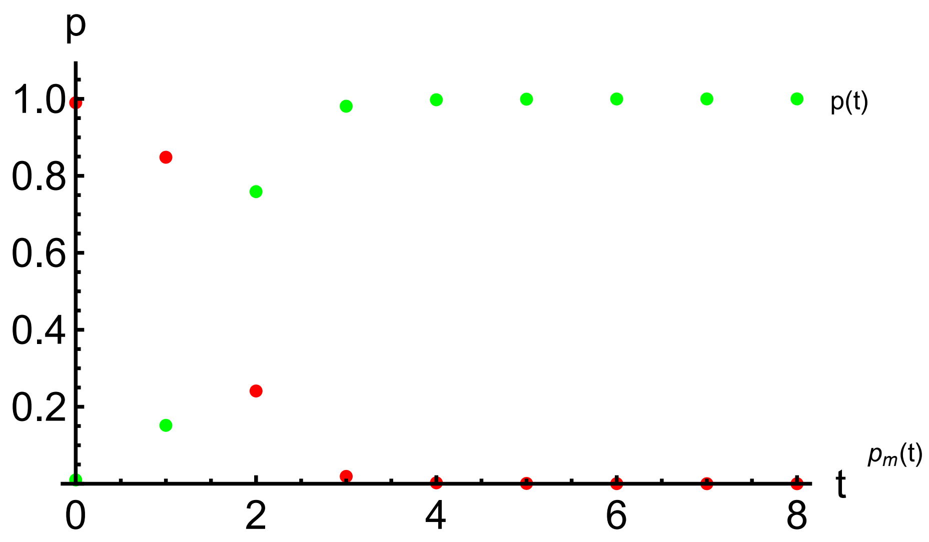


Figure S26: *Frequency of altruistic microbe, p and selfish mutant, $p_m$, through time with host intervention. Host fitness with altruistic microbe, $W = W_o + b\,x$. Altruistic microbe carrying capacity, $k = k_o - c\,x$. Host fitness with selfish microbe, $W_m = W_o + b\,y$. Selfish microbe carrying capacity, $k_m = k_o - c\,y$. Here, $d_m = 0.0075$, $d = 0.1$, $W_o = 1$, $k_o = 100$, $b = 0.03$, $c = 1$, $x = 33.3333$, $y = 0$, $H_{init} = 100$, $G_{a,init} = 1$, $G_{m,init} = 99$, $t_{max} = 8$.*

**Microbiome-Host Game.** How, and to what end would a host intervene in the colonization parameter of a microbial strain in its microbiome? One way the host might influence the colonization parameters of its microbial strains results with its immune system. Suppose that a host initially "sees" a microbe that colonizes it as a deleterious invader. If so, it creates antibodies to the microbe which has the effect of lowering the microbe's colonization parameter. That is, if a microbe is deactivated by a host's immune system upon entry into the host, in effect the colonization does not occur. The colonization parameter, $d$, refers to the net colonization after the action of the host's immune system is taken into account. So the question now becomes how much the host's immune system is expected to deactivate the entering microbes as a function of how altruistic the microbe is.

**Host Antibodies.** Suppose the number of antibodies a host makes toward a microbe results from the benefit and cost of antibody production. Let $z$ be the quantity of antibodies that the host makes in response to a microbe. Suppose the fitness benefit to antibody production, $z$, shows a decreasing return to scale,

$$B(z) = \alpha\,(1 - e^{-\beta z}) \tag{74}$$

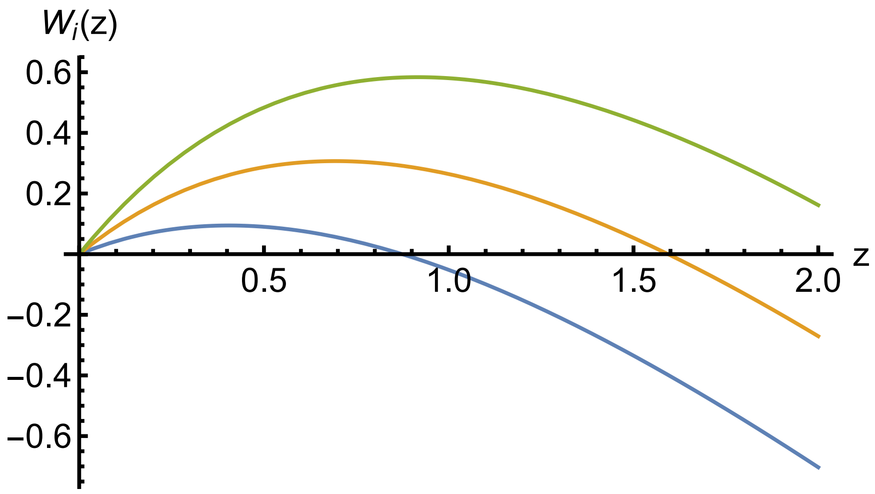


Figure S27: *Net fitness benefit, $W_i(z)$, to host as a function of antibody production, $z$. Curves from left to right are for $\alpha = 1.5$, 2.0, and 2.5 respectively. Here, $\beta = 1$ and $\gamma = 1$.*

where $\alpha$ is the amplitude of the benefit and $\beta$ measures the fall off in benefit as the number of antibodies becomes large. Suppose the fitness cost of antibody production is linear in $z$

$$C(z) = \gamma z \tag{75}$$

where $\gamma$ is the cost per unit antibody. So the net fitness benefit from the immune system of antibody production, $W_i(z)$, is

$$W_i(z) = B(z) - C(z) = \alpha\,(1 - e^{-\beta z}) - \gamma z \tag{76}$$

Figure S27 illustrates several curves of $W_i(z)$ for various values of the benefit amplitude, $\alpha$. The amount of antibody production, $\hat{z}$, that maximizes the net fitness benefit, $W_i(z)$ from Eq. 75 is

$$\hat{z} = \frac{\ln(\frac{\alpha\beta}{\gamma})}{\beta} \tag{77}$$

The $\hat{z}$, is an increasing function of the benefit amplitude, $\alpha$.

Now, suppose further that the benefit amplitude for antibody production depends on the altruism being supplied by the microbe. After all, the benefit to the host of making antibodies to a microbe is lessened if the microbe is supplying some nutrients to the host. Accordingly, suppose that

$$\begin{aligned} \alpha(x) &= \alpha_o - \mu x \;\text{ for }\; x \le x_{\alpha 0} \\ &= 0 \;\text{ for }\; x > x_{\alpha 0} \end{aligned} \tag{78}$$

where

$$x_{\alpha 0} = \alpha_o / \mu \tag{79}$$

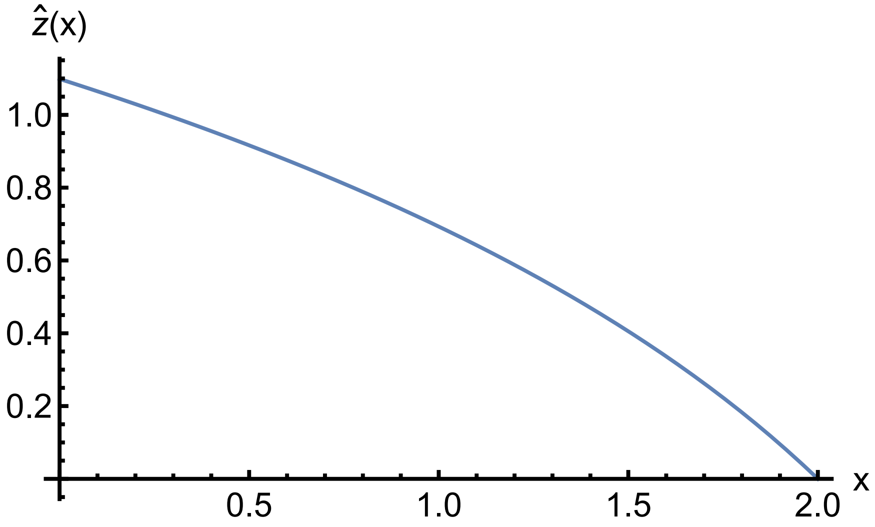


Figure S28: *Optimal antibody production, $\hat{z}(x)$, as a function of the altruism, $x$, being supplied by the microbe to the host. Here, $\hat{x}_{max} = 2$, $\hat{z}_{max} = 1.10$, $\alpha_o = 3$, $\mu = 1$, $\beta = 1$ and $\gamma = 1$.*

is the amount of microbial altruism that makes the benefit to the host of any antibody production equal 0, not taking into account the cost of antibody production. Then, taking costs into account, the optimal antibody production $\hat{z}$ from Eq. 76 becomes a function of the microbe's degree of altruism, $x$,

$$\begin{aligned} \hat{z}(x) &= \frac{\ln(\frac{(\alpha_o - \mu\, x)\,\beta}{\gamma})}{\beta} \;\; x \leq \hat{x}_{max} \\ &= 0 \;\text{ for }\; x > \hat{x}_{max} \end{aligned} \tag{80}$$

where

$$\hat{x}_{max} = \frac{\alpha_o - \gamma/\beta}{\mu} \tag{81}$$

is obtained by setting the argument of ln() equal to 1 in Eq, 79 and the solving for $x$. The $\hat{x}_{max} < x_{\alpha 0}$ as required. That is, if the microbe is supplying a degree of altruism greater than $\hat{x}_{max}$, then the host does not make any antibodies at all. And if $0 < x < \hat{x}_{max}$ then the host makes $\hat{z}(x)$ amount of antibodies depending on how much altruism, $x$, is being supplied. Natural selection on host nuclear genes maximizes $W_i$ leading to the evolution of the optimal immune response, $\hat{z}(x)$, to a microbe supplying $x$ nutrients to it. An illustration appears in Figure S28.

Furthermore, if the microbe is completely selfish, $x = 0$, then the antibody response is

$$\hat{z}_{max} = \frac{\ln(\frac{\alpha_o\,\beta}{\gamma})}{\beta} \tag{82}$$

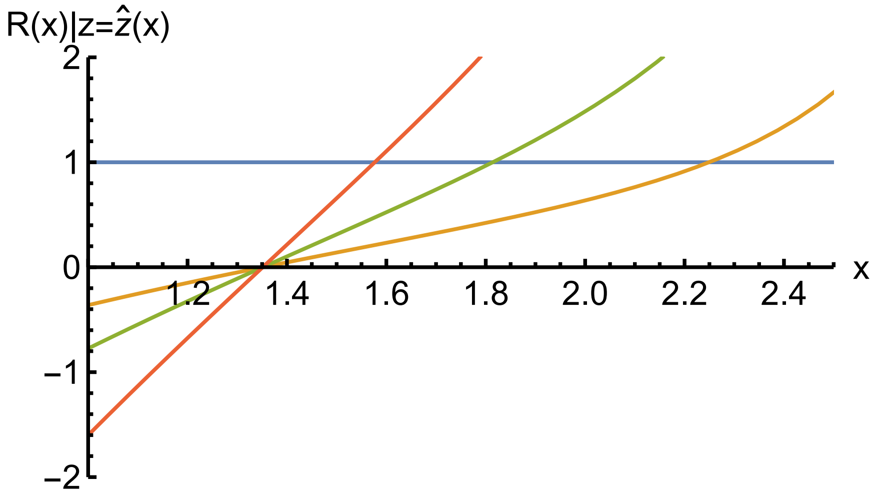


Figure S29: *Microbial increase-when-rare condition,* $R(x)|_{z=\hat{z}(x)}$*, as a function of the altruism,* $x$*, being supplied by the microbe to the host taking into account the host's antibody response,* $\hat{z}(x)$*. From left to right, curves crossing the line at* $r = 1$ *are for* $k_o = 32$*,* $k_o = 16$ *and* $k_o = 8$*. Here,* $\hat{x}_{max} = 2$*,* $\hat{z}_{max} = 1.10$*,* $\alpha_o = 3$*,* $\mu = 1$*,* $\beta = 1$*,* $\gamma = 1$*,* $d_o = 0.1$*,* $\nu = 0.2$*,* $b = 0.03$*,* $c = 1$*, and* $W_o = 1$*.*

**Microbial Altruism.** A side effect of the host's antibody production is to lower the microbe's colonization parameter

$$d(z) = d_o - \nu z \text{ for } z \leq z_{max} \tag{83}$$

$$= 0 \text{ for } z > z_{max} \tag{84}$$

where

$$z_{max} = d_o/\nu \tag{85}$$

is the maximum amount of host antibody that allows a positive colonization microbe colonization parameter. In view of Eq. 81, if

$$\hat{z}_{max} > z_{max} \tag{86}$$

then the host responds to a completely selfish microbe $x = 0$, with enough antibodies to exclude it.

The minimum amount of altruism the microbe must supply to be allowed to colonize is found as the value of $x$ for which the microbe can increase when rare. From Eq. 13 (or Eq. 12 with $W_0 = 1$), the microbe can increase when rare if $R$ given by

$$R \equiv d\,k\,W_1 > 1 \tag{87}$$

The $R$, upon substituting expressions for the phenotypic tradeoffs, Eqs. 55, 56, 73, 74, 77, and 82, is

$$R(x)|_{z=\hat{z}(x)} = (d_o - \nu z)\,(k_o - c\,x)\,(W_o + b\,x + (\alpha_o - \mu\,x)\,(1 - e^{-\beta z}) - \gamma z) \tag{88}$$

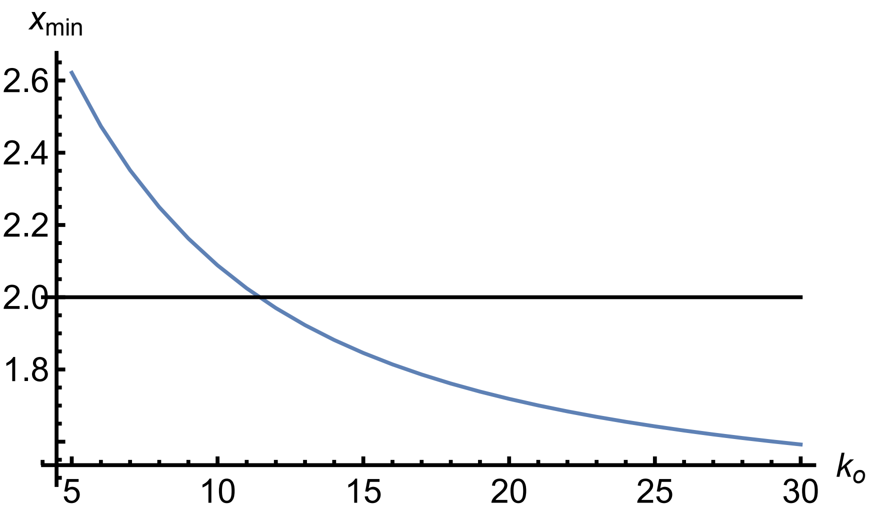


Figure S30: *Minimum altruism, $x_{min}(k_o)$, the microbe must supply to the host to increase when rare as a function of its within-host base carrying capacity, $k_o$. The horizontal line is $\hat{x}_{max}$, the degree of altruism that causes the host not make any antibodies at all. The intersection of the $x_{min}(k_o)$ curve with the $\hat{x}_{max}$ line occurs at $\tilde{k}_o = 11.43$. The $\tilde{k}_o$ marks the cutoff between insufficiently altruistic strains excluded from the holobiont and the altruistic strains capable of joining. Here, $\hat{x}_{max} = 2$, $\alpha_o = 3$, $\mu = 1$, $\beta = 1$, $\gamma = 1$, $d_o = 0.1$, $\nu = 0.2$, $b = 0.03$, $c = 1$, and $W_o = 1$.*

where $z = \hat{z}(x)$ from Eq. 79. The $R(x)$ is an increasing function of $x$ between 0 and $x_{max}$. Thus, increasing the altruism beyond that just needed to enter the microbiome raises the speed at which the microbe can increase when rare because the loss of the microbe's multilevel fitness from contributing to the host is more than compensated by the host's lowering antibody production which in turn increases the microbe's colonization parameter.

Figure S29 shows curves of $R(x)|_{z=\hat{z}(x)}$ for three values of $k_o$ near the place where the curves intersect 1. For each $k_o$, the intersection point is the minimum $x$ that the microbe must supply, $x_{min}$, to keep the host antibody response low enough for the colonization parameter to support the microbe's increasing when rare. Using *Mathematica*'s FindRoot[] function with $R(x)|_{z=\hat{z}(x)} = 1$, yields Figure S30 that shows $x_{min}$ as a function of $k_o$. The figure also shows $\hat{x}_{max}$ as a horizontal line. The value of $k_o$ at which the $x_{min}(k_o)$ curve intersects the $\hat{x}_{max}$ line is $\tilde{k}_o$.

In Figure S30, $\tilde{k}_o = 11.43$. If $k_o > \tilde{k}_o$, then the $\hat{x}_{max}$ horizontal line is above $x_{min}$ curve. To illustrate, say $k_o$ is 16. The $x_{min}$ for this $k_o$ is 1.814 whereas $\hat{x}_{max}$ is 2. Therefore, all strains supplying an $x < 1.814$ cannot increase when rare and are excluded.

Alternatively, if $k_o < \tilde{k}_o$, say $k_o = 8$, then the $\hat{x}_{max}$ horizontal line at 2 is below the $x_{min}$ curve. Hence no strain with $k_o = 8$ can increase when rare because $\hat{x}_{max} = 2$ is the maximum altruism it can supply whereas for this $k_o$, an $x_{min} = 2.25$ would be needed to increase when rare.

Assuming $k_o > \tilde{k}_o$ so that the microbe can enter the holobiont, what degree of altruism, $x$, should the microbe provide to the host? For a given $k_o$, $x$ must be greater than $x_{min}(k_o)$ to enter. The $r(x)|_{z=\hat{z}(x)}$ is an increasing function of $x$, so the more altruism the better from the standpoint of the speed at which the microbe can increase when rare. However, $x$ should not exceed $x_{max}$ because at that value of altruism the host ceases making antibodies and any altruism beyond that value incurs cost with not added benefit. So, if $k_o > \tilde{k}_o$, does the holobiont population dynamics and evolution culminate at $x = x_{max}$? No, because the dynamics are complex and, as noted before, the criterion for increase when rare is necessary but not sufficient for coexistence and provides only a limited guide to the final outcome.

Specifically, consider three examples. In the first, strain-1 has $x = x_{max}$ and strain-2 has $x = x_{min}(k_o)$. As Figure S31 shows, the maximally altruistic strain excludes the minimally altruistic strain.

Next, let strain-2 provide a bit more altruism than the minimal, say 2.5% more. In this case, Figure S32 shows that the maximally altruistic strain comes to a polymorphism with the minimal plus 2.5% strain.

Finally, let strain-2 provide still a bit more altruism than the minimal, say 5% more. Here, Figure S33 shows that the maximally altruistic strain is shed by the minimal plus 5% strain.

Examining Figure S33 reveals what is going on. The figure shows that the maximally altruistic strain initially increases faster than the minimal plus 5% strain in accordance with its having a higher $r(x)|_{z=\hat{z}(x)}$. But as the dynamics proceed, the minimal plus 5% strain increases enough to colonize sufficient two-strain holobionts from which it then excludes strain-1, leaving the maximally altruistic strain with barely enough single-strain holobionts to replace itself. Because strain-1 can still increase when rare, it is shed, not excluded, from the holobiont population—its absolute abundance slowly increases but its percentage of the microbiome approaches zero.

Thus, the holobiont population winds up with microbes that have a bit more altruism than the minimum necessary along with hosts that supply a corresponding antibody response. Nonetheless, this arguably imperfect outcome, which is less than the best possible for both parties, does represent host-microbiome integration. Through its antibody response to the altruism offered by various microbial strains, the host determines which microbes can enter its microbiome. Holobiont assembly based on host-orchestrated species sorting (HOSS) of the microbes, rather than coevolution, is the likely cause of host-microbiome integration. A diagram illustrating the process of holobiont assembly appears in Figure 4 of the main article. The top left insert in Figure 4 is adapted from Figure S30.

**Code Repository.** *Mathematica* notebooks to generate the figures in this Supplementary Material are available for download at: https://doi.org/10.5061/dryad.fqz612jw7

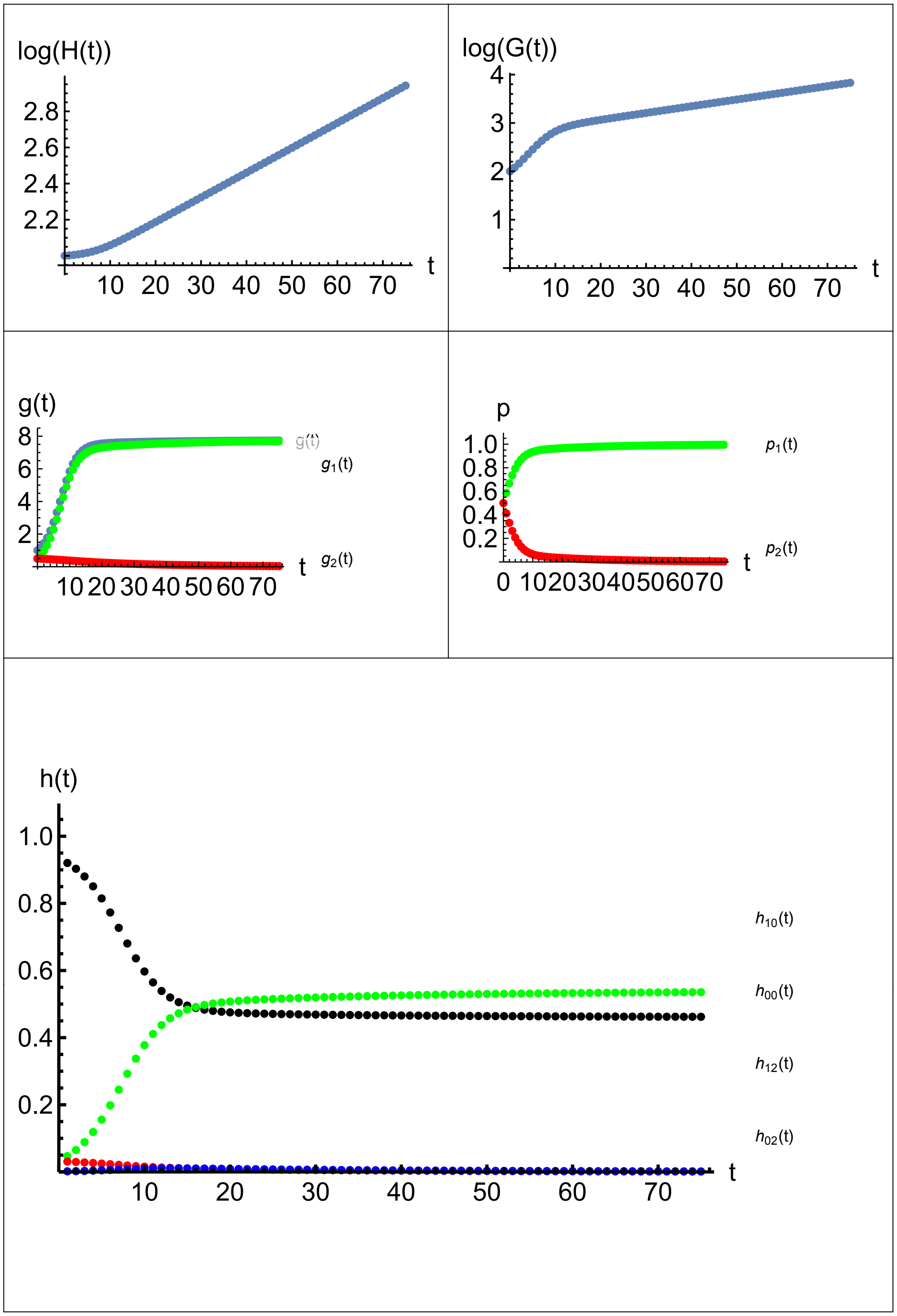


Figure S31: *Minimally altruistic strain-2 excluded by maximally altruistic strain-1. Here,* $x_1 = \hat{x}_{max}$, $x_2 = x_{min}(k_o)$, $k_o = 16$, $x_{min}(k_o) = 1.814$, $\hat{x}_{max} = 2$, $\alpha_o = 3$, $\mu = 1$, $\beta = 1$, $\gamma = 1$, $d_o = 0.1$, $\nu = 0.2$, $b = 0.03$, $c = 1$, $W_o = 1$, $H_{init} = 100$, $G_{1,init} = 50$, $G_{2,init} = 50$, $t_{max} = 75$.

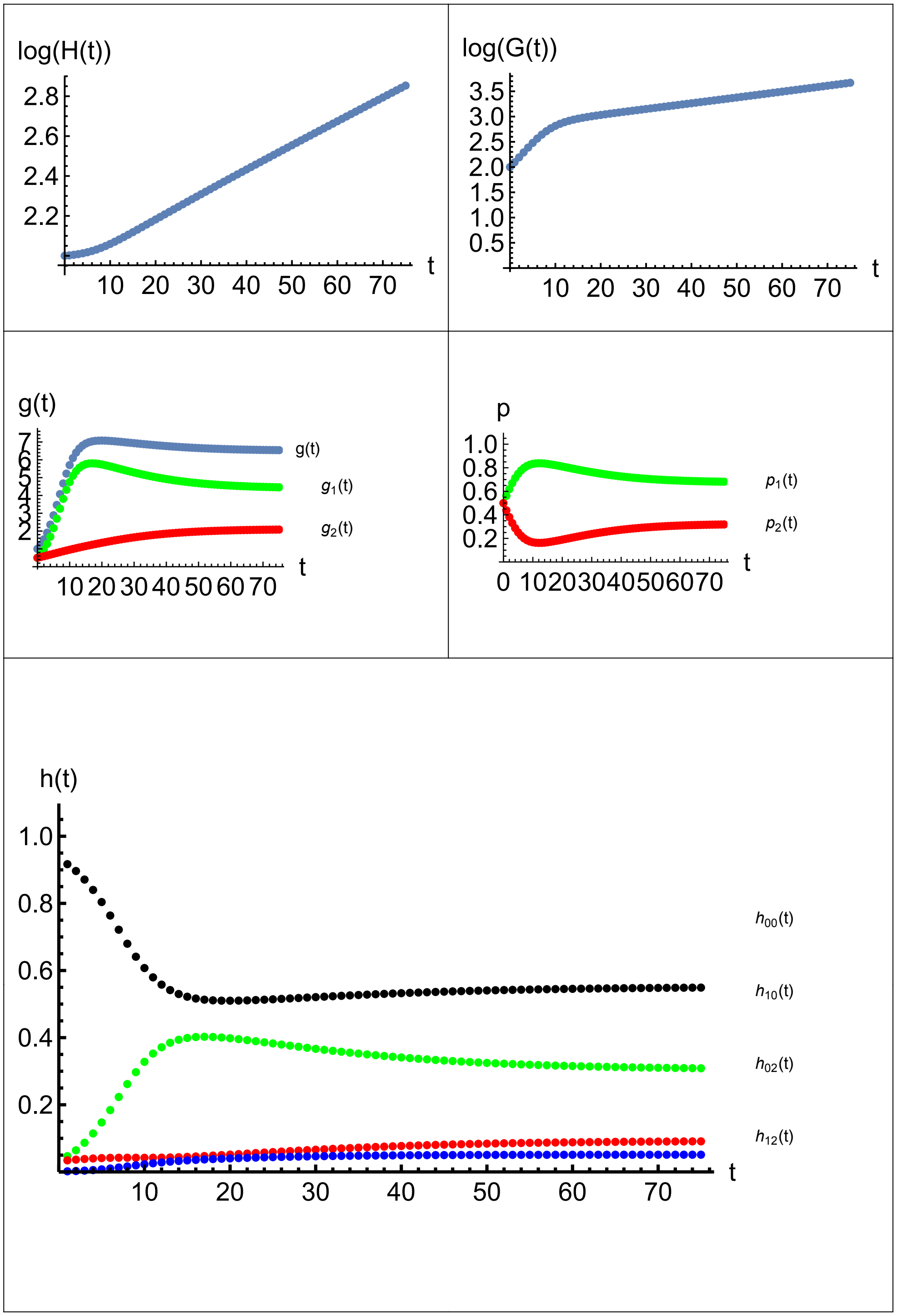


Figure S32: *Minimally altruistic plus 2.5% strain-2 coexists with maximally altruistic strain-1. Here,* $x_1 = \hat{x}_{max}$, $x_2 = 1.025 \times x_{min}(k_o)$, $k_o = 16$, $x_{min}(k_o) = 1.814$, $\hat{x}_{max} = 2$, $\alpha_o = 3$, $\mu = 1$, $\beta = 1$, $\gamma = 1$, $d_o = 0.1$, $\nu = 0.2$, $b = 0.03$, $c = 1$, $W_o = 1$, $H_{init} = 100$, $G_{1,init} = 50$, $G_{2,init} = 50$, $t_{max} = 75$.

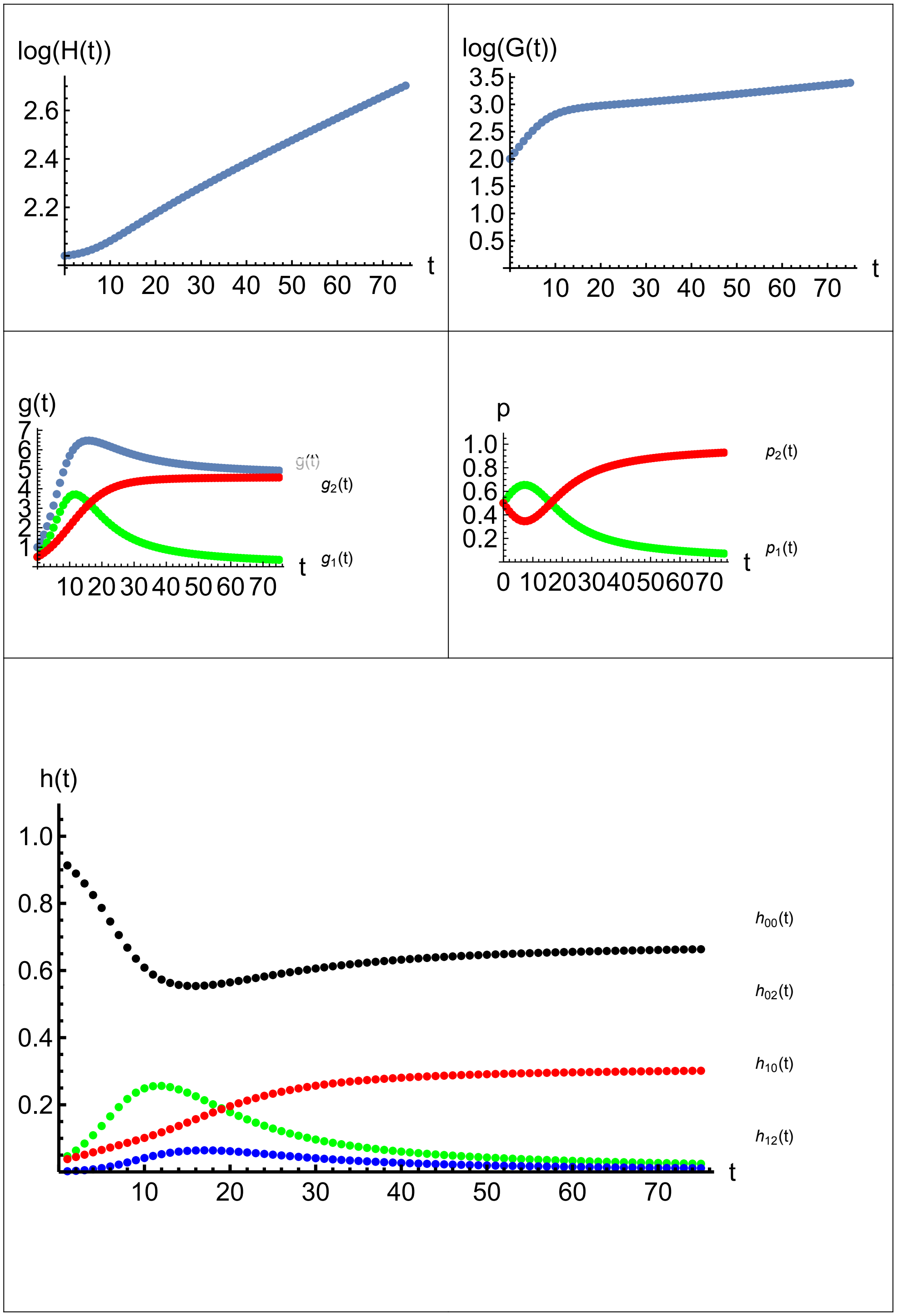


Figure S33: *Minimally altruistic plus 5% strain-2 sheds the maximally altruistic strain-1. Here,* $x_1 = \hat{x}_{max}$, $x_2 = 1.050 \times x_{min}(k_o)$, $k_o = 16$, $x_{min}(k_o) = 1.814$, $\hat{x}_{max} = 2$, $\alpha_o = 3$, $\mu = 1$, $\beta = 1$, $\gamma = 1$, $d_o = 0.1$, $\nu = 0.2$, $b = 0.03$, $c = 1$, $W_o = 1$, $H_{init} = 100$, $G_{1,init} = 50$, $G_{2,init} = 50$, $t_{max} = 75$.
